# Non-coding RNAs control metabolic state in mycobacteria

**DOI:** 10.1101/660928

**Authors:** Dolly Mehta, K Anjali, A Achuthan, Ritu Gupta, Arati Ramesh

**Affiliations:** National Centre for Biological Sciences, GKVK Campus, Bellary Road, Bangalore, India 560065.; SASTRA UNIVERSITY, Tirumalaisamudram, Thanjavur - 613401; Institute for Stem Cell biology and Regenerative Medicine, NCBS-TIFR campus, GKVK Campus, Bellary Road, Bangalore 560065.

## Abstract

Non-coding RNAs play pivotal roles in bacterial signaling. However, RNAs from certain phyla (specially high-GC actinobacteria) remain elusive. Here, by revamping existing approaches we discover a family of structurally conserved RNAs in actinobacteria. These RNAs function by the recruiting ANTAR proteins to select transcripts; regulating them via translational repression. By overlapping with ORF start sites, these RNAs provide mechanisms by which even leader-less transcripts are regulated. In mycobacteria, transcripts marked by ANTAR-target RNAs are few but encode important redox enzymes especially involved in lipid metabolism. Notably, the cellular response to ANTAR-regulation is hierarchical, wherein immediate metabolic changes induced by ANTAR-RNA binding are amplified through a global transcriptomic response. This includes several genes from oxidative/reductive pathways; ultimately switching cells towards reductive metabolism. This discovery of ANTAR-target RNAs and associated regulation places RNAs as crucial players in controlling metabolic flexibility of mycobacteria, proposing a prominent role for ANTAR regulation across actinobacteria.

## INTRODUCTION

The diverse functions of RNAs in the cell, stem from their ability to form complex folds and structures. Through these structures, RNAs create the chemical complexity and geometry required to selectively recognize their interacting partners, which range from proteins to other RNAs and even small molecules. Structures within coding and non-coding regions of mRNA have specially been in the spotlight as signal-sensing and sometimes signal-integrating RNA hubs that can directly influence core processes like translation, transcription and RNA degradation.

Natural RNA sensors such as riboswitches that reside in the 5’ untranslated regions (UTRs) of mRNAs, alter their structure in response to a ligand(1)(2). This altered structure may sequester a transcriptional terminator or ribosome-binding site to control transcription elongation or translation initiation respectively. Ribozymes such as the glmS RNA self-cleave upon metabolite binding(3). The newly formed RNA terminus recruits a specific ribonuclease that degrades the transcript. In a distinct example of signal integration mediated by RNAs, the mRNA transcript encoding ethanolamine catabolic enzymes in *Enterococcus, Listeria* and *Clostridium*, is regulated by both the enzyme’s cofactor (B12) and its substrate (ethanolamine)(4). The UTR for this transcript harbors both a B12-responsive riboswitch as well as a small structured RNA motif that recruits an ethanolamine-responsive RNA-binding protein; thus integrating cofactor and substrate sensing through the same UTR. Given the extent of cross talk between cellular metabolites across metabolic pathways, such mechanisms of RNA-mediated signal integration may be more widely prevalent in biology, than have yet been appreciated.

A recent study identified a conserved family of dual stemloop RNAs (called ANTAR-target RNAs) within 5’ UTRs of select mRNAs across several species of firmicutes and proteobacteria(4). This class of RNA recruits an RNA-binding domain called ANTAR. ANTAR is widely present across all bacteria and typically interacts with RNA to stabilize an anti-terminator conformation of the UTR. This allows read-through, of an otherwise termination prone transcript. The majority of ANTAR proteins are part of a signal-responsive phosphocascade wherein the ANTAR protein gets phosphorylated by its cognate histidine kinase in the presence of an activating small molecule signal. This phosphorylation then activates ANTAR for RNA binding. These studies positioned ANTAR-target RNAs as important signaling nodes in bacterial phyla such as firmicutes and proteobacteria, where ANTAR-target RNAs were computationally identified and experimentally validated(5)(6)(7). Previous attempts to identify ANTAR-target RNAs used standard covariance based computational approaches that use conservation of base-pairing potential (covariance) as an indication of structural conservation among RNAs. Although ANTAR proteins themselves are ubiquitously present across all bacteria, these covariance based searches failed to identify RNAs in certain bacterial phyla-such as those characterized by unusual nucleotide propensities(4). Notable among these are the high GC genomes of actinobacteria, which includes human pathogens such as *Mycobacteria*, plant pathogens such as *Rhodococcus* and the largest class of antibiotic producers like *Streptomyces*, among others.

Separately, RNA-seq data from mycobacteria under stationary and exponential phase growth has catalogued some small RNAs that are present in the cell (8)(9), but their roles and functions still remain a mystery(10). In several firmicutes and proteobacteria, the small RNA chaperone - Hfq has proved to be excellent bait for identifying new (chaperone-bound) small RNAs(11)(12). However, since actinobacteria lack the more common RNA chaperones such as Hfq, this rules out such approaches as a means to fish out small RNAs. Although ANTAR-target RNAs have not been identified previously in actinobacteria, we observed a disproportionately large number of ANTAR proteins per actinobacterial genome (Figure 1); arguing for not only the presence of ANTAR-target RNAs but also a significant role for these RNA protein complexes across actinobacteria.

**Fig 1.**
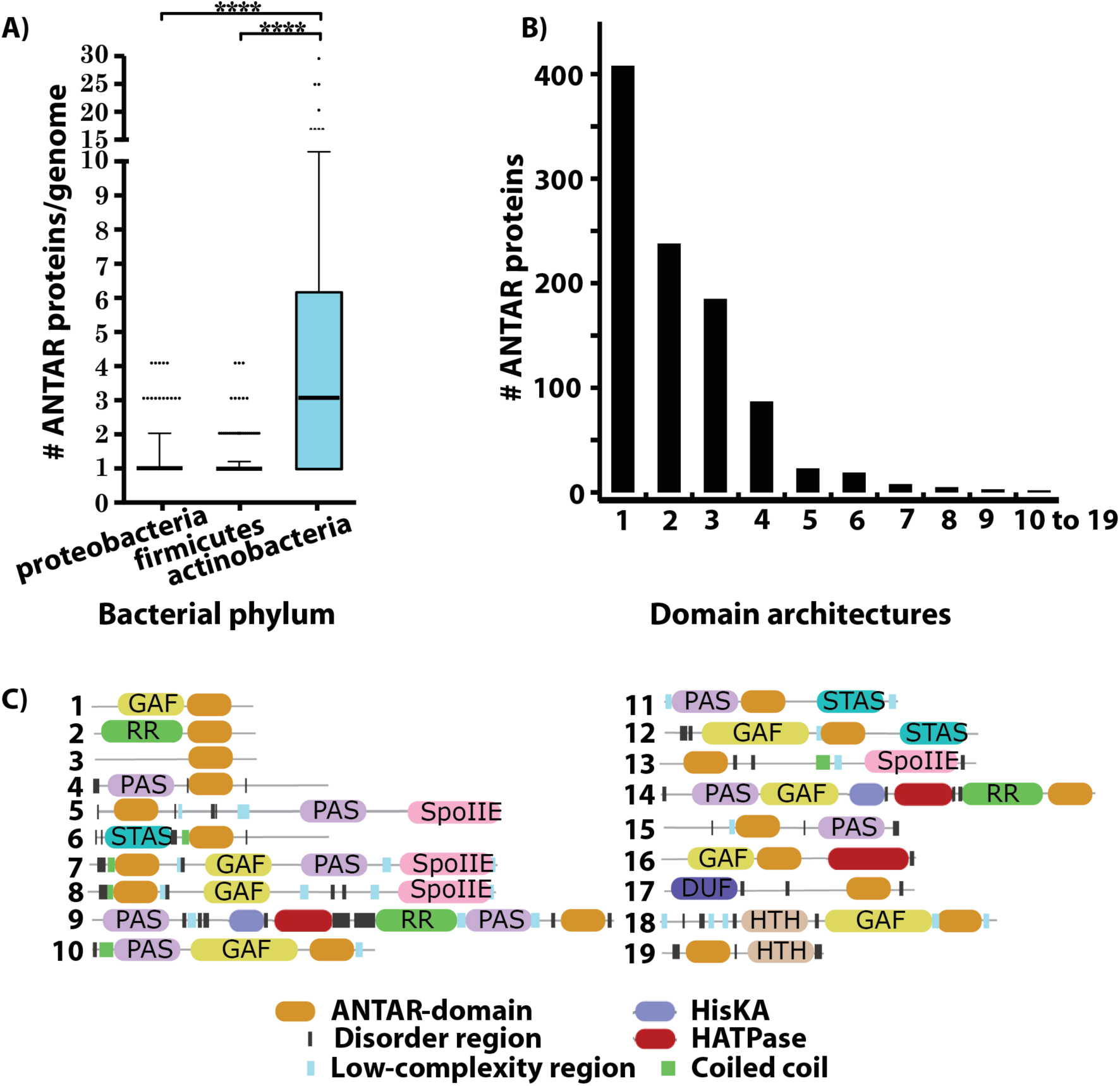
ANTAR proteins are significantly over-represented in GC rich actinobacteria. The family of ANTAR domain containing proteins as listed in the protein family database (Pfam ID: PF03861) was analyzed. A) Plot shows the distribution of ANTAR domain containing proteins per genome across major bacterial phyla. Median (black line), 10-90 percentiles (whiskers) and outliers (black circles) are shown. Statistical significance (p-value <0.0001 shown as asterisks) was calculated using one-way ANOVA. Actinobacterial genomes possess an average of three ANTAR proteins per genome when compared to proteobacteria and firmicutes. B) Bar plot showing number of mycobacterial ANTAR proteins listed in pfam for each type of domain architecture. 19 distinct domain architectures are present for ANTAR proteins in mycobacteria. C) Details of 19 domain architectures from panel B, are shown. A majority of ANTAR domains co-occur with the ubiquitous GAF(17) signal sensing domain (domain architecture 1 of panel B) or are part of response regulators (domain architecture 2 of panel B). Different domains are indicated in different colors as shown in the index.

In this current study, we use ANTAR proteins as a handle for discovering non-coding RNAs in actinobacteria, and uncover the function of these ANTAR-RNA complexes in the cell. Using an integrative approach of computational RNA search, in vitro biochemistry, gene expression profiling and in vivo phenotypic analyses, we have discovered a structurally conserved family of small non-coding RNAs, present ubiquitously across actinobacteria. We use mycobacteria as a relevant representative of actinobacteria and find that the identified RNA candidates recruit the mycobacterial ANTAR protein (Rv1626), in an ANTAR phosphorylation dependent manner. ANTAR-RNA binding results in translational repression of select genes encoding lipid metabolism and redox-dependent enzymes. Further, the final cellular response to ANTAR mediated regulation extends beyond the direct response to binding. This manifests as a global transcriptomic change involving several redox-dependent enzymes, stress response proteins and oxidative/reductive processes. We propose a mechanism by which ANTAR induces a hierarchical gene expression response, ultimately switching cells towards a more reduced metabolic state. Our collective findings reveal an ANTAR protein and RNA mediated regulatory network that controls the metabolic flexibility of mycobacterial cells.

## RESULTS

### ANTAR proteins are significantly over-represented in actinobacteria

Previous studies have described ANTAR as a highly conserved, RNA-binding effector domain, that modulates transcription(4)(6, 13)(14). This transcriptional control comes via ANTAR’s recognition of a dual stemloop RNA motif that resides in the 5’ UTRs of select mRNA transcripts and whose secondary structure is conserved, with little sequence conservation. Notably however, the particular signals that activate the cascade leading to ANTAR-RNA recognition and transcriptional regulation, are diverse across different bacteria. For example, AmiR in *Pseudomonas sp*. responds to amides(15), EutV in *Enterococcus* responds to ethanolamine(5)(4), NasR in *Klebsiella* responds to nitrates(13) etc. This suggests that different bacteria may have co-opted ANTAR as an effector domain to control completely different cellular pathways.

We made three intriguing observations-1) Although ubiquitous in bacteria, the protein family-pfam(16) database reports a disproportionate and significantly higher number (p<0.0001) of ANTAR proteins in the GC rich class of actinobacteria (Figure 1A), 2) previous studies centered around searching for ANTAR-target RNAs in firmicutes (low GC bacteria) in fact failed to identify any RNA targets in actinobacteria(4), and 3) the domain architectures of ANTAR proteins in actinobacteria are distinct from those in other bacteria, with a bias towards GAF(17) and PAS(18) signaling domains typically occurring in tandem with ANTAR (Figure 1B-C, S1). These observations raise the possibility that actinobacteria might use ANTAR based signaling for distinct pathways not known to be linked to ANTAR. Given that we have no idea about the distribution of ANTAR-target RNAs in actinobacteria, we hypothesized that ANTAR might provide a handle towards a molecular level understanding of small RNA based regulation which, in actinobacteria, has not been explored comprehensively thus far. Here we use mycobacteria as a relevant representative of actinobacteria to identify ANTAR-target RNAs and understand their function in the cell.

### The ANTAR protein in mycobacteria is phosphorylated by a cognate histidine kinase

Our primary inspiration to look for ANTAR-target RNAs in actinobacteria stems from our observations of the ANTAR proteins. Hence we first characterized these proteins. Analysis of the *Mycobacterium tuberculosis* genome revealed two ANTAR encoding genes. One of these, called Rv1626, has a typical phosphoreceiver domain found in bacterial response regulators. This indicates the possibility of phosphorylation by a histidine kinase. We used an *in vitro* phosphorylation assay with P^32^-γATP as the phosphodonor, to measure the degree of phosphorylation of recombinantly expressed and purified histidine kinase and ANTAR proteins. This assay confirmed that Mtb ANTAR protein (Rv1626) gets phosphorylated *in vitro* by its cognate histidine kinase called Rv3220c (Figure 2A), as also reported previously (19, 20). Unlike the majority of bacterial histidine kinases, we find that autophosphorylation of Rv3220c and subsequent phoshotransfer to Rv1626 doesn’t require any additional small molecule as signal. Instead this phosphorylation is seen simply upon addition of physiologically relevant amounts of magnesium (2mM) and ATP (1mM). To investigate if signal independent phosphorylation is governed by the kinase or the ANTAR protein, we measured phosphorylation of Rv1626 by a non-cognate histidine kinase EutW (Figure 2B). EutW from *E. faecalis* is known to autophosphorylate only in presence of its signal-ethanolamine, and subsequently phosphorylate the *E. faecalis* ANTAR protein EutV(4). When the *in vitro* phosphorylation assay was carried out using Rv1626 and EutW, we observed that the signal dependence is restored (Figure 2B). Collectively, these experiments raise the possibility that in *mycobacteria*, this two-component kinase (Rv3220c) and ANTAR (Rv1626) system would be in a constitutively active, phosphorylated state (Schematic in Figure 2D). Regulation might therefore come from either controlling the level of the transcripts/protein of the kinase or the ANTAR protein, or from a signal that inhibits phosphorylation.

**Fig 2.**
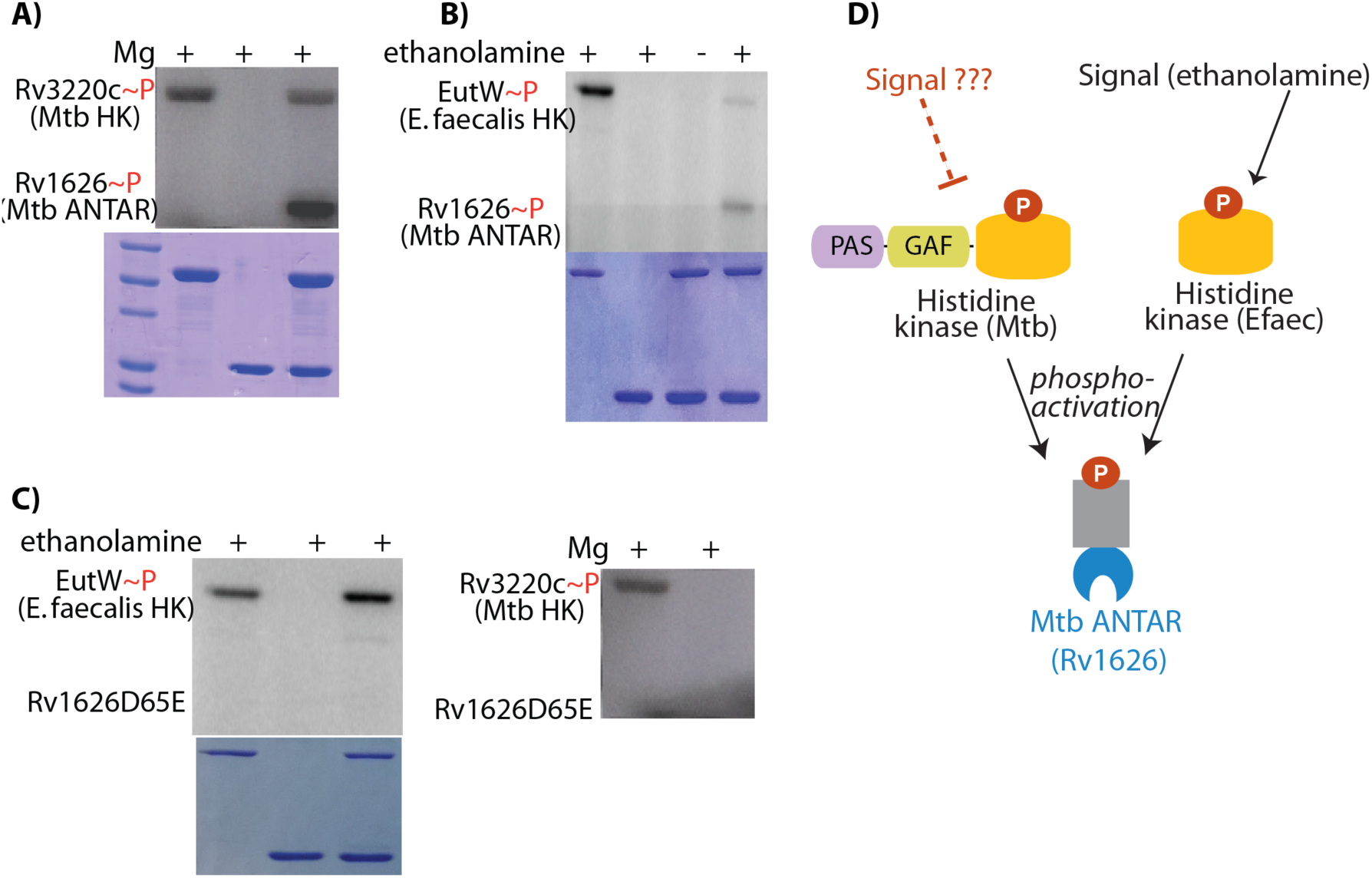
The mycobacterial ANTAR protein is part of a signaling phosphocascade. In vitro phosphorylation assay was performed using P^32^-ATP to follow phosphorylation of recombinantly purified histidine kinase and response regulator. Proteins were seprated using electrophoresis and visualized with autoradiography (top) and coomassie staining (bottom). A) A cognate histidine kinase Rv3220c from Mtb gets autophosphorylated in the presence of 2mM Mg, 1mM ATP. In the absence of Rv3220c, the Mtb ANTAR protein Rv1626 shows no phosphorylation. Phosphotransfer to Rv1626 is seen only in the presence of Rv3220c. B) Rv1626 is phosphorylated by *E. faecalis* histidine kinase called EutW when the reaction is performed in presence of the activating small molecule ethanolamine. C) Rv1626 protein with a D65E mutation does not get phosphorylated by either *E. faecalis* kinase (left) or the cognate mycobacterial kinase (right). D) Schematic showing ANTAR phosphorylation by cognate and non-cognate histidine kinases. Phosphorylation of Mtb kinase Rv3220c, even in the absence of activating small molecules raises the possibility of a signal that inhibits phosphorylation.

**Fig 3.**
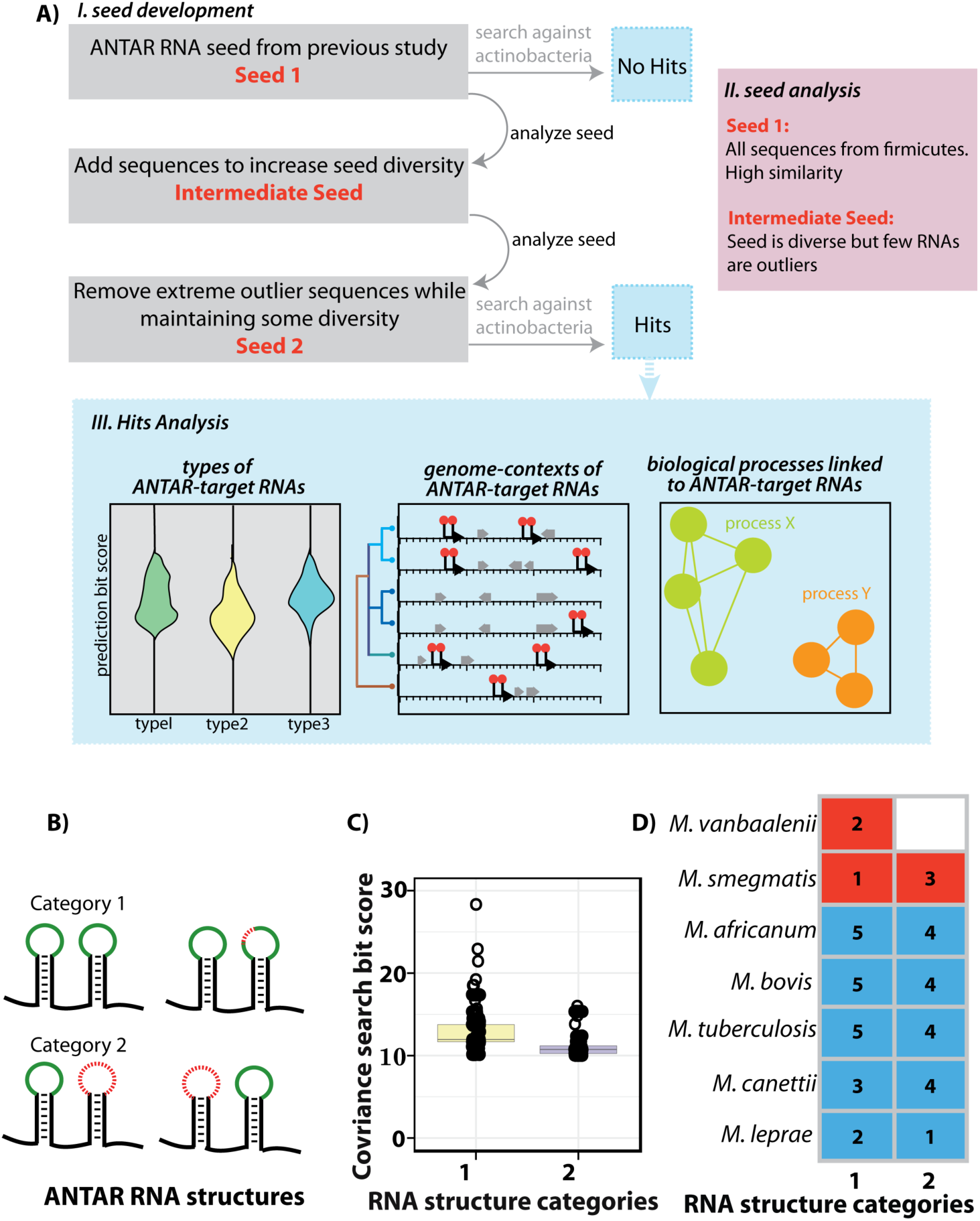
Covariance search identifies ANTAR-target RNAs in mycobacteria. Computational search for RNAs performed using Infernal v1.0.2. A) Schematic shows steps in identifying RNAs-Seed development, Seed analysis and Hits analysis. Seed 1 from previous studies yielded no RNA hits in actinobacteria. Analysis of the seed showed lack of sequence diversity. This was corrected in the Intermediate Seed by including diverse ANTAR-target RNAs from firmicutes/ proteobacteria. Analysis of this seed identified outliers that deviated significantly from required consensus. These outliers were excluded to create Seed 2, which now yielded hits in actinobacteria. Analysis of hits (blue box) identifies structural categories, genome contexts and biological processes linked to the RNA hits. B) Cartoon showing Category1 (identical or single mismatch from consensus) and Category 2 (one stem deviates from consensus) ANTAR-target RNA structures found in mycobacteria. Red regions indicate deviation from consensus. C) Box plot shows the distribution of covariance search bit scores for RNA structure categories, with median (black line). Category1 RNAs show higher bit scores compared to Category 2. D) Distribution of RNAs in select pathogenic (blue) and saprophytic (red) mycobacteria. The number of RNAs is marked within each box.

A multiple sequence alignment of Rv1626 with other ANTAR containing response regulators predicted a single aspartate (D65 in Rv1626) as the likely site of phosphorylation (Figure S2). We substituted D65 with a glutamate residue, and find that substituting aspartate with a glutamate at this single site (D65E) abolishes phosphorylation by Rv3220c (Figure 2C). Further, this mutation functions as a phosphoactive mimic (Figure 4). This allowed us to use Rv1626 D65E protein as a signal-independent version of ANTAR∼P in several experiments described below.

**Fig 4.**
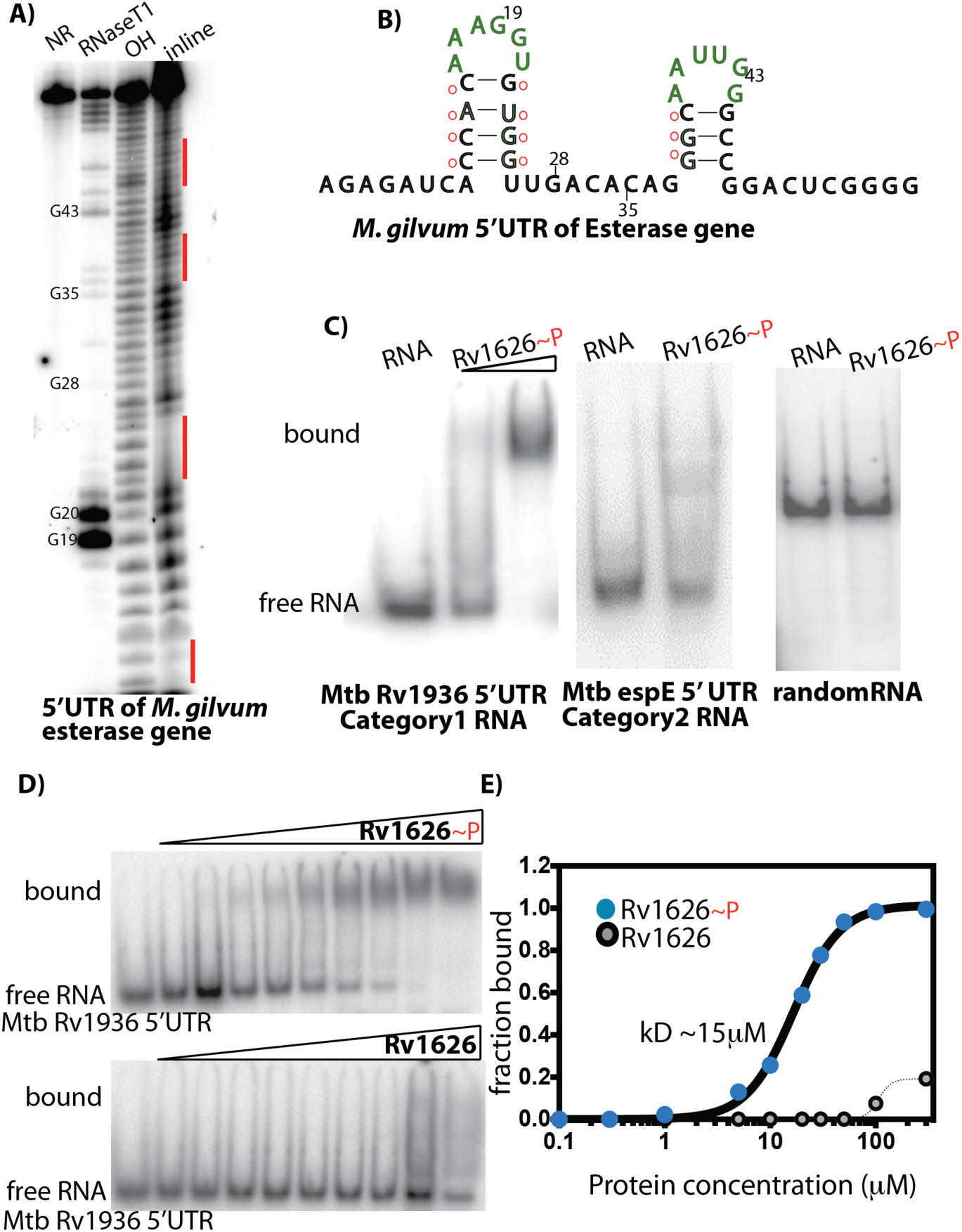
Bioinformatically predicted RNAs form a dual stemloop structure and recruit ANTAR∼P specifically. A) RNA secondary structure analysis of a predicted category 1 RNA (5’ UTR of *M. gilvum* Esterase gene) using in-line probing assay. 5’-P^32^labeled RNA was incubated in alkaline buffer for 38 hours, allowing spontaneous backbone cleavage. RNA cleavage products (inline) were separated on a 10% denaturing Urea PAGE and visualized using autoradiography, along side controls (NR-unreacted, OH-alkaline cleavage at every nucleotide position and RNaseT1-cleavage at G positions). Structured regions appear as gaps (marked in red) whereas flexible regions of the RNA appear as bands. B) Secondary structure of 5’ UTR of *M. gilvum* Esterase gene based on panel A shows a dual stem loop structure where base paired regions (red dots) appear protected in the in line probing assay. C) Electrophoretic mobility shift assay (EMSA) to study interaction of different predicted RNAs with respective ANTAR proteins (as indicated in figure). 5’-P^32^labeled RNA was incubated with recombinantly expressed and purified protein and sample was resolved on a 5% non-denaturing gel. RNA-protein complex showed retarded mobility as compared to free RNA. Candidates from both RNA category1 and 2 showed binding to Rv1626∼P (Rv1626D65E) protein. D) Category 1 RNA (5’UTR of Mtb Rv1936) was titrated with Rv1626∼P (Rv1626D65E) or Rv1626 protein and subjected to EMSA. RNA binding affinity for Rv1626∼P (Rv1626D65E) was notably higher than that of Rv1626 protein. E) EMSAs from panel D were quantified and a binding curve showing fraction RNA bound versus protein concentration was plotted. Rv1626∼P binds RNA with a kD of ∼15uM as compared to Rv1626 where kD could not be determined.

### Identifying a family of ANTAR RNAs across mycobacteria

Previous work on ANTAR in *Enterococcus faecalis* had identified a dual stemloop RNA motif that selectively recruits the ANTAR protein-EutV. The motif was characterized by two short stems separated by a linker, each containing a hexanucleotide loop with positions A1 and G4 of the loop showing sequence conservation(4) (Figure 3C). To see if similar RNAs may be present in other bacteria, a bioinformatic approach had been used. In this approach, an alignment is first created with known RNA sequences that define the search model (seed) and then genome sequence databases are scanned for similar RNAs, scoring on the basis of a combination of sequence and structural consensus(21). While that search identified ∼400 RNAs in diverse bacterial phyla, it notably failed to find more than a couple of RNAs in actinobacteria.

To find ANTAR-target RNA motifs in actinobacteria we modified the above approach (Figure 3A). First we took the 17 sequences (Seed 1) used as a search model in the previous study and analyzed them using cmbuild(21), a program that builds a statistical profile of alignments. We found that these 17 sequences clustered together based on high (54% sequence identity and a CM score of 0.75) sequence similarity (Figure 3A-B). Since all these sequences were from three firmicute species-*Enterococcus, Listeria* and *Clostridium*, we reasoned that highly similar sequences may bias the search against finding hits in actinobacteria. To remove this bias we enriched the seed in diversity, even while maintaining the core RNA structure. For this we created an intermediate seed which consisted of 39 ANTAR-target RNA sequences found in different bacteria (reported in the previous study) and retained 12 sequences from the original Seed 1. The resulting 51 sequences were once again subjected to cmbuild analysis. We found that 47 of these sequences clustered together (with ∼51% sequence identity), marking 4 as outliers that show too much diversity. We now used the 47 sequences as seed (Figure 3B-C, TableS1) and searched against all 128 sequenced mycobacterial genomes. This search now predicted nearly 220 ANTAR-target RNA candidates across mycobacteria (Figure 3D-E, TableS2, TableS3) and over ∼230 RNAs in other actinobacteria such as *Rhodococcus* and *Streptomyces*.

Further analyzing these putative hits from mycobacteria, RNAs that completely match the dual stemloop consensus of the search model or have a single nucleotide substitution in either one of the loops were assigned to category 1. RNAs where one of the two stemloops deviates from the seed consensus were classified into category 2 (Figure 3D, S3). RNA hits were not restricted to any particular species and were obtained in both the pathogenic and saprophytic mycobacteria, typically mapping to the upstream regions of ORFs (Figure 3E). Transcription start site mapping performed on *M. tuberculosis* and *M. bovis* transcripts have identified +1 transcription start sites that map upstream of some of our RNA candidates(22)(23)(24), indicating that these RNAs are indeed present in the cell.

### Phosphorylated ANTAR protein binds predicted RNAs with specificity

With the list of ANTAR-target RNA candidates identified via bioinformatics, we first tested if these RNAs in fact fold into the predicted structures *in vitro*. For this we used the in-line probing assay where 5’-P^32^-labeled RNA is incubated in alkaline buffer and allowed to undergo spontaneous backbone cleavage(25). When analyzed on a denaturing gel, flexible (cleaved) versus structured/base paired (protected) regions of the RNA can be identified with single-nucleotide resolution. In-line probing confirmed that candidate ANTAR-target RNAs from mycobacteria adopt secondary structures similar to that predicted by the covariance analysis. A representative in-line probing assay perfomed on the 5’ UTR of the *M. gilvum* esterase transcript shows that regions encompassing the two stems appear protected whereas the connecting linker region and the hexanucleotide loops show cleavage (Figure 4A-B). Next, using electrophoretic mobilty shift assays (EMSAs), we tested the binding of ANTAR protein (Rv1626) to several RNAs belonging to the two categories of dual stemloop RNAs, predicted in different species of *mycobacteria*. We find that the RNAs are recognized *in vitro* by Rv1626, whereas a structured riboswitch RNA used as a negative control shows no binding (Figure 4C and Figure S4A). This confirmed that our approach indeed identifies RNAs targeted by ANTAR and that these RNAs are widely distributed among different mycobacterial species. In all cases, ANTAR binds RNAs only in its phosphorylated state. For example, Rv1626∼P binds an *M. tuberculosis* Category 1 RNA target (5’UTR of Rv1936), with an affinity of ∼15μM whereas the binding affinity for RNA could not be measured for the unphosphorylated protein (Figure 4D-E). RNA binding by the phosphoactive Rv1626D65E mutant was comparable to that of Rv3220c-activated Rv1626, showing that the D65E mutant protein indeed functions as a phosphoactive mimic (Figure S4B). The binding affinity seen for Rv1626∼P is weaker than that observed previously (1μM) for the *E. faecalis* protein EutV∼P(4). It is possible that these affinites are fine-tuned to reflect the cellular concentration of ANTAR-target RNAs in different bacteria. Collectively, we find that these ANTAR-RNA interactions are highly specific for the activated form of the ANTAR protein, indicating a role for these RNAs in cellular signaling.

Under *in vitro* conditions mycobacterial candidate RNAs also show binding to the *Enterococcus faecalis* ANTAR protein-EutV. Similarly, the eutP 5’ UTR from *E. faecalis* (a known ANTAR-target RNA) binds P∼Rv1626 (Figure S4C). This cross reactivity argues for conserved mechanisms of protein-RNA recognition, despite the sequence level differences that we find in ANTAR-target RNAs from different bacteria.

### ANTAR-target RNAs typically reside upstream of lipid metabolism and redox enzymes

We analyzed the genomic contexts of all the bioinformatically identified ANTAR-target RNAs, several of which we had experimentally validated (Figure 5A). This analysis shows that a striking majority of these are found upstream of genes involved in lipid metabolism and related redox-dependent enzymes. These include genes annotated as lipases, esterases, epoxide hydrolases, short-chain dehydrogenases and reductases and their distribution is spread across all mycobacterial species (Figure 5A, Figure S5A and Table S2).

**Fig 5.**
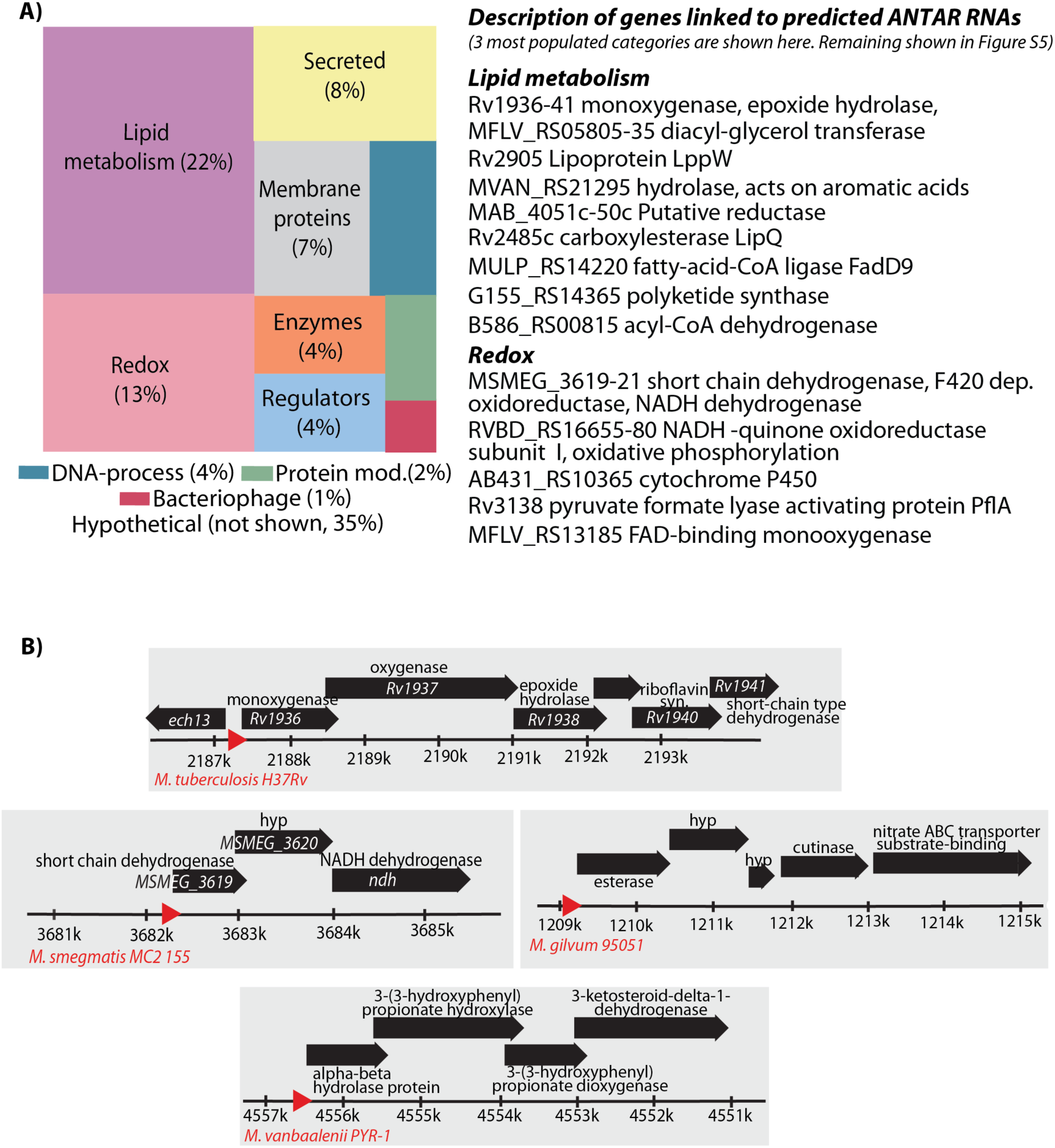
Computationally predicted ANTAR-target RNAs are linked to lipid metabolism and redox-dependent enzymes. Genome contexts of predicted ANTAR-target RNAs were analyzed and categorized on the basis of NCBI gene annotations coupled with manual curation. A) Treemap (left) shows the cellular processes/functions of genes downstream of predicted ANTAR-target RNA candidates. RNA hits from all species of mycobacteria were included in the analysis. Identical RNAs from different strains belonging to the same species were not included. Distribution of genes in each category is shown (%). Descriptions of the genes linked to ANTAR-target RNAs are shown (right). Processes involving lipid metabolism and redox are highly represented. B) Representative genomic contexts of predicted ANTAR-target RNAs (red arrow) are shown with their downstream ORFs (black bars) and their operonic arrangement. Mycobacterial species names are indicated (red). Gene annotations are marked above the ORF. *hyp* indicates hypothetical protein.

In *M. tuberculosis* and ∼50 strains of the MTB complex, we found a top scoring RNA candidate upstream of melF (Figure 5B, Table S2), the first gene of the *mel2* operon (*mycobacterial enhanced infection loci-* Rv1936 to 1941). This operon contains two flavin-dependent oxygenases, an Fe-S dehydrogenase, an epoxide hydrolase and short-chain type reductases for lipid metabolism. While the specific substrates for these enzymes are not yet known, deletion of this operon in *M. tuberculosis* results in accummulation of cell wall virulence lipids(26) (triacylglycerides and DGIM), in response to propionate or oxidative stress; suggesting that repression of the mel2 operon may be important for maintaining lipid homeostasis during stress. Accordingly, the mel2 operon is implicated in mycobacterial resistance to both reactive oxygen and nitrogen species(27). Another bioinformatically predicted RNA from *M tuberculosis* that we validated experimentally, resides upstream of espE (*ESX-1 secretion associated protein E*), the first gene of the ESX-1 operon in *M. tuberculosis*(28). This RNA is conserved among the strains of the MTB complex (Figure 5B). ESX-1 is a secretory system responsible for virulence of Mtb, where ESX-1 associated effector proteins are secreted upon infection and interfere with the host immune response(29). Interestingly, the operon containing EspE is transcriptionally controlled by WhiB6, a redox-responsive regulator(30). However, constitutive expression of WhiB6 under rich media conditions does not result in EspE locus upregulation and a possible mechanism of post-transcriptional regulation has been proposed for this locus. Our results suggest that this post-transcriptional regulation may occur via ANTAR. Based on these observations we speculate that ANTAR-target RNAs in *M. tuberculosis* and related pathogens are likely to be overproduced upon infection of host.

In *M. vaccae* and *M. vanbalenii* both saprophytic *mycobacteria*, we find a similar link between ANTAR and lipid catabolism (Figure 5B). We find candidate RNAs upstream of an operon which in *E.coli* and *Rhodococcus* is implicated in the break down of aromatic acids (hydroxy-phenyl propanoid compounds). These molecules are widely available in nature and several soil-dwelling bacteria metabolize them as a sole carbon source(31).

ANTAR’s link with lipid metabolism and redox is further illustrated in *M. smegmatis.* Here we predict an ANTAR-target RNA motif directly upstream of an operon (MSMEG_3619 to 3621) that encodes a short-chain dehydrogenase, a putative luciferase-like oxidoreductase and *ndh-* a non-proton pumping NADH dehydrogenase(32) (Figure 5B). MSMEG_3619 is an NADH-dependent reductase and mutations in *ndh* are shown to increase the NADH/NAD+ ratio in the *M. smegmatis*(33), suggesting that this operon may directly influence cellular redox via control of reductive metabolites.

Taken together, our analysis of the genomic contexts of ANTAR-target RNAs show that although the particular genes/operons regulated by ANTAR may be distinct in different species of mycobacteria, there is strong convergence in the types of redox and lipid-centered cellular pathways that ANTAR controls across all mycobacteria.

### ANTAR-target RNAs act as cis-regulatory elements controlling the expression of select genes

Previous work describes ANTAR as a transcription antiterminator that controls genes post-transcription initiation(34)(6)(35)(4). To investigate if phosphoactivation of ANTAR might similarly play a regulatory role in mycobacteria, we analyzed the RNA sequences within their genomic contexts (Figure 6A). We found representatives where the dual stemloops of the ANTAR-target RNA harbour the ribosome binding site (RBS) for the gene immediately downstream (Figure 6A). Based on this we hypothesized that ANTAR binding would prevent access of the RBS to the ribosome and prevent translation initiation. In the absence of ANTAR∼P, the RBS would be accessible, resulting in translation.

**Fig 6.**
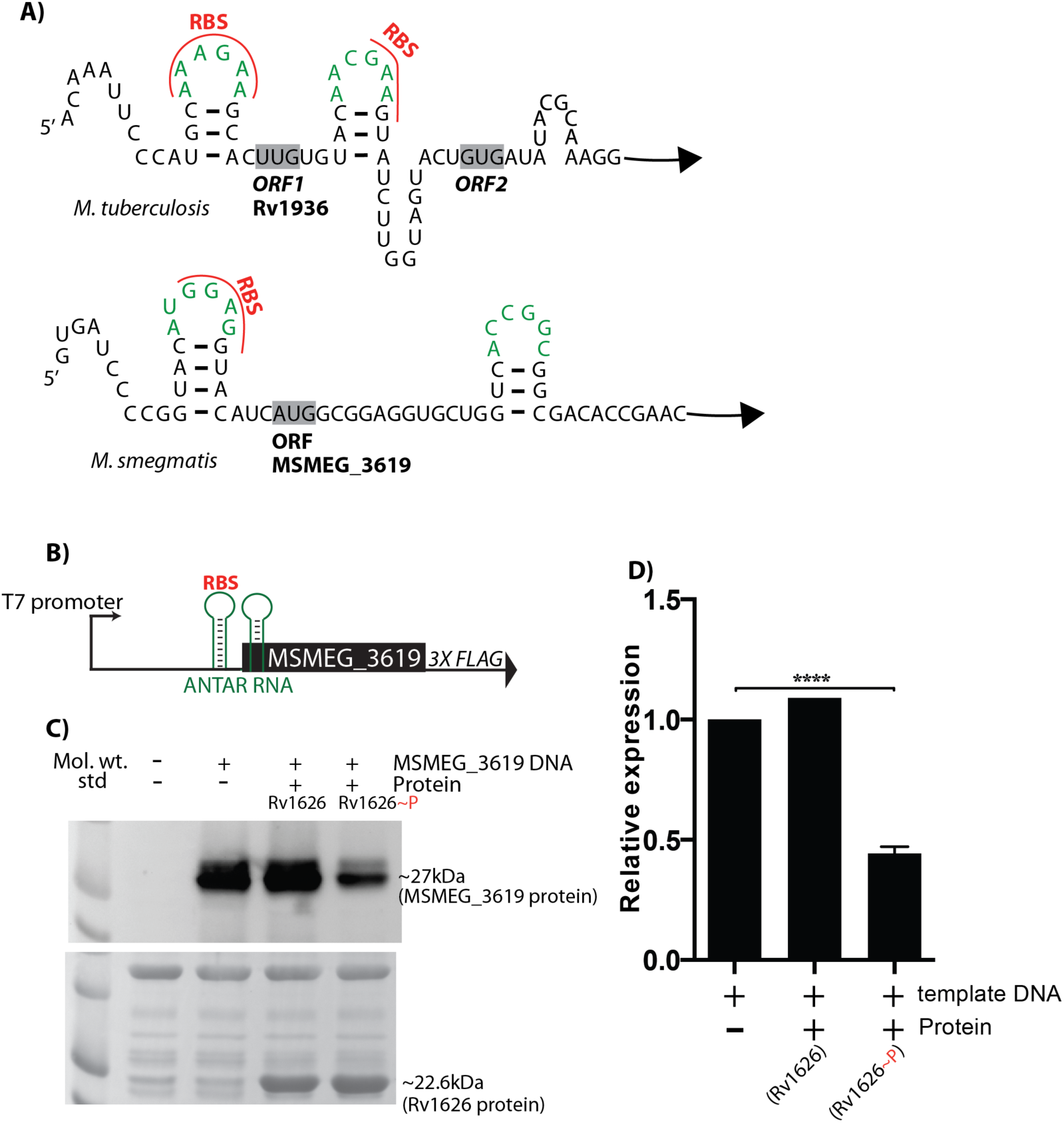
ANTAR-target RNAs act as cis-regulatory elements that control translation. A) Schematic showing predicted ANTAR-target RNAs upstream of the Mtb Rv1936 and *M. smegmatis* MSMEG_3619 genes. Dual stem loops of ANTAR-target RNAs with base paried region (dashed line) and loops (green) are shown. NCBI ORF-finder was used to predict ORF start positions (marked in bold). Two putative start sites were found for Rv1936, as indicated. Ribosome binding sites (RBS, shown in red) lie within the ANTAR RNA stem loop structure. Based on this, ANTAR-binding would occlude the ribosome binding site, preventing translation initiation. B) *In vitro* transcription-translation coupled assay was used to examine the effect of ANTAR-target RNAs on the translation of the downstream gene. Schematic (top) shows details of the DNA template containing a T7 promoter, predicted ANTAR-target RNA (green) and the downstream MSMEG_3619 gene fused to a 3X FLAG tag. RBS (red) is marked. C) Protein synthesized using this assay is visualized on a western blot detected using anti-flag antibodies (top). Ponceau stained image of the blot (bottom) confirms equal loading. Protein synthesis is comparable in the absence of Rv1626 or the presence of unphosphorylated Rv1626, whereas it shows a decrease upon addition of Rv1626∼P (Rv1626D65E) protein. D) Quantification of the blot from panel B shows 50% decrease in protein levels in the presence of Rv1626∼P protein.

To test this hypothesis, we reconstituted ANTAR-RNA regulation *in vitro* using a commercially available purified *E.coli* translation system. In this assay, purified DNA containing a T7 promoter followed by a bioinformatically predicted *M. smegmatis* ANTAR-target RNA in tandem with its downstream MSMEG_3619 gene with a C-terminal 3X FLAG tag sequence, was provided as template for transcription coupled with translation (schematic in Figure 6B). Proteins synthesized in this assay were resolved using electrophoresis and the level of protein synthesis was quantified on a western blot, probed with anti-FLAG tag antibodies (Figure 6C). We find greater than 50% decrease in protein synthesis in the presence of ANTAR∼P when compared to unphosphorylated ANTAR protein or the complete absence of ANTAR protein (Figure 6D). This confirms that ANTAR regulates genes at the level of translation, most likely by blocking the ribosome binding site and preventing translation initiation.

While we demonstrate that ANTAR can modulate translation, other mechanisms of control may also exist. For example, in addition to ANTAR-target RNAs that harbour RBS sites we also found other examples where the dual stemloops of the ANTAR-target RNA possess alternate base-pairing potential (Figure S5B). This would result in a mutually exclusive alternate RNA conformation. In these examples, the dual stemloop RNAs and the alternate conformation are upstream of the RBS and hence unlikely to control translation initiation. In these cases we speculate that the alternate conformation may affect transcription of the downstream mRNA (akin to rho-independent transcription terminators) whereas ANTAR binding to the dual stemloop conformation may elicit an opposite effect on the transcript. In both these examples, ANTAR in mycobacteria would function as an RNA-binding regulator that controls expression of select mRNAs via *cis-* regulatory RNA elements.

### ANTAR activation induces an amplified transcriptomic response, upregulating reductive metabolism in the cell

Our computational approach uses prior knowledge of what defines ANTAR-RNA recognition, as a starting point to search for ANTAR-target RNAs in actinobacteria. In parallel we employed an unbiased approach to assess the effects of ANTAR-mediated regulation in *mycobacterial* cells. To this end, we performed transcriptome analyses of wild type *M. smegmatis* cells compared to cells overexpressing ANTAR∼P. Since the physiological signal/conditions leading to ANTAR phosphorylation are not yet known, we performed a gain-of-function experiment using Rv1626D65E. This allowed us to dial up the ANTAR∼P protein in the cell, in a signal-independent manner. This in turn allowed us to identify mRNAs whose expression is linked to ANTAR∼P. Next-generation sequencing of the transcriptome reveals 45 unique transcripts with altered abundance in cells overexpressing ANTAR∼P compared to control (Figure 7A and Figure S6A-C). 39 of these transcripts show greater than 2-fold decrease in abundance in ANTAR∼P conditions, with the remaining 6 transcripts showing an increase, all within a p-value cut off ≤0.05 (Figure S6A-C). The observed expression changes were independently validated using qRT-PCR (Figure 7B). The response to ANTAR∼P overexpression was not limited to the first gene in an operon and we consistently observed expression changes across the operon.

**Fig. 7.**
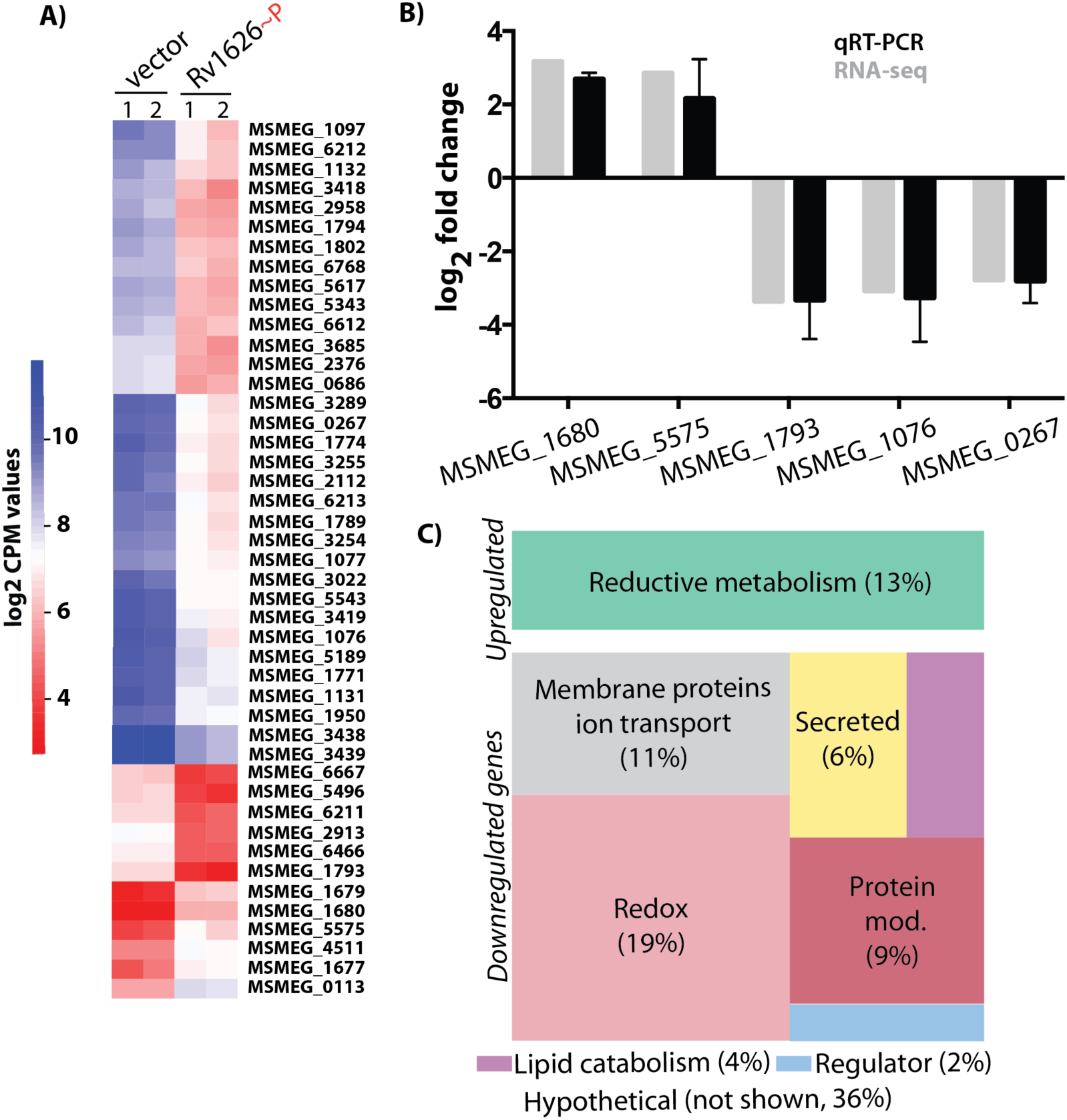
Amplified transcriptomic response to overexpression of ANTAR∼P in M. smegmatis cells. Transcriptomic profiles from log phase cultures of *M. smegmatis* expressing Rv1626∼P (Rv1626D65E) or empty vector were compared using RNA-seq. A) Heat map shows the log_2_ CPM values for 45 genes that showed >2-fold differential expression (p-value≤0.05). Biological replicates were analyzed for gene expression changes using edgeR. 6 genes show upregulation in the presence of Rv1626∼P whereas 39 genes are downregulated. B) qRT-PCR was performed in triplicates, on total RNA extracted from *M. smegmatis* cells overexpressing Rv1626∼P or empty vector. Transcript levels were measured for 5 of the differentially expressed genes identified via RNA-seq. Log_2_ fold change in gene expression measured using qRT-PCR (black) correlates with RNA-seq (grey). C) Differentially expressed genes identified via RNA-seq were categorized based on Gene Ontology terms, gene annotations and manual curation. Treemap shows cellular processes/functions of genes that are upregulated (6 genes) or downregulated (39 genes) upon Rv1626∼P overexpression. Distribution of genes in each category is shown (%). Upregulated genes were enriched in functions related to reductive metabolism. Processes involving oxidative stress response, oxidative phosphorylation, ion transport, etc. were over represented in the downregulated genes (detailed description of genes is shown in Figure S6E).

The transcriptome analysis identified 45 genes whose expression was altered with ANTAR∼P overexpression. We investigated if these genes are also functionally connected. First we performed a GO enrichment analysis(36)(37), which yielded only broad annotations such as membrane protein, conserved hypothetical etc. We then combined functional information from previous work, the STRING database(38) along with GO enrichment for each gene, to more reliably predict pathways where these genes are involved (Figure 7C and S6E). Remarkably, these genes fall into very similar functional categories. The majority of down-regulated genes are involved in oxidative stress response or oxidative processes. This includes enzymes of the respiratory chain complex, several membrane bound ion transporters and aquaporins, a transcriptional regulator ChaB that controls ion transport, multiple FAD/NAD dependent redox-active enzymes and a catalase responsible for defense against ROS/RNS species (see Figure S6E and Table S4 for details). Apart from a very obvious connection to oxidative processes, the list of downregulated genes also includes genes involved in lipid metabolism-such as a glycosyl transferase (MSMEG_1097), lipases (MSMEG_2913, 0267) and lipid transport associated proteins (MSMEG_3418). In the presence of ANTAR∼P protein, these oxidative processes are likely to be down regulated

Only 6 out of 45 differentially expressed genes are upregulated in the ANTAR∼P conditions (Figure 7C and S6E). All of these are directly linked to reductive metabolic processes such as nitrogen and sulfur assimilation. The maximum upregulation is seen for an operon involved in nitrogen metabolism. This operon includes AspA (MSMEG_1677), an aspartate ammonia lyase that catalyzes the reversible conversion of aspartate to fumarate to affect both aspartate and glutamine metabolism, and an amidase (MSMEG_1679) whose substrate is not known. Genes downstream of these are implicated in reductive purine biosynthesis (MSMEG_1681) and purine/pyrimidine transport (MSMEG_1683). Another example of upregulation is of TauC (MSMEG_0113) is a permease that allows scavenging of taurine –a source of sulfur for asssimilation into reduced sulfur metabolites such as mycothiol. Sulfur/cysteine starvation induces the TauABCD operon of *E.coli*(39). Concomitant with an increase in nitrogen and sulfur assimilatory processes, we find that a major repressor of sugar transport(40) (MSMEG_5575) is upregulated in ANTAR∼P cells. This would result in a decreased ability of cells to utilize carbohydrates as a carbon source. Another upregulated gene-MbtE (MSMEG_4511) is involved in the biogenesis of mycobactins(41)(42). These are siderophores that are reponsible for scavenging iron from iron-deplete host environments. This increase in iron scavenging is likely required to cater to the increase in reductive processes. Overall, the transcriptomic analysis showed a very specific subset of 45 genes linked to oxidative and reductive processes, as being responsive to ANTAR∼P.

Our bioinformatic search had predicted only four ANTAR-target RNAs in the *M. smegmatis* genome, yet the gene-expression profiles changed for ∼45 genes. This suggested that not all transcripts were altered due to direct interactions with ANTAR. To distinguish which of these altered transcripts are directly bound by ANTAR and which ones show changes due to downstream effects of ANTAR signaling, we analyzed the transcript sequences along with their upstream region of 100 bases for structures resembling ANTAR-target RNAs. The upstream regions of 13 of these 45 transcripts possess a 2-stemloop structure (Figure S6D). However, none of these structures met the criteria for ANTAR recognition. Additionally, using EMSAs we investigated if these transcripts could interact with the ANTAR protein and we found that none of these RNAs bind ANTAR∼P protein *in vitro*. This strongly suggests that the observed transcriptomic changes are not a result of direct binding by ANTAR, but are instead a *global response* to ANTAR∼P in cells.

### ANTAR RNAs control the reductive/oxidative capacity of the cell

The remarkable convergence of ANTAR-linked genes to reductive/oxidative metabolism inspired us to ask if the link between ANTAR and the cellular metabolic state could be recapitulated *in vivo*. Since cells overexpressing ANTAR∼P showed upregulation of genes involved in reductive metabolism, we first investigated the general reductive capacity of cells. For this we used resazurin, a cell-permeable, redox sensitive dye which gains fluorescence (excitation at 560nm and emission at 582nm) when reduced to resorufin (Figure 8A). We find that cells overexpressing ANTAR∼P show significantly higher resorufin fluorescence when compared to cells expressing the empty vector. This difference was observable in cells grown to exponential phase in rich media, akin to conditions used for the transcriptome analysis (Figure 8A), suggesting an ANTAR∼P dependent increase in reductive capacity. This then led us to hypothesize that ANTAR∼P overexpressing cells would possess a growth advantage under oxidative stress, since the increased reductive capacity is likely to have a neutralizing effect. To test this, we examined the ability of cells to grow under increasing levels of oxidative stress induced by H_2_O_2_ (Figure 8B and Figure S7). Indeed, as predicted we find that cells overexpressing ANTAR∼P show a growth advantage compared to wild type, when subjected to oxidative stress. Notably, these differences are prominent in moderate levels of oxidative stress (between 0.3 and 1mM H_2_O_2_), whereas at levels of H_2_O_2_ greater than 3mM both strains behave identically and fail to grow. This suggests that ANTAR∼P dependent regulation enables a fine-tuned redox response in cells.

**Fig 8.**
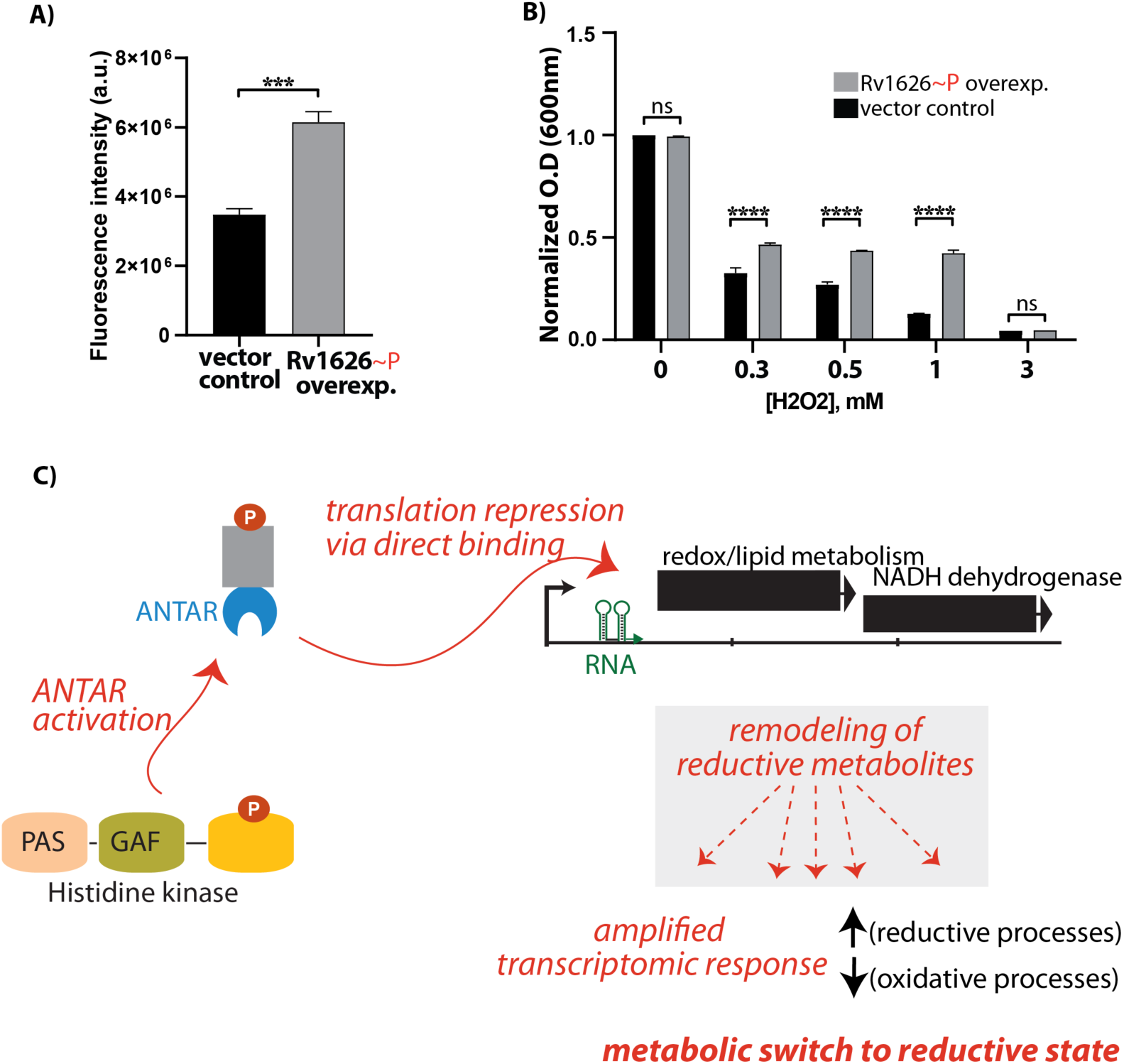
ANTAR∼P alters the metabolic state of M. smegmatis cells. A) Reductive capacity of *M. smegmatis* cells expressing Rv1626∼P (Rv1626D65E) or empty vector was measured using resazurin dye. Reduction of resazurin to resorufin results in gain in fluorescence. Bar graph showing resorufin fluorescence normalized to cell density, shows higher reductive potential in Rv1626∼P cells compared to empty vector. Data from biological triplicates is plotted with p-value of 10^-3^ (***). B) Bar graph shows normalized cell density (O.D. at 600nm) for Rv1626∼P overexpressing cells (grey) and empty vector (black) with increasing H_2_O_2_ concentration. Rv1626∼P cells show a significant (p-value 10^-4^) increase in their ability to grow in H_2_O_2_ concentrations ranging from 0.3 to 1mM. Growth of both strains is comparable in the absence of H_2_O_2_ or at 3mM H_2_O_2_. C) Proposed model for ANTAR-RNA-mediated metabolic control in *mycobacteria*. Activation of ANTAR protein by the cognate histidine kinase, results in recognition and direct binding of ANTAR-target RNA targets. This leads to translational repression of lipid metabolism and related redox enzymes. This constitutes the *direct* response to ANTAR. Repression of relevant enzymes is further amplified into a *global* regulatory response via transcriptomic changes wherein genes/operons involved in reductive metabolism are upregulated and those involved in oxidative processes are downregulated. Based on the direct targets of ANTAR, it is likely that the steps leading to this amplified response involves remodeling of the reductive metabolite pool in the cell (grey box).

Put together, these results reveal a common theme wherein ANTAR controls the metabolic state of the cell (model proposed in Figure 8C). A heirarchy of signaling is established, where ANTAR directly represses the translation of genes/operons encoding specific metabolic and redox-dependent enzymes, by interacting with upstream RNA motifs. This regulation likely results in an increase in the cellular pool of reductive small molecule species, which ultimately manifests as an amplified global transcriptomic response that involves several redox-sensitive processes.

## DISCUSSION

The motivation for this study stems from our observation that to date very few RNA families have been described in actinobacteria; even though this phylum comprises several human/plant pathogens and bacteria that possess complex lifestyles that likely require cellular plasticity(43)(44). We reasoned that a known RNA-binding domain such as ANTAR would provide an excellent handle to identify non-coding RNAs as well as understand the roles that RNAs may play in actinobacteria.

Here we report the discovery of a family of structurally conserved RNAs in actinobacteria and elucidate its role in regulating the metabolic state of mycobacterial cells. We find that these RNAs specifically interact with ANTAR domain proteins in a phosphorylation-dependent manner. ANTAR-target RNAs in mycobacteria reside upstream of transcripts encoding redox-dependent enzymes and lipid metabolism enzymes and recruitment of ANTAR proteins to these transcripts results in translational repression of these genes. Remarkably, the cell’s response to ANTAR regulation is multilayered and extends beyond the genes that are directly repressed. We find an ANTAR-dependent global transcriptomic response wherein oxidative enzymes, the ETC, oxidative stress response proteins and transporters are repressed and multiple reductive processes are upregulated; ultimately committing the cell towards reductive metabolism. Combining these results with phenotypic analyses of *M. smegmatis* cells, we present several lines of evidence that gain of ANTAR function increases the overall reductive state of the cell.

The *direct* binding sites for ANTAR (i.e. the RNA targets) are present in the UTRs of lipid metabolism and related redox enzymes. We show these genes to be translationally repressed by ANTAR∼P. In mycobacteria, lipid breakdown as well as biosynthesis is strongly tied to the redox environment of the cell(45). The break down of lipids is an oxidative process. Additionally, mycobacteria are known to produce specific fatty acids and lipids as a response to oxidative stress(46)(47). Under conditions of reductive metabolism, oxygen-dependent processes such as lipid metabolism may need to be repressed. Our data suggests that these two metabolic outcomes are integrated via ANTAR regulation.

The ability of mycobacteria to sense oxygen and oxidative stress and the sensors involved in these has been the subject of study for decades(48)(49). However, recent studies highlight the metabolic flexibility of these bacteria, evident in their ability to utilize nitrogen and sulfur for growth and survival(50). Nitrogen metabolism is relevant for growth and virulence of pathogenic mycobacteria like Mtb in oxygen-starved niches such as the lung granulomas. Saprophytic species such as *M. smegmatis* experience a much higher diversity of nitrogen sources in the environment and are also likely to rely on nitrogen for growth under certain conditions. By controlling the reductive state of cells, ANTAR may be an important player in the metabolic flexibility across mycobacteria.

Our work elucidates the molecular details of signaling events downstream of ANTAR-RNA binding that link ANTAR to a reductive metabolic state. Future work would need to address what the upstream signaling events may be that lead to ANTAR activation. In mycobacteria the majority of ANTAR domains appear in tandem with GAF(17)/PAS(18) domains, or phospho-receiver domains (as in Rv1626) that interact with PAS_GAF-containing histidine kinases (as in Rv3220c). Both GAF and PAS domains are ubiquitous signal sensing domains where GAF domains sense cyclic second messengers such as cAMP and cGMP(17). PAS domains also bind diverse ligands like FMN, heme, small organic molecules and respond to signals ranging from light, oxygen to redox(18). We speculate that the signals leading to ANTAR activation would be related to redox and lipid signaling and might be sensed through the PAS-GAF domains of Rv3220c. Mycobacterial genomes typically contain several ANTAR coding genes, and future work will address the roles of individual ANTAR systems and how they tie in with the observed metabolic switching in mycobacteria.

Key to our discovery of ANTAR-target RNAs in actinobacteria was a modified approach where we revamped the computational method currently used(21) for identifying structurally conserved RNAs. We now describe an approach wherein removing the bias introduced by “very similar RNA sequences” and allowing more diversity in the search model allows identification of ANTAR-target RNAs in actinobacteria. The implicit logic and search method is broadly applicable to finding any RNA family.

Mycobacteria are known to possess a large number of leader-less transcripts(22)(51), which would suggest that regulatory RNAs that otherwise reside in the 5’ untranslated regions of transcripts may not play a significant role in such bacteria. However, our finding that simple two stem loop RNA motifs (like ANTAR-target RNAs) overlapping with the ribosome binding site and the ORF start sites can control translation, provides new insights into how regulation of even leader-less transcripts can be achieved via regulatory RNAs.

The cellular response to ANTAR is hierarchical. ANTAR proteins directly bind and regulate few target transcripts and this has a cascading effect, which amplifies into a broader transcriptomic response. The exact mechanism of this amplification is unclear. We were unable to find any transcription factors that were directly controlled by ANTAR and whose regulation could result in an amplified transcriptional response. Based on the nature of the genes under direct ANTAR control, we speculate that this amplification is mediated via a remodeling of metabolites in the cell. In any case, the complex, multilayered architecture of ANTAR regulation is remarkable and underlines the role of RNAs in nuanced regulatory outcomes for the cell.

## METHODS

### Bacterial strains and Media

*M. smegmatis* MC^2^155 was cultured in Difco MiddleBrook 7H9 broth with 0.5% glycerol and 0.05% Tween-80 (or 7H10 Agar with 0.2% glycerol) enriched with 10% oleic acid, albumin, dextrose and catalase (BD BBL™). When required kanamycin (25 µg/ml), hygromycin (50 µg/ml) were added. *E. coli* DH5α used for cloning and BL21(DE3) cells used for recombinant protein expression were grown in LB broth, supplemented with kanamycin (50 µg/ml)/hygromycin (50 µg/ml) as required. Rv1626D65E gene was cloned in pMV261 plasmid under the control of an hsp60 promoter and expressed in *M smegmatis* MC^2^155 strain by electroporation of ∼1µg of the plasmid into electrocompetent cells at 2500 V, 25µF, and the pulse controller (Bio-Rad) set to 1000ohms. Empty vector control strain expressed the pMV261 plasmid.

### Recombinant protein expression and preparation

Plasmids encoding N-terminal 6-his tagged proteins-Rv1626, Rv1626D65E, EutVD54E (ANTAR∼P phosphomimic protein from *E. faecalis*) and EutW, proteins were constructed as described in Table S5. These were transformed in to *E. coli* BL21 (DE3) cells and grown to OD 0.8, induced by addition of 0.5mM IPTG and subsequently grown at 25°C overnight. Protein was purified following previous protocols(5). Briefly, Cells were lysed in buffer containing 25mM sodium phosphate pH7.0, 500mM sodium chloride, 2mM β-mercaptoethanol, 10% (v/v) glycerol, 1mM PMSF, 1mg/ml DNase and 0.1mg/ml lysozyme. Following centrifugation at 12,000Xg for 20 mins, the supernatant was loaded on TALON affinity resin (Clontech), washed with buffer containing 10mM imidazole and eluted in buffer supplemented with 400mM imidazole. Purified protein was dialyzed against buffer containing 25mM sodium phosphate pH7.0, 150mM sodium chloride, 2mM β-mercaptoethanol and 10% (v/v) glycerol for 12 hours, along with TEV protease (1mg per 20mg of recombinant 6his protein). The sample was then passed through Ni-NTA agarose resin (Qiagen), to obtain pure protein which was finally dialysed against 20mM Tris pH8.0, 150mM sodium chloride, 2mM β-mercaptoethanol and 10% (v/v) glycerol. Absorbance ratios 260/280nm ∼0.7 suggested that no DNA or RNA contamination was present in the protein samples.

### In vitro phosphorylation assays

Relevant proteins were incubated in buffer containing 20mM Tris pH 8.0, 150mM NaCl, 2 mM MgCl_2_ and 1 mM ATP (spiked with γ-^32^P ATP) for 30 minutes at 24°C. For EutW containing assays 1mM ethanolamine was used to activate phosphorylation. Samples were subjected to SDS-PAGE in duplicate and further stained with Coomassie Brilliant blue or visualized using a PhosphorImager (Amersham). Autoradiograms were quantified using imageJ.

### RNA preparation

RNA was prepared for *in vitro* studies, as described previously(52). DNA templates were created by PCR using relevant primers (details in Table S5), adding the T7 promoter sequence at the 5’ end of the template. In vitro transcription was performed using T7 RNA polymerase, at 37°C for 2.5 hours. Reactions were quenched with dye containing 8 M urea and products were resolved by denaturing 6% PAGE and visualized by UV shadowing. RNA was extracted using passive elution, following which ethanol precipitation was performed and the RNA was quantified using absorbance at 260nm.

### In-line probing assays

RNA corresponding to the predicted ANTAR-target RNA upstream of *M. gilvum* Esterase gene (sequence details in Table S5) was produced by in vitro transcription and 5’-radiolabeled with γ-^32^P ATP as described previously(53). RNA (10fmol, ∼400CPS) was incubated in buffer containing 50mM Tris pH 8.3, 100 mM KCl and 20 mM MgCl_2,_ for 40 hours at 24°C and subjected to electrophoresis on a 10% Urea polyacrylamide gel. Controls: partial digestion by RNase T1 (cleavage after G), OH ladder (cleavage at every position) and unreacted RNA were run alongside the in-line sample. Gels were imaged using a PhosphorImager (Amersham) and analyzed using imageJ.

### Electrophoretic mobility shift assays

RNAs were 5’ end-labeled using γP^32^-ATP and incubated with increasing concentrations of protein at 24°C for 1 hour in buffer comprising 20mMTris, pH 8, 10mM DTT, 150mM NaCl, 5mM MgCl_2_, 10% glycerol (v/v) and yeast total RNA (2.5 ng/µl). Samples were resolved on 5% TBE polyacrylamide gels (acrylamide:bis 80:1) that were pre-run for 15 minutes at 40 volts, and subsequently at 100 volts for 1 hour. Gels were exposed and visualized using a PhosphorImager (GE Health Sciences). The gels were quantified using imageJ and data plotted using GraphPad Prism.

### In vitro transcription-translation coupled assay

DNA template for this assay was obtained using PCR (details of primers in Table S5), and consisted of a 757-bp amplicon with the T7 promoter followed by the 5’UTR and full length MSMEG_3619 gene, to which a C-terminal 3XFLAG tag was appended. PCR products were purified using phenol-chloroform extraction and ethanol precipitation and the resulting pellet was dissolved in buffer containing 10mM Tris-HCl pH 8. The assay was performed using the PURExpress In Vitro Protein Synthesis Kit (E6800S New England Biolabs, Inc.), following manufacturer’s instructions. Where needed, the reaction was supplemented with either Rv1626D65E protein (30μM) or Rv1626 protein (30μM). Samples were subjected to SDS-PAGE and transferred on to a nitrocellulose membrane using the Trans-Blotcell (Bio-Rad) at 110V for 90 min. Proteins synthesized in this assay were detected using anti-FLAG antibodies (F3165 SIGMA, 1:4000 dilution) and the Anti-mouse IgG, HRP-linked Antibody (1:4000 dilution) (7076S-Cell Signaling Technology) and developed using the ECL detection system (RPN 2232 GE Healthcare). Blots were quantified using imageJ and data plotted using GraphPad Prism.

### Transcriptome analysis

Total RNA was extracted from exponentially growing (OD 0.8) *M smegmatis* MC^2^155 cells by bead beating with Glass Mill Beads, 0.5 mm (4×30 sec beating and cooling cycles) in presence of Trizol. RNA was further purified using Direct-zol RNA Miniprep Zymo-Spin IIC Columns and further subjected to DNAse treatment and cleanup using RNA Clean & Concentrator-5/25 Zymo-Spin IC Columns. RNA was quantified at every stage using Qubit and Bioanalyser using Agilent Bioanalyzer 2100-RNA Pico Kit or RNA 6000 Nano Kit. Ribosomal RNA was removed using Ribo-Zero rRNA removal kit for bacteria (Illumina MRZB12424) as per manufacturer instructions and samples further purified using RNA Clean and Concentrator-5 (Zymo Research R1015). cDNA Libraries were constructed (Illumina TruSeq RNA Sample prep Kit v2) and the amplified library was quantified using qubit and subsequently analyzed using the Agilent 2100 Bioanalyzer DNA high sensitivity chip. Sequencing was carried out using the Illumina MiSeq Reagent Kit v3 (2×75 bp, 150 cycle format). Experiments were performed with two biological replicates.

RNA-seq reads were checked for sequencing quality using FastQC (https://www.bioinformatics.babraham.ac.uk/projects/fastqc). Reads mapping to rRNAs were removed using SortMeRNA v2.1 with e-value≤1e-4. Reads were mapped to the *M. smegmatis* MC^2^155 genome (NCBI accession: NC_008596.1) using BWA-MEM v0.7.12-r1039(54) and reads mapping uniquely (mapping quality≥20) were considered for further analyses. These were processed using SAMtools(55), following which the coverage for each gene was calculated using the default parameters in BEDtools(56). Gene-specific read counts were then used for differential gene expression analysis using the EdgeR v3.24.1 quasi-likelihood pipeline in R as described(57). Genes with >10 reads and appearing in both biological replicates were taken and gene-wise read counts were normalized to library size using upperquartile method. Log_2_ CPM (counts per million) values were calculated. Two-fold differentially expressed genes with FDR-corrected p-value≤0.05 were considered significant. All analyses and plots were obtained using R v3.5.3.

### Quantitative real-time PCR analysis

200 ng of total RNA from *M. smegmatis* wild-type or Rv1626D65E over-expression strain was used for DNase I (AM2238, Thermo Fisher Scientific) treatment. cDNA was synthesized with random primers (48190011, Thermo Fisher Scientific) and SuperScript III reverse transcriptase (18080085, Thermo Fisher Scientific) according to the manufacturer’s protocol. cDNA quantification was done by real-time PCR on an ViiA 7 real-time PCR system (Thermo Fisher Scientific) using Maxima SYBR Green/ROX qPCR Master Mix (K0222, Thermo Fisher Scientific). 5 ng of cDNA was used as a template. *SigA* was used as an internal normalization control. All qRT-PCRs were performed in triplicates using two independent biological replicates. Statistical significance was determined using a Student *T*-test (GraphPad Prism 7). The primer sequences for qRT-PCRs are listed in Table S5.

### H _2_O_2_ induced oxidative stress assays

*M*. *smegmatis* MC^2^155 strains with empty vector or overexpression of Rv1626D65E were grown to log phase (OD_600nm_ ∼0.8–1.0). Cells were pelleted and re-suspended in 7H9 broth containing 0.05% Tween-80 and 0.5% glycerol, supplemented with 10% ADN (5g/L albumin, 2g/L dextrose, 0.85g/L NaCl). The culture was then diluted 100 times into the same media and allowed to grow up to OD_600nm_ of 0.3 (∼ 12 hours). These cells were then treated with the indicated concentrations of H_2_O_2_ (0, 0.3, 0.5, 1 or 3 mM) and allowed to grow till late log phase (OD_600nm_=2). OD_600nm_ was measured every 1 hour (Figure S7). Maximum growth rate differences between empty vector and Rv1626D65E overexpressing strains were observed at 25 hours post H_2_O_2_ treatment. Hence, cell density (after 25 hours) for different H_2_O_2_ concentrations was normalized against cell density of untreated cells. All experiments were performed as biological triplicates. Statistical analysis was carried out using two-way Annova with a p-value threshold of 0.05 and plotted using Graphpad v6.07.

### Alamar Blue Assay

*M*. *smegmatis* MC^2^155 strains with empty vector or overexpression of Rv1626D65E were grown to early log phase (OD_600nm_ ∼0.3) and treated with 10%(v/v) of Alamar Blue reagent (Invitrogen: DAL1025). Cells were allowed to grow overnight at 37°C at 200 rpm and fluorescence was measured using Fluorolog® - Horiba with excitation at 560nm and emission being recorded at 582nm. OD_600nm_ was measured in parallel and used to calculate fluorescence normalized to cell density. All experiments were performed as biological triplicates. Statistical analysis was carried out using unpaired two-tailed t-test with a p-value threshold of 0.05 and plotted using Graphpad v6.07.

### Computational analyses

#### Sequence and domain analyses of ANTAR proteins

Number of ANTAR proteins and its domain architectures in different organisms were obtained from Pfam v31.0 (PF03861). Statistical analysis was carried out using one-way ANOVA with p-value threshold of 0.05 in GraphPad v6.07. Sequence alignment of three ANTAR proteins from *E. faecalis, M. tuberculosis* and *M. smegmatis* was carried out using Clustal Omega v1.2.4. Domain architecture for *M. tuberculosis* ANTAR protein is obtained from PDB: 1S8N.

#### Covariance search to predict RNAs

17 RNA sequences (Seed 1) used as search model in a previous study(4), were taken. These RNAs were subjected to cmbuild analysis as part of Infernal v1.0.2(58). We used the –cmaxid option under cmbuild that allowed us to specify a cut off of 51% sequence identity to be imposed during clustering. This resulted in a single cluster of 17 sequences, which showed a covariation score (CM score) of 0.75. Higher CMscore was taken as an indication of highly similar sequences. We then created an Intermediate Seed, by combining 12 sequences from Seed1 with 39 ANTAR-target RNA sequences from diverse bacteria (shown in Table S1). These 39 RNAs were picked manually, from the list of ANTAR-target RNAs reported previously(4). This Intermediate Seed was again subjected to cmbuild analysis and showed 5 clusters where the biggest cluster consisted of 47 sequences (CM score 0.61) and the other 4 were outliers. These 47 sequences (Seed 2) were then used as search model against 128 mycobacterial strains (from RefSeq v76) using Infernal v1.0.2. Hits with an arbitrary bit score ≥10 and within 500nt upstream of ORFs were retained. RNAs that we validated experimentally (7 RNAs) if used as a stand-alone search model, result in identifying additional -target RNAs that are validated in vitro. Based on secondary structure (analyzed using in-house PERL scripts and manual curation) we assigned RNA hits to 2 categories. Category 1 includes RNAs matching the search model and single nucleotide point variants. Category 2 consists of RNAs where one of the two hairpins deviates from the search model. Consensus RNA secondary structure was drawn using R-scape. Description of genes downstream to the predicted ANTAR-target RNAs was obtained using NCBI annotations and manual curation and used to cluster the genes into processes/functions. This was visualized using treemap v2.4.2 R package.

#### Gene Ontology (GO) analysis

45 differentially expressed genes identified via transcriptomic analysis were subjected to GO analysis using the gene ontology annotation pipeline in BLAST2GO(59) and GOA(60). These gave us GO ids and terms for the 45 differentially expressed genes. We then subjected these to GO term enrichment using GO-TermFinder v0.82 with p-value threshold of 1.0. These GO terms, along with NCBI gene annotations and manual curation were then used to categorize the genes into processes/functions. Plots were obtained using Treemap v2.4.2 R package.

## ACKNOWLEDGEMENTS

We acknowledge funding and support from the Wellcome DBT-India Alliance (Intermediate fellowship IA/I/14/2/501521 to AR) and National Center for Biological Sciences, India. We thank Dr. Vineetha Zachariah and Sumedha Agashe for initiating the *in vivo* studies in *mycobacteria* and Dr. Sandhya Visweswariah, IISc, India for reagents/ access to BSL2 facilities. We are grateful to Sunil Laxman (inStem) and Akash Gulyani (InStem), for critical comments on the manuscript.

## REFERENCES

1. Nahvi, A., Sudarsan, N., Ebert, M. S., Zou, X., Brown, K. L., and Breaker, R. R. (2002) Genetic control by a metabolite binding mRNA. Chem. Biol. 9, 1043

2. Winkler, W. C., and Breaker, R. R. (2005) Regulation of bacterial gene expression by riboswitches. Annu. Rev. Microbiol. 59, 487–517

3. Collins, J. A., Irnov, I., Baker, S., and Winkler, W. C. (2007) Mechanism of mRNA destabilization by the glmS ribozyme. Genes Dev. 21, 3356–68

4. Ramesh, A., DebRoy, S., Goodson, J. R., Fox, K. A., Faz, H., Garsin, D. A., and Winkler, W. C. (2012) The Mechanism for RNA Recognition by ANTAR Regulators of Gene Expression. PLOS Genet. 8, e1002666

5. Fox, K. A., Ramesh, A., Stearns, J. E., Bourgogne, A., Reyes-Jara, A., Winkler, W. C., and Garsin, D. A. (2009) Multiple posttranscriptional regulatory mechanisms partner to control ethanolamine utilization in Enterococcus faecalis. Proc. Natl. Acad. Sci. U. S. A. 106, 4435–4440

6. Chai, W., and Stewart, V. (1998) NasR, a novel RNA-binding protein, mediates nitrate-responsive transcription antitermination of the Klebsiella oxytoca M5al nasF operon leader in vitro. J. Mol. Biol. 283, 339–51

7. Ueki, T., and Lovley, D. R. (2010) Novel regulatory cascades controlling expression of nitrogen-fixation genes in Geobacter sulfurreducens. Nucleic Acids Res. 38, 7485–99

8. Arnvig, K. B., Cortes, T., and Young, D. B. (2014) Noncoding RNA in Mycobacteria. Microbiol. Spectr.

9. Arnvig, K. B., Comas, I., Thomson, N. R., Houghton, J., Boshoff, H. I., Croucher, N. J., Rose, G., Perkins, T. T., Parkhill, J., Dougan, G., and Young, D. B. (2011) Sequence-based analysis uncovers an abundance of non-coding RNA in the total transcriptome of Mycobacterium tuberculosis. PLOS Pathog. 7, e1002342

10. Heueis, N., Vockenhuber, M.-P., and Suess, B. (2014) Small non-coding RNAs in streptomycetes. RNA Biol. 11, 464–469

11. Dambach, M., Irnov, I., and Winkler, W. C. (2013) Association of RNAs with Bacillus subtilis Hfq. PLOS One. 8, e55156–e55156

12. Faner, M. A., and Feig, A. L. (2013) Identifying and characterizing Hfq-RNA interactions. Methods. 63, 144–159

13. Goldman, B. S., Lin, J. T., and Stewart, V. (1994) Identification and structure of the nasR gene encoding a nitrate- and nitrite-responsive positive regulator of nasFEDCBA (nitrate assimilation) operon expression in Klebsiella pneumoniae M5al. J. Bacteriol. 176, 5077–85

14. Shu, C. J., and Zhulin, I. B. (2002) ANTAR: an RNA-binding domain in transcription antitermination regulatory proteins. Trends Biochem. Sci. 27, 3–5

15. Norman, R. a, Poh, C. L., Pearl, L. H., O’Hara, B. P., and Drew, R. E. (2000) Steric hindrance regulation of the Pseudomonas aeruginosa amidase operon. J. Biol. Chem. 275, 30660–7

16. Bateman, A., Birney, E., Cerruti, L., Durbin, R., Etwiller, L., Eddy, S. R., Griffiths-Jones, S., Howe, K. L., Marshall, M., and Sonnhammer, E. L. L. (2002) The Pfam protein families database. Nucleic Acids Res. 30, 276–280

17. Aravind, L., and Ponting, C. P. (1997) The GAF domain: an evolutionary link between diverse phototransducing proteins. Trends Biochem. Sci. 22, 458–459

18. Taylor, B. L., and Zhulin, I. B. (1999) PAS domains: internal sensors of oxygen, redox potential, and light. Microbiol. Mol. Biol. Rev. 63, 479–506

19. Morth, J. P., Feng, V., Perry, L. J., Svergun, D. I., and Tucker, P. a (2004) The crystal and solution structure of a putative transcriptional antiterminator from Mycobacterium tuberculosis. Structure. 12, 1595–605

20. Preu, J., Panjikar, S., Morth, P., Jaiswal, R., Karunakar, P., and Tucker, P. a (2012) The sensor region of the ubiquitous cytosolic sensor kinase, PdtaS, contains PAS and GAF domain sensing modules. J. Struct. Biol. 177, 498–505

21. Nawrocki, E. P., and Eddy, S. R. (2013) Infernal 1.1: 100-fold faster RNA homology searches. Bioinformatics. 29, 2933–2935

22. Shell, S. S., Wang, J., Lapierre, P., Mir, M., Chase, M. R., Pyle, M. M., Gawande, R., Ahmad, R., Sarracino, D. A., Ioerger, T. R., Fortune, S. M., Derbyshire, K. M., Wade, J. T., and Gray, T. A. (2015) Leaderless Transcripts and Small Proteins Are Common Features of the Mycobacterial Translational Landscape. PLOS Genet. 11, e1005641–e1005641

23. Rose, G., Cortes, T., Comas, I., Coscolla, M., Gagneux, S., and Young, D. B. (2013) Mapping of genotype-phenotype diversity among clinical isolates of mycobacterium tuberculosis by sequence-based transcriptional profiling. Genome Biol. Evol. 5, 1849–1862

24. Dinan, A. M., Tong, P., Lohan, A. J., Conlon, K. M., Miranda-CasoLuengo, A. A., Malone, K. M., Gordon, S. V, and Loftus, B. J. (2014) Relaxed Selection Drives a Noisy Noncoding Transcriptome in Members of the Mycobacterium tuberculosis Complex. MBio. 10.1128/mBio.01169-14

25. Li, Y., and Breaker, R. R. (1999) Kinetics of RNA Degradation by Specific Base Catalysis of Transesterification Involving the 2‘-Hydroxyl Group. J. Am. Chem. Soc. 121, 5364–5372

26. Janagama, H. K., Tounkang, S., Cirillo, S. L. G., Zinniel, D. K., Barletta, R. G., and Cirillo, J. D. (2013) Molecular analysis of the Mycobacterium tuberculosis lux-like mel2 operon. Tuberculosis. 93, S83–S87

27. Subbian, S., Mehta, P. K., Cirillo, S. L. G., and Cirillo, J. D. (2007) The Mycobacterium marinum mel2 locus displays similarity to bacterial bioluminescence systems and plays a role in defense against reactive oxygen and nitrogen species. BMC Microbiol. 7, 4

28. Unnikrishnan, M., Constantinidou, C., Palmer, T., and Pallen, M. J. (2017) The Enigmatic Esx Proteins: Looking Beyond Mycobacteria. Trends Microbiol. 25, 192–204

29. Stoop, E. J. M., Bitter, W., and van der Sar, A. M. (2012) Tubercle bacilli rely on a type VII army for pathogenicity. Trends Microbiol. 20, 477–484

30. Bosserman, R. E., Nguyen, T. T., Sanchez, K. G., Chirakos, A. E., Ferrell, M. J., Thompson, C. R., Champion, M. M., Abramovitch, R. B., and Champion, P. A. (2017) WhiB6 regulation of ESX-1 gene expression is controlled by a negative feedback loop in Mycobacterium marinum. Proc. Natl. Acad. Sci. 114, E10772–E10781

31. Díaz, E., Ferrández, A., and García, J. L. (1998) Characterization of the hca cluster encoding the dioxygenolytic pathway for initial catabolism of 3-phenylpropionic acid in Escherichia coli K-12. J. Bacteriol. 180, 2915–2923

32. Vilchèze, C., Weisbrod, T. R., Chen, B., Kremer, L., Hazbón, M. H., Wang, F., Alland, D., Sacchettini, J. C., and Jacobs Jr, W. R. (2005) Altered NADH/NAD+ ratio mediates coresistance to isoniazid and ethionamide in mycobacteria. Antimicrob. Agents Chemother. 49, 708–720

33. Miesel, L., Weisbrod, T. R., Marcinkeviciene, J. A., Bittman, R., and Jacobs Jr, W. R. (1998) NADH dehydrogenase defects confer isoniazid resistance and conditional lethality in Mycobacterium smegmatis. J. Bacteriol. 180, 2459–2467

34. Wilson, S. A., Wachira, S. J. M., Norman, R. A., Pearl, L. H., and Drew, R. E. (1996) Transcription antitermination regulation of the. 15, 5907–5916

35. O’Hara, B. P., Norman, R. a, Wan, P. T., Roe, S. M., Barrett, T. E., Drew, R. E., and Pearl, L. H. (1999) Crystal structure and induction mechanism of AmiC-AmiR: a ligand-regulated transcription antitermination complex. EMBO J. 18, 5175–86

36. Ashburner, M., Ball, C. A., Blake, J. A., Botstein, D., Butler, H., Cherry, J. M., Davis, A. P., Dolinski, K., Dwight, S. S., Eppig, J. T., Harris, M. A., Hill, D. P., Issel-Tarver, L., Kasarskis, A., Lewis, S., Matese, J. C., Richardson, J. E., Ringwald, M., Rubin, G. M., and Sherlock, G. (2000) Gene ontology: tool for the unification of biology. The Gene Ontology Consortium. Nat. Genet. 25, 25–29

37. Mi, H., Huang, X., Muruganujan, A., Tang, H., Mills, C., Kang, D., and Thomas, P. D. (2017) PANTHER version 11: expanded annotation data from Gene Ontology and Reactome pathways, and data analysis tool enhancements. Nucleic Acids Res. 45, D183–D189

38. Snel, B., Lehmann, G., Bork, P., and Huynen, M. A. (2000) STRING: a web-server to retrieve and display the repeatedly occurring neighbourhood of a gene. Nucleic Acids Res. 28, 3442–3444

39. Eichhorn, E., van der Ploeg, J. R., and Leisinger, T. (2000) Deletion analysis of the Escherichia coli taurine and alkanesulfonate transport systems. J. Bacteriol. 182, 2687–2695

40. Titgemeyer, F., Amon, J., Parche, S., Mahfoud, M., Bail, J., Schlicht, M., Rehm, N., Hillmann, D., Stephan, J., Walter, B., Burkovski, A., and Niederweis, M. (2007) A genomic view of sugar transport in Mycobacterium smegmatis and Mycobacterium tuberculosis. J. Bacteriol. 189, 5903–5915

41. Chavadi, S. S., Stirrett, K. L., Edupuganti, U. R., Vergnolle, O., Sadhanandan, G., Marchiano, E., Martin, C., Qiu, W.-G., Soll, C. E., and Quadri, L. E. N. (2011) Mutational and phylogenetic analyses of the mycobacterial mbt gene cluster. J. Bacteriol. 193, 5905–5913

42. LaMarca, B. B. D., Zhu, W., Arceneaux, J. E. L., Byers, B. R., and Lundrigan, M. D. (2004) Participation of fad and mbt genes in synthesis of mycobactin in Mycobacterium smegmatis. J. Bacteriol. 186, 374–382

43. Arnvig, K. B., and Young, D. B. (2009) Identification of small RNAs in Mycobacterium tuberculosis. Mol. Microbiol. 73, 397–408

44. Mentz, A., Neshat, A., Pfeifer-Sancar, K., Pühler, A., Rückert, C., and Kalinowski, J. (2013) Comprehensive discovery and characterization of small RNAs in Corynebacterium glutamicumATCC 13032. BMC Genomics. 14, 714

45. Lovewell, R. R., Sassetti, C. M., and VanderVen, B. C. (2016) Chewing the fat: lipid metabolism and homeostasis during M. tuberculosis infection. Curr. Opin. Microbiol. 29, 30–36

46. Singh, A., Crossman, D. K., Mai, D., Guidry, L., Voskuil, M. I., Renfrow, M. B., and Steyn, A. J. C. (2009) Mycobacterium tuberculosis WhiB3 Maintains Redox Homeostasis by Regulating Virulence Lipid Anabolism to Modulate Macrophage Response. PLOS Pathog. 5, e1000545

47. You, D., Xu, Y., Yin, B.-C., and Ye, B.-C. (2019) Nitrogen Regulator GlnR Controls Redox Sensing and Lipids Anabolism by Directly Activating the whiB3 in Mycobacterium smegmatis. Front. Microbiol.. 10, 74

48. Sherman, D. R., Sabo, P. J., Hickey, M. J., Arain, T. M., Mahairas, G. G., Yuan, Y., Barry 3rd, C. E., and Stover, C. K. (1995) Disparate responses to oxidative stress in saprophytic and pathogenic mycobacteria. Proc. Natl. Acad. Sci. U. S. A. 92, 6625–6629

49. Pagán-Ramos, E., Song, J., McFalone, M., Mudd, M. H., and Deretic, V. (1998) Oxidative stress response and characterization of the oxyR-ahpC and furA-katG loci in Mycobacterium marinum. J. Bacteriol. 180, 4856–4864

50. Agapova, A., Serafini, A., Petridis, M., Hunt, D. M., Garza-Garcia, A., Sohaskey, C. D., and de Carvalho, L. P. S. (2019) Flexible nitrogen utilisation by the metabolic generalist pathogen Mycobacterium tuberculosis. Elife. 8, e41129

51. Cortes, T., Schubert, O. T., Rose, G., Arnvig, K. B., Comas, I., Aebersold, R., and Young, D. B. (2013) Genome-wide mapping of transcriptional start sites defines an extensive leaderless transcriptome in Mycobacterium tuberculosis. Cell Rep. 5, 1121–1131

52. Wakeman, C. A., Winkler, W. C., and Dann 3rd, C. E. (2007) Structural features of metabolite-sensing riboswitches. Trends Biochem. Sci. 32, 415–424

53. Dann, C. E., Wakeman, C. a, Sieling, C. L., Baker, S. C., Irnov, I., and Winkler, W. C. (2007) Structure and mechanism of a metal-sensing regulatory RNA. Cell. 130, 878–92

54. Li, H., and Durbin, R. (2009) Fast and accurate short read alignment with Burrows-Wheeler transform. Bioinformatics. 25, 1754–1760

55. Li, H., Handsaker, B., Wysoker, A., Fennell, T., Ruan, J., Homer, N., Marth, G., Abecasis, G., Durbin, R., and Subgroup, 1000 Genome Project Data Processing (2009) The Sequence Alignment/Map format and SAMtools. Bioinformatics. 25, 2078–2079

56. Quinlan, A. R., and Hall, I. M. (2010) BEDTools: a flexible suite of utilities for comparing genomic features. Bioinformatics. 26, 841–842

57. Chen, Y., Lun, A. T. L., and Smyth, G. K. (2016) From reads to genes to pathways: differential expression analysis of RNA-Seq experiments using Rsubread and the edgeR quasi-likelihood pipeline. F1000Research. 5, 1438

58. Nawrocki, E. P., Kolbe, D. L., and Eddy, S. R. (2009) Infernal 1.0: inference of RNA alignments. Bioinformatics. 25, 1335–1337

59. Conesa, A., Terol, J., García-Gómez, J. M., Talón, M., Robles, M., and Götz, S. (2005) Blast2GO: a universal tool for annotation, visualization and analysis in functional genomics research. Bioinformatics. 21, 3674–3676

60. Huntley, R. P., Sawford, T., Mutowo-Meullenet, P., Shypitsyna, A., Bonilla, C., Martin, M. J., and O’Donovan, C. (2015) The GOA database: gene Ontology annotation updates for 2015. Nucleic Acids Res. 43, D1057–D1063

61. Durbin, R., and Eddy, S. R. (1994) RNA sequence analysis using covariance models. Nucleic Acids Res. 22, 2079–2088

