## Supplementary File for "Non-coding RNAs control metabolic state in mycobacteria"

Supplemental Information

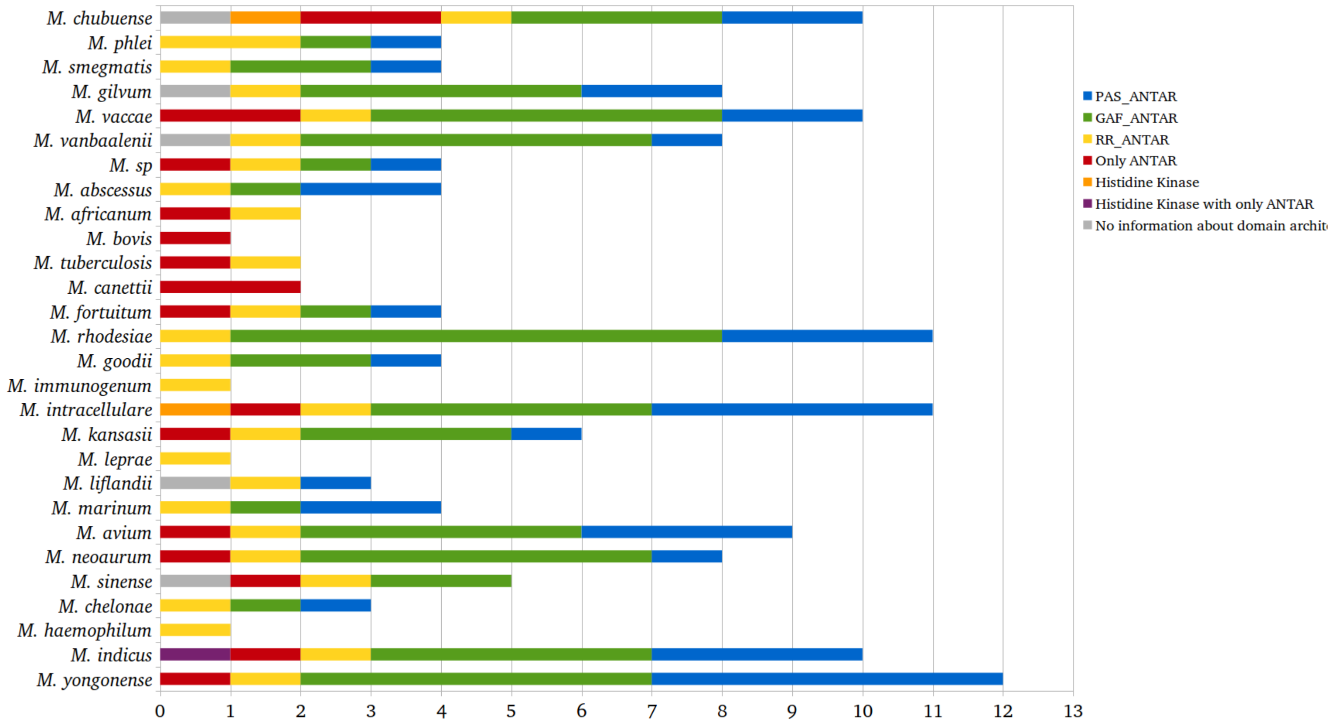


Figure S1. *Mycobacterial species contain ANTAR-proteins which associates with PAS and GAF domains.* A) Bar-plot showing number of ANTAR-proteins in different mycobacterial species along with its domain architecture. To identify ANTAR-proteins, HMM search is carried out in 128 mycobacterial strains using ANTAR-domain profile from Pfam v31.0 (PF03861) with e-value threshold 1e-4. Domain architecture information was obtained from Pfam and NCBI gene annotations.

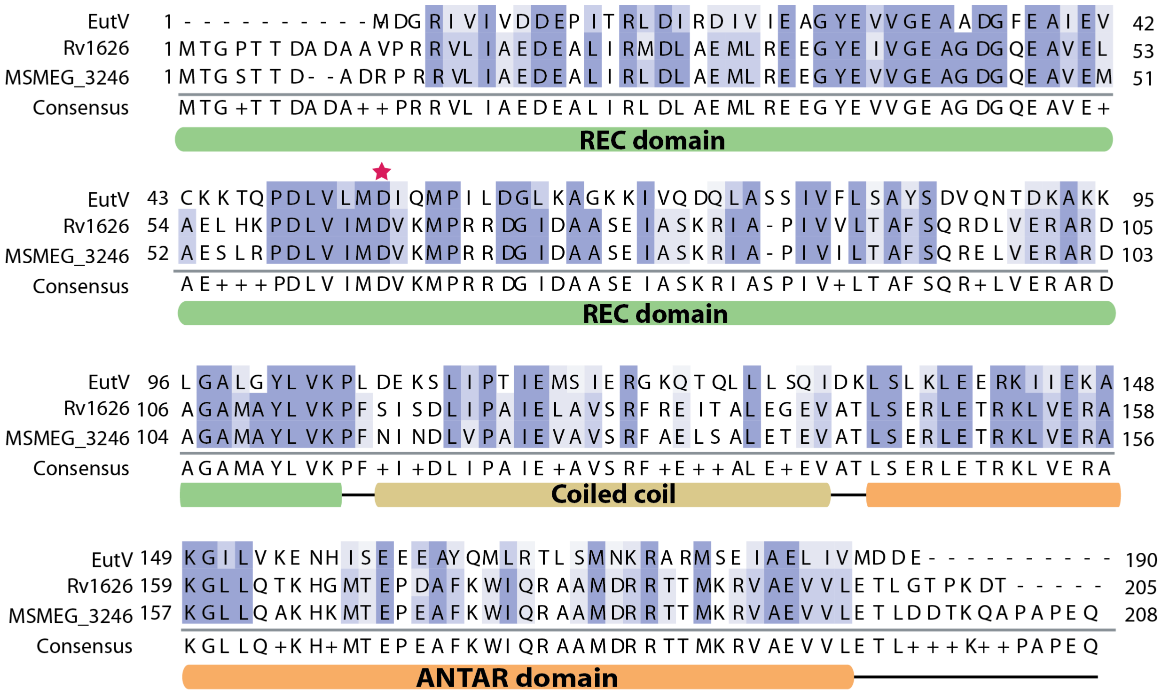

Figure S2. *ANTAR-proteins are conserved in Mycobacteria and Enterococcus sp.*
A) Multiple sequence alignment of ANTAR proteins from *Enterococcus faecalis (EutV)*, *Mycobacterium tuberculosis H37Rv* (Rv1626) and *Mycobacterium smegmatis MC2-155* (MSMEG_3246) is obtained using Clustal Omega v1.2.4 and visualized using JalView v2.10.5. Domain organization of Rv1626 (1S8N) is shown below the alignment. Aspartate residue that undergoes phosphorylation is marked (red asterisk). Mycobacterial proteins are >90% similar to each other while >65% similar to EutV.

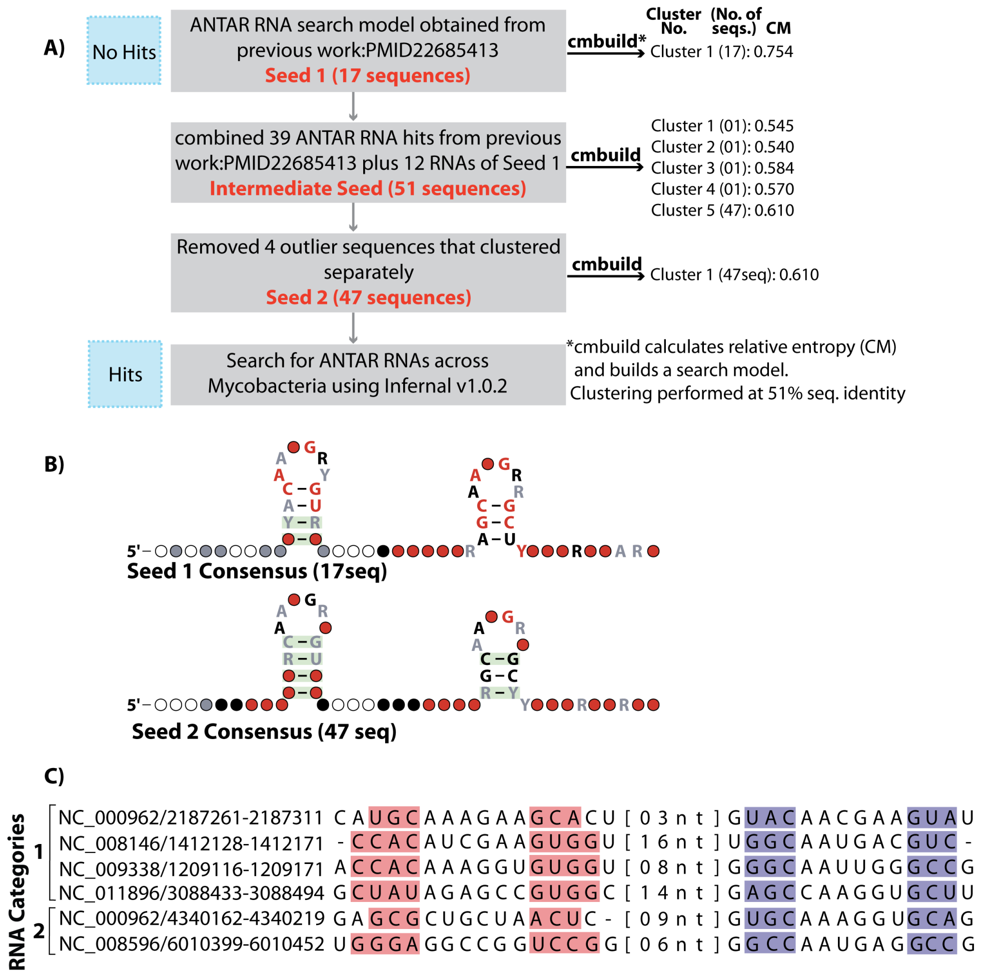


Figure S3. *Mycobacterial ANTAR RNA shows variations in structured motif.*
A) Flowchart represents the steps followed to computationally predict ANTAR-target RNAs in mycobacteria. 17 sequences used as search model (Seed 1) in previous work(4) were subjected to cmbuild analysis. Asterisk (*) marks the details of the cmbuild analysis. All sequences clustered with high CM score of 0.75. This search failed to identify RNAs in mycobacteria. We then added diverse ANTAR-target RNA hits from previous work to make an intermediate seed of 51 sequences, which was again analyzed using cmbuild. 47 sequences clustered together (Cluster 5) with CM score of 0.61, marking the other 4 as outliers. These 47 sequences were used as Seed 2, and used to search for RNAs in 128 sequenced mycobacterial genomes. This seed resulted in 222 RNA hits across mycobacterial species. B) Consensus structure for Seed 1 (previous work) and Seed 2 (this work) is shown. Co-varying positions are highlighted (green) and nucleotide identity above 97% (red), 90% (black) and 75%(grey) are marked. Seed 2 shows higher covariation within the two base paired stems. C) Representative RNA sequences from each category are shown along with accession number and RNA coordinates. Shaded regions highlight base pairing in stem 1 (pink) and stem 2 (blue).

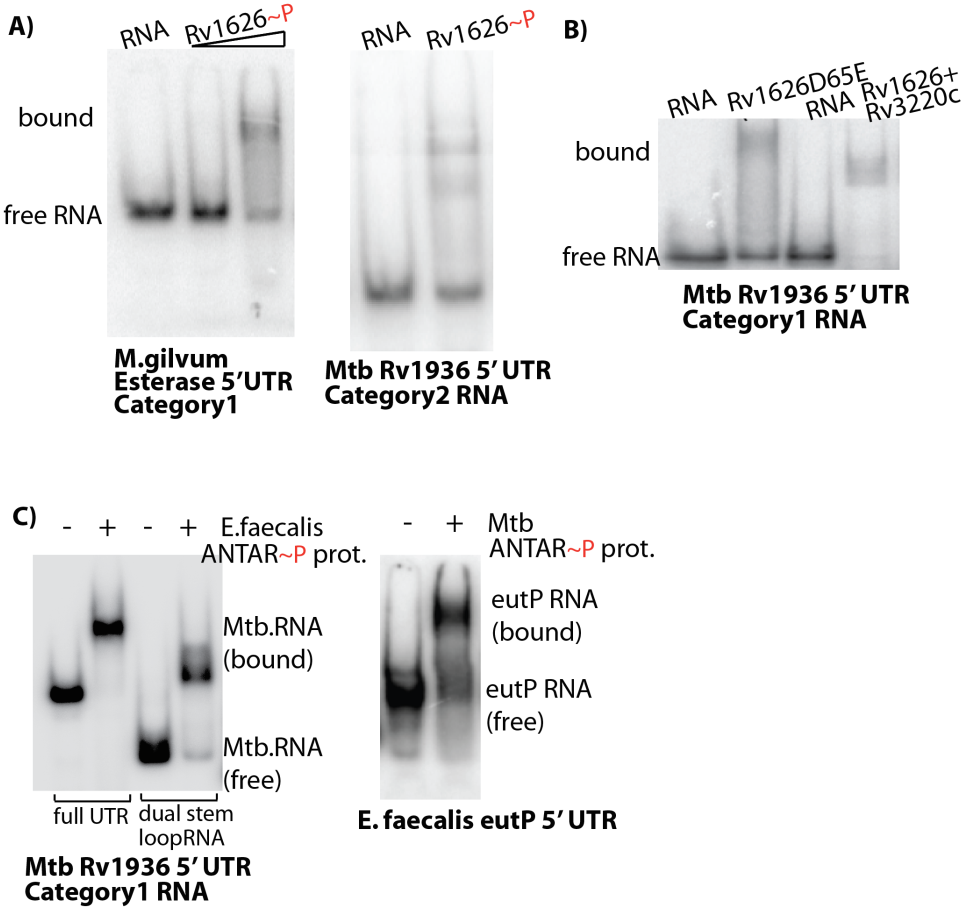


Figure S4. *Interaction between ANTAR RNA candidates with different ANTAR proteins*
A) EMSA reports on binding of Category 1 and 2 RNA candidates to Mtb ANTAR protein Rv1626D65E on a native gel. B) EMSA comparing binding of Mtb RNA (Rv1936 5' UTR) by ANTAR when phosphorylated by the kinase (Rv1626+Rv3220c) versus as a phosphomimic mutation (Rv1626D65E). No significant difference is observed between the two. C) EMSA shows cross recognition between ANTAR proteins and RNAs from different species. Mtb RNA (Rv1936 5' UTR) binds EutV protein from *E. faecalis* (left) and *E. faecalis* RNA (eutP 5' UTR) binds Rv1626 from Mtb (right).

**A)** ***Description of genes linked to predicted ANTAR RNAs***

*(Continued from Figure 5)*

***Secreted***

Rv3864-78 Esx-I secretion system and virulence
MJLS_RS12880 ATPase AAA
Rv2875 major secreted immunogenic protein Mpt70

***Membrane proteins***

Rv0872c PE-PGRS family protein PE-PGRS15
Rv3104c-03c Mechanosensitive ion channel protein MscS
ABG82_RS23295 amino acid permease

***Regulators***

MASS_RS17330 transcriptional regulator
MAH_RS12745 TetR family transcriptional regulator
MAB_4498c Putative two-component system response regulator, LuxR family
ABG82_RS25935 GntR family transcriptional regulator

***DNA- process***

MYCRHN_RS03655 DNA-directed RNA pol beta
MSPYR1_RS20440 transposase
MSMEG_5451 DNA-N1-methyladenine dioxygenase
MIP_RS11885 DNA-directed RNA polymerase sigma-70 factor

***Enzymes***

MYCCH_RS09880 glutamine synthetase
MSMEG_5948 glycosyl transferase family protein
MJLS_RS06690 glycosyl transferase

***Protein modification***

G155_RS30190 16S rRNA methyltransferase G

***Bacteriophage***

MAB_1787Bacteriophage protein

**B)**

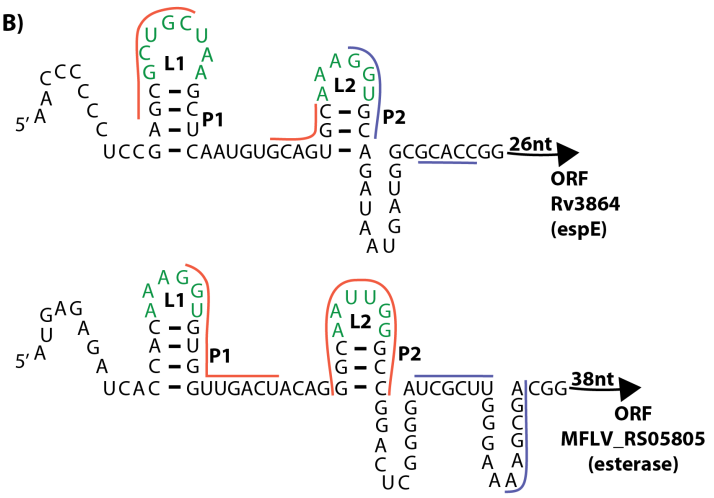


Figure S5. *Alternate base pairing within ANTAR RNAs*
A). Description of genes directly downstream of predicted ANTAR RNAs. Genes from the top 3 categories are shown in Figure 5A. Genes belonging to the remaining categories are shown here. B) Schematic showing predicted ANTAR RNAs upstream of the Mtb Rv3864 (espE) and *M. gilvum* PYR-GCK MFLV_RS05805 genes. Dual stem loops of ANTAR RNAs with base paried region (dashed line) and loops (green) are shown. ORF start positions (marked in bold) are obtained from NCBI. Alternate structure predicted using RNAfold server is indicated (Red, Blue lines, indicative of base-pairing). Based on this, ANTAR -binding would prevent the formation of a mutually exclusive structure.

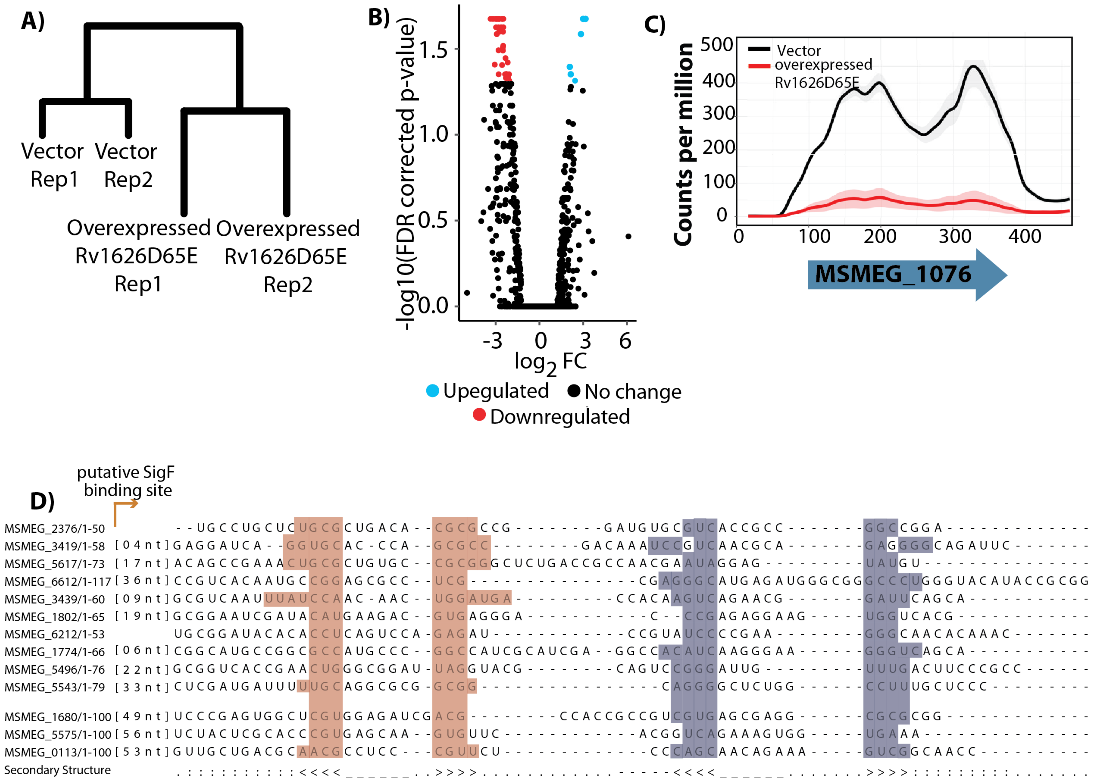


E)

***Nitrogen metabolism***

MSMEG_0133 transporter, nitrogen/carbon utilization
MSMEG_4511 isomerase- Mycobactin synthesis
MSMEG_5575 repressor for dNTP/NTP breakdown
MSMEG_1677-80 Asp/Ala metabolism, nitrogen cycle, MurNAC alanine amidase, DUF1028

***Redox***

MSMEG_6768 e- transfering flavoprotein dehydrogenase
MSMEG_0686 Fe-S cluster, e- transport chain protein
MSMEG_1076-77 oxidoreductase, acts on nitroimidazoles
MSMEG_1794 oxidoreductase on CHNH donors, NO2 stress
MSMEG_1132 FAD-linked oxidoreductase
MSMEG_5189 NAD(P) using dehydratase for dTDP-glucose
MSMEG_6212 Fe/Cd binding
MSMEG_6213 nitric oxide response, oxygen stress, peroxidase

***Membrane proteins/transport***

MSMEG_1131 integral membrane protein
MSMEG_5496 osmolarity control
MSMEG_6466 osmolarity control
MSMEG_3254-55 Li/Na/K antiporter

***Regulator***

MSMEG_1802 ChaB regulator of ion transport

Figure S6. *Overexpression of ANTAR~P in M. smegmatis alters the transcriptome.* A) Unsupervised hierarchical clustering using normalized CPM values indicates clustering of replicates of empty vector control and replicates of ANTAR~P overexpressing cells. B) Volcano plot of edgeR analysis shows significant expression changes with > 2-fold downregulated (red) and upregulated (blue) genes. C) Representative coverage plot of CPM versus nucleotide position, for a differentially expressed *M. smegmatis* gene- MSMEG_1076 shows downregulation under ANTAR~P conditions compared to empty vector control. D) Manually identified dual stem motifs in 100nt upstream region of 13 differentially expressed genes. These do not match ANTAR RNA consensus. The two stems are shaded in pink and blue and putative sigF binding site is highlighted. E) Detailed annotations of genes/operons in the different functional categories from panel A are shown.

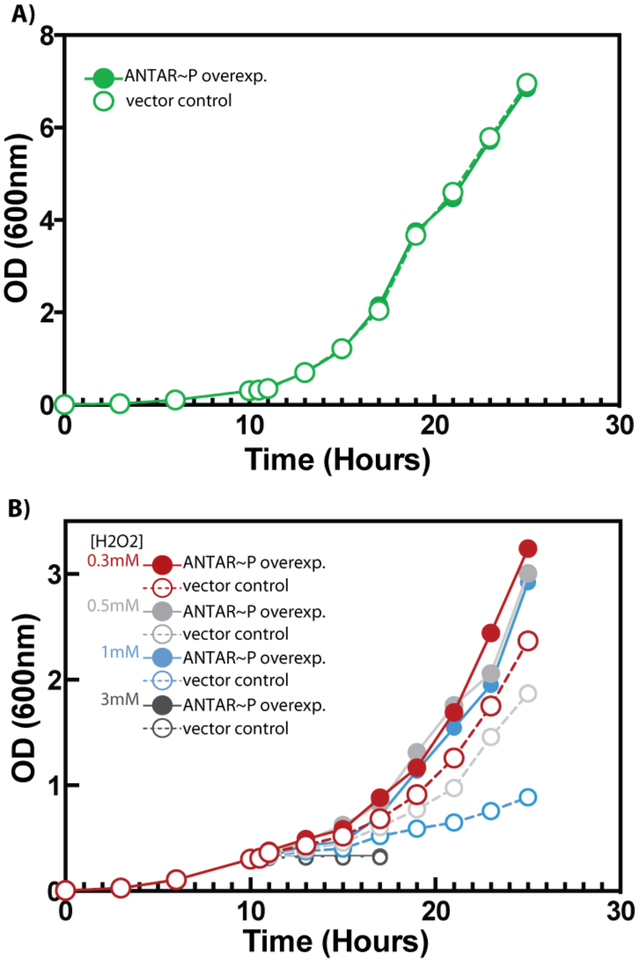


Figure S7. *ANTAR~P overexpression confers a growth advantage in oxidative stress.* A) Growth curves for ANTAR~P overexpression (solid) and vector control cells (dashed) in rich media, indicate identical growth rates. B) Growth curves for ANTAR~P overexpression (solid) and vector control cells (dashed) grown in indicated concentrations of H2O2 induced oxidative stress. At H2O2 concentrations ranging from 0.3mM to 1mM, cells with ANTAR~P overexpression show a distinct growth advantage.

*Table S1: Covariance search seed for ANTAR RNA dual stemloop motif*

Sequences from diverse bacteria are used to make the seed where base paired stems are shaded in pink and blue. Twelve sequences from the Seed 1 shown in grey are included along with 35 new sequences (black) to assemble the seed (Seed 2). Secondary structure of the RNA is shown in WUSS notation where '<' and '>' indicates base-pairing positions.

| Accession/Start-End | Sequence |
|---|---|
| NC_009089.1/2209707_2209768/1-62 | --UAUAAUGAAAGUAUCACAAAGGCGUGAUAC----------------UUUAGUAGAAAGCAAUGGAGCUUCUUAGGAAU |
| NC_004668.1/1589608_1589547/1-62 | --UAUAAUGUGGGUAGCACAACGGCGUGCUUC----------------AAAAUUUAAGAGCAAAGAAGCUCCUUAGUAGA |
| NC_009089.1/2212731_2212788/1-58 | --AAUAAUAACUCAUCUACAAAGUAGUAGUU--------------------UUAAUUGAGCAAAGAGGCUUAAGGUUAUU |
| NC_004668.1/1592374_1592320/1-55 | -------AAUCAGAAACACAAUGGCGUGUUUU------------------AACAAAUCGGCAAAGGAGCCCAAGACUAAG |
| NC_004557.1/2307829_2307768/1-62 | --AAUAUUAAUAAUUAUACAAAGGCGUAUAAU----------------UAUCGGUAAGAGCAAUGGAGCUCCUUAAAAUA |
| NC_008555.1/1141572_1141629/1-58 | --AAAAUAUUUUACAGUACAAAGGCGUACUG--------------------UUUACUUAGCAAAGAAGCUUUGAGUUAGA |
| NC_003210.1/1202056_1202113/1-58 | --AAUUCUAUCCAUGCUACAAAGAAGUAGC--------------------UAUGAAAAAGCGAUGAAGCUUAAAGCCAAG |
| NC_008261.1/1052251_1052307/1-57 | -----AAUAAAGAUAUCACAAAGGUGUGAUUA------------------AGUUUUCGAGCAAUGUAGCUUCUUAAAGGU |
| NC_003210.1/1198518_1198575/1-58 | --AAAAUAUUUUACGGUACAAAGGCGUACUG--------------------UUUACUUAGCAAAGAAGCUUUGAGUUGGA |
| NC_003212.1/1151140_1151197/1-58 | --AAAAUAUUUUACGGUACAAAGGCGUACUG--------------------UUUACUUAGCAAAGAAGCUUUGAGUUGGA |
| NC_003212.1/1154678_1154736/1-59 | --AAUUCAAUCCAUGCUACAAAGAAGUAGCU-------------------AUGAAAAAAGCGAUGAAGCUUAAAGCCAAG |
| NC_008555.1/1145109_1145166/1-58 | --AUUUCAAUCCAUGCUACAAAGAAGUAGC--------------------UAUGAAAAAGCGAUGAAGCUUAAAGCCAAG |
| NC_009633.1/290271_290331/1-61 | -AAUUUCAAACAAUUGCACAAUGGCGUGCAAU------------------GAUUAAAAAGCGAUGAAGCUUGCAGGAAGG |
| NC_009009.1/512876_512934/1-59 | ----AAUUCAAUAUAGUACAAAGGGGUACUGA-----------------AGAUAAAUAAGCAAUGAAGCUAUGUGAAAAU |
| NC_004557.1/2309190_2309131/1-60 | -UAUAAAAUGAAAUUGCACAAUGAUGUGCAA-------------------UAGUAGAGAGUAAUGAAGCUCCUAAAGAUU |
| NC_009009.1/466893_466954/1-62 | -------GAAAACGAACACAACGGAGUGUUUG-----------AACUACAUAUAAGUGAGCAACGGUGCUACAAAAAAUG |
| NC_008261.1/1050699_1050753/1-55 | --UAUAUUAAAAAUAACACGAUGUGGUGU-----------------------UUAGAGAGCAAAGAAGCUCCAUAGGAAA |
| NC_009633.1/293287_293341/1-55 | ----AAUAAAAUCAAGCACAAUGGUGUGUU---------------------UGUUCAAGGCAAUGGGGCCAAGGAUCUAU |
| NC_003454.1/716227_716290/1-64 | --AAUUAAAUAAGAUAUACAAUGGAGUAUAUU--------------AUAAAAAUAGAAAGCAAUGAAGCUGAUGGACAGG |
| NC_007498_2_2447483_2447533/1-51 | ---------CACAUGACACAACGGCGUGUC--------------------AGGCUUACGGCAACGACGCCCCAUUCAGGA |
| NC_009253.1/3046896_3046842/1-55 | --AACAACAUACUCUGAACAACGAUGUUCU-----------------------AGUCAGGCAAUGAUGCCUUUUGUAAGA |
| NC_002516_2_1855754_1855810/1-57 | -------GGAAAAGGCCACAGCGAUGUGGCUU----------------UUUUCUUAAAAGAAAAGUCUCUCAGUGACAAA |
| NC_007498_2_2456540_2456594/1-55 | ---------AACGGGGCACAAUGGCGUGCCUG----------------GCUGAGAAACGGCAACGAGGCCCGGUGGGAUU |
| NC_009654.1/3196287_3196231/1-57 | -----AUUAUUGAUUGUCCAAUGUAGGAUUUU------------------CUUUGAAUGGCAAAGGCGCCUGCGAACACA |
| NC_009253.1/3045296_3045244/1-53 | ------AAUUGAAAAGCACGAUGAUGUGCU---------------------AACGAAUGGCAAUGAAGCCUACAGCGGGU |
| NC_008726.1/6033961_6033904/1-58 | ----AAGAGAGAUCACCACAAAGGUGUGGUUG------------------ACUACACGGGCAAAGUGGCCGGUUUUCGGG |
| NC_003197.1/3479948_3480000/1-53 | --------UAUACCUGAAGAAAGGACUUUAG-------------------AUGCUUUUAGCGAUGGCGCUACUCAUAAUU |
| NC_007907.1/5011991_5011936/1-56 | ---------UGUUACUUUUAAUGCAGAAAUUU---------------UAUAAAUGAAGAGCAAAGGCGCUCAUAAGGAAU |
| NC_007492_2_3807050_3807001/1-50 | ---------AUCGAUGCGCAAAGGCGUGCU---------------------AUGGAAACUCAAGGGAGGGUUAAAGCAUG |
| NC_008228.1/836271_836321/1-51 | -------AAUUUGGAGCAUAAGGAUAUGUUUC----------------------GUCUUGCAAUGGAGCAUUCAGCCAUA |
| NC_009485.1/6902858_6902810/1-49 | ---------AUAUCGUCGCAAGGAUGCGG----------------------AGUGAAGGGCAAUGAAGCUCUUAGAUAAU |
| NC_008726.1/4556568_4556520/1-49 | ---------UAUAGUGCACAUGGAUGUGCG---------------------AUAAAUGAGCAAAGU-GCUUACAAAAUAC |
| NC_009972.1/5354031_5353982/1-50 | ----UAUAAUGCCCGCAGCAACGACGCUGA--------------------------UGAGCAAUGGUGCUAAUUCACAUA |
| NC_009922.1/35107_35156/1-50 | --------AAUCUAUGAAAAAUAGUUUUU----------------------CAAAUUAAGCAAAGGAGCUCAGAUAGAUG |
| NC_009792.1/4243606_4243658/1-53 | --------UAUACCUGAAGAAAGGACUUUAG-------------------AUGCUUUUAGCUACGGCGCUGUUAAUUAUU |
| NC_009972.1/2941344_2941396/1-53 | ----------AAAUGCAACGGAGCCGUUGUAU-----------------UUUGGAAUGCACAAAGGAGUGCCAUAAUGCU |
| NC_007643.1/2914531_2914583/1-53 | -----------AUGGCGGCAAAGGCGCCUGA----------------CCUCAUGAUGCGGCAAAGGCGCCCGAGCUAAAA |
| NC_006350.1/1450675_1450740/1-66 | --------ACUAUCAAUACAAAGAAGUAUUCG------AUUGACAUGAAUCAACGCAAGGCAACGGCGCUCAUGAGAUUU |
| NC_000913_2_1424328_1424385/1-58 | ---------ACUCUGUUACAGAGAUGUGAUGG-------------UGUCUUUAAUUAAAUCGAUGAUGCUCCUGGAGAAA |
| NC_009922.1/1019131_1019178/1-48 | ------------AUGGACCAAUGUCGUUUG---------------------AGGCCAUAGAAAAGACUUUUAAAAAAGGG |
| NC_008255.1/3523741_3523812/1-72 | ----AAUUUUAUUUCUGUAAAAGACGUUACUG----AUAUUCCCGGCCUGGUUAAAGAAGCAUUGGAGCUUAAAAAAAAU |
| NC_007086.1/3508976_3509026/1-51 | ------------GGUCCACAAUGGCGAGCCGC------------------CCAUUAUUUGCAAUGCAGCAUCUGGCGAAA |
| NC_007651.1/3312922_3312859/1-64 | ----------UAUCAACACAAAGAAGUAUUCG------AUUGACAUGAAUCAACGCAAAGCAACGGUGCUCAUGAGAUUU |
| NC_008255.1/1241209_1241266/1-58 | ----AAUAAUAAUGACGCCAAAGAUUUGUAU------------------UAUAUGUGCGCAAAGGCGCUACGUUUCAGG |
| NC_003198.1/2954650_2954705/1-56 | -------AAUUUGAAUACCAAUAAUGUUAUUU-------------------AACAAAAGGCAUUGAUGCUCUGGCCAGAA |
| NC_006177.1/1341787_1341840/1-54 | -------UGAAAUUACCAGAAAGAAAUGGUCG-------------------CUGAUUUCGCAAGAAAGCGCCGAAUUAAG |
| NC_010001.1/4467323_4467260/1-64 | ---UAUAUUGUAUAAGGAUAAUGGAAUGCCCU-------------CAUUUUUAUUCAAUGCAAGGAAGCACUUUGCGUAA |
| RF | ..................cAa.G..g..................................gcAa.G..gc........... |
| SS_cons | ...............<<<<......>>>>..............................<<<......>>>.......... |

Table S2: *Mycobacterial ANTAR RNAs identified through covariance search*
ANTAR RNAs identified using Infernal v1.0.2 are shown for each mycobacterial strain given with the genome accession ID. Bit score is the value assigned by the covariance search to indicate similarity to the search model. A higher score suggests that the predicted RNA is more similar to the search model. Arbitrary bit score threshold of 10.00 is used in this study to reduce occurrence of false-positives. RNA co-ordinates are as predicted by Infernal. Putative ANTAR RNA candidates are categorized based on structure similarity to the search model as shown in Figure 3 and Figure S3. Immediate gene in *cis* to the predicted RNA is indicated as NCBI Gene annotations.

| Organism | Bit Score | ANTAR RNA hit co-ordinates | Strand | RNA Category | Annotation of the downstream gene (Locus Tag) |
|---|---|---|---|---|---|
| *Mycobacterium_chubuense_NBB4* (NC_018027.1) | 11.2 | 2095742-2095811 | 1 | 2 | glutamine synthetase (MYCCH_RS09880) |
| *Mycobacterium_phlei_strain=CCUG21000* (NZ_CP014475.1) | 12.92 | 1931198-1931132 | -1 | 1 | hypothetical protein (MPHLCCUG_RS09165) |
| *Mycobacterium_smegmatis_str_MC2_155_1* (NC_008596.1) | 11.04 | 3682246-3682309 | 1 | 1 | short chain dehydrogenase (MSMEG_3619) |
| | 10.77 | 6497108-6497173 | 1 | 2 | hypothetical protein (MSMEG_6428) |
| | 10.75 | 6010452-6010399 | -1 | 2 | glycosyl transferase family protein (MSMEG_5948) |
| | 10.55 | 5536087-5536176 | 1 | 2 | alkylated DNA repair protein (MSMEG_5451) |
| *Mycobacterium_smegmatis_str_MC2_155_2* (NC_018289.1) | 11.04 | 3683859-3683922 | 1 | 1 | short-chain dehydrogenase (MSMEI_RS17805) |
| | 10.77 | 6498717-6498782 | 1 | 2 | hypothetical protein (MSMEI_RS31480) |
| | 10.75 | 6012063-6012010 | -1 | 2 | glycosyl transferase (MSMEI_RS29110) |
| | 10.55 | 5537699-5537788 | 1 | 2 | DNA-N1-methyladenine dioxygenase (MSMEI_RS26655) |
| *Mycobacterium_smegmatis_str_MC2_155_3* (NZ_CP009494.1) | 11.04 | 3682286-3682349 | 1 | 1 | short-chain dehydrogenase (LJ00_RS17790) |
| | 10.77 | 6497148-6497213 | 1 | 2 | hypothetical protein (LJ00_RS31460) |
| | 10.75 | 6010495-6010442 | -1 | 2 | glycosyl transferase (LJ00_RS29095) |
| | 10.55 | 5536133-5536222 | 1 | 2 | DNA-N1-methyladenine dioxygenase (LJ00_RS26635) |
| *Mycobacterium_smegmatis_strain=INHR1* (NZ_CP009495.1) | 11.04 | 3682320-3682383 | 1 | 1 | short-chain dehydrogenase (LI99_RS17790) |
| | 10.77 | 6497210-6497275 | 1 | 2 | hypothetical protein (LI99_RS31465) |
| | 10.75 | 6010565-6010512 | -1 | 2 | glycosyl transferase (LI99_RS29100) |
| | 10.55 | 5536187-5536276 | 1 | 2 | DNA-N1-methyladenine dioxygenase (LI99_RS26640) |
| *Mycobacterium_smegmatis_strain=INHR2* (NZ_CP009496.1) | 11.04 | 3682308-3682371 | 1 | 1 | short-chain dehydrogenase (LI98_RS17785) |
| | 10.77 | 6497179-6497244 | 1 | 2 | hypothetical protein (LI98_RS31465) |
| | 10.75 | 6010533-6010480 | -1 | 2 | glycosyl transferase (LI98_RS29100) |
| | 10.55 | 5536168-5536257 | 1 | 2 | DNA-N1-methyladenine dioxygenase (LI98_RS26640) |
| *Mycobacterium_smegmatis_strain=NCTC8159_NCTC8159* (NZ_LN831039.1) | 11.04 | 3744254-3744317 | 1 | 1 | short-chain dehydrogenase (AT701_RS18055) |
| | 10.77 | 6514456-6514521 | 1 | 2 | hypothetical protein (AT701_RS31495) |
| | 10.55 | 5543050-5543139 | 1 | 2 | DNA-N1-methyladenine dioxygenase (AT701_RS26625) |
| *Mycobacterium_gilvum_PYR-GCK* (NC_009338.1) | 18.51 | 1209116-1209171 | 1 | 1 | esterase/ diacyl-glycerol transferase (MFLV_RS05805) |
| | 11.26 | 2744062-2744015 | -1 | 1 | FAD-binding monooxygenase (MFLV_RS13185) |
| *Mycobacterium_gilvum_Spyr1* (NC_014814.1) | 18.51 | 5213367-5213312 | -1 | 1 | esterase/ diacyl-glycerol transferase (MSPYR1_RS24685) |
| | 11.26 | 2104189-2104142 | -1 | 1 | FAD-binding monooxygenase (MSPYR1_RS10115) |
| | 11.83 | 4314715-4314648 | -1 | 2 | transposase (MSPYR1_RS20440) |

| Organism | Bit Score | ANTAR RNA hit co-ordinates | Strand | RNA Category | Annotation of the downstream gene (Locus Tag) |
|---|---|---|---|---|---|
| *Mycobacterium_vaccae_95051* (NZ_CP011491.1) | 22.91 | 5794907-5794851 | -1 | 1 | esterase/ diacyl-glycerol transferase (MYVA_RS27135) |
| | 11.84 | 3023516-3023552 | 1 | 1 | Carboxyl esterase (MYVA_RS14175) |
| | 13.83 | 1708870-1708817 | -1 | 2 | hypothetical protein (MYVA_RS08015) |
| *Mycobacterium_vanbaalenii_PYR-1* (NC_008726.1) | 28.36 | 6033960-6033904 | -1 | 1 | esterase/ diacyl-glycerol transferase (MVAN_RS28400) |
| | 21.41 | 4556572-4556520 | -1 | 1 | alpha/beta hydrolase fold protein (MVAN_RS21295) |
| *Mycobacterium_sp_EPa45* (NZ_CP011773.1) | 15.09 | 2186394-2186461 | 1 | 1 | cytochrome P450 (AB431_RS10365) |
| *Mycobacterium_sp_JLS* (NC_009077.1) | 12.42 | 2700344-2700396 | 1 | 1 | ATPase AAA (MJLS_RS12880) |
| | 10.42 | 1434237-1434280 | 1 | 1 | glycosyl transferase (MJLS_RS06690) |
| *Mycobacterium_sp_JS623* (NC_019958.1) | 12.38 | 29809-29757 | -1 | 1 | hypothetical protein (MYCSM_RS33235) |
| *Mycobacterium_sp_JS623* (NC_019966.1) | 10.71 | 3367677-3367739 | 1 | 2 | hypothetical protein (MYCSM_RS16585) |
| *Mycobacterium_sp_KMS* (NC_008705.1) | 12.42 | 2714603-2714655 | 1 | 1 | ATPase AAA (MKMS_RS12840) |
| | 10.42 | 1416437-1416480 | 1 | 1 | glycosyl transferase (MKMS_RS06575) |
| *Mycobacterium_sp_MCS* (NC_008146.1) | 12.42 | 2696715-2696767 | 1 | 1 | ATPase AAA (MMCS_RS12745) |
| | 10.42 | 1412128-1412171 | 1 | 1 | glycosyl transferase (MMCS_RS06550) |
| *Mycobacterium_sp_NRRL_B-3805* (NZ_CP011022.1) | 14.17 | 4181243-4181304 | 1 | 1 | hypothetical protein (MyAD_RS19495) |
| | 10.32 | 1555180-1555232 | 1 | 2 | hypothetical protein (MyAD_RS07380) |
| *Mycobacterium_sp_QIA-3* (NZ_CP010071.1) | 11.18 | 3306365-3306418 | 1 | 2 | cytochrome P450 (Chelonae_RS16150) |
| *Mycobacterium_sp_VKM_Ac-1817D* (NZ_CP009914.1) | 14.91 | 6321182-6321128 | -1 | 1 | 16S rRNA methyltransferase G (G155_RS30185) |
| | 11.05 | 2975165-2975230 | 1 | 1 | polyketide synthase (G155_RS14365) |
| | 10.22 | 5424084-5424140 | 1 | 1 | hypothetical protein (G155_RS25930) |
| *Mycobacterium_abscessus_ATCC_19977* (NC_010397.1)) | 14.49 | 4098501-4098456 | -1 | 1 | reductase (MAB_4051c) |
| | 13.68 | 1789584-1789640 | 1 | 1 | Bacteriophage protein (MAB_1787) |
| | 10.24 | 4575249-4575201 | -1 | 2 | Putative two-component system response regulator, LuxR family (MAB_4498c) |
| *Mycobacterium_abscessus_strain=FLAC003* (NZ_CP014950.1) | 10.42 | 3909717-3909764 | | 1 | reductase (A3N95_RS19335) |
| | 11.53 | 4379382-4379334 | -1 | 2 | DNA-binding response regulator (A3N95_RS21510) |
| | 10.25 | 3287342-3287395 | 1 | 2 | cytochrome P450 (A3N95_RS16215) |
| *Mycobacterium_abscessus_strain=FLAC004* (NZ_CP014951.1) | 10.25 | 3721005-3721058 | 1 | 2 | cytochrome P450 (A3N96_RS18920) |
| | 10.19 | 4374182-4374135 | -1 | 2 | reductase (A3N96_RS22325) |
| *Mycobacterium_abscessus_strain=FLAC005* (NZ_CP014952.1) | 14.49 | 3992094-3992049 | -1 | 1 | reductase (A3N97_RS19695) |
| | 10.25 | 3364829-3364882 | 1 | 2 | cytochrome P450 (A3N97_RS16545) |
| | 10.24 | 4425574-4425526 | -1 | 2 | DNA-binding response regulator (A3N97_RS21715) |
| *Mycobacterium_abscessus_strain=FLAC007* (NZ_CP014953.1) | 10.25 | 3586037-3586090 | 1 | 2 | cytochrome P450 (A3N98_RS18115) |
| | 10.19 | 4197314-4197267 | -1 | 2 | reductase (A3N98_RS21205) |
| *Mycobacterium_abscessus_strain=FLAC008* (NZ_CP014954.1) | 10.25 | 3686526-3686579 | 1 | 2 | cytochrome P450 (A3N99_RS18710) |
| | 10.19 | 4297803-4297756 | -1 | 2 | reductase (A3N99_RS21800) |
| *Mycobacterium_abscessus_strain=FLAC013* (NZ_CP014955.1) | 14.49 | 4105821-4105776 | -1 | 1 | reductase (A3O00_RS20355) |
| | 13.68 | 1772773-1772829 | 1 | 1 | hypothetical protein (A3O00_RS08870) |

| Organism | Bit Score | ANTAR RNA hit co-ordinates | Strand | RNA Category | Annotation of the downstream gene (Locus Tag) |
|---|---|---|---|---|---|
| | 10.24 | 4582443-4582395 | -1 | 2 | DNA-binding response regulator (A3O00_RS22565) |
| *Mycobacterium_abscessus_strain=FLAC029* (NZ_CP014956.1) | 14.49 | 4220810-4220765 | -1 | 1 | reductase (A3O01_RS21075) |
| | 10.24 | 4697506-4697458 | -1 | 2 | DNA-binding response regulator (A3O01_RS23280) |
| *Mycobacterium_abscessus_strain=FLAC031* (NZ_CP014957.1) | 14.49 | 4178235-4178190 | -1 | 1 | reductase (A3O02_RS20770) |
| | 10.25 | 3490346-3490399 | 1 | 2 | cytochrome P450 (A3O02_RS17205) |
| *Mycobacterium_abscessus_strain=FLAC045* (NZ_CP014958.1) | 10.25 | 3693121-3693174 | 1 | 2 | cytochrome P450 (A3O03_RS18825) |
| | 10.19 | 4304398-4304351 | -1 | 2 | reductase (A3O03_RS21915) |
| *Mycobacterium_abscessus_strain=FLAC048* (NZ_CP014959.1) | 10.42 | 4078670-4078623 | -1 | 1 | reductase (A3O04_RS20205) |
| | 11.72 | 4510794-4510746 | -1 | 2 | DNA-binding response regulator (A3O04_RS22210) |
| | 10.25 | 3453708-3453761 | 1 | 2 | cytochrome P450 (A3O04_RS17045) |
| *Mycobacterium_abscessus_strain=FLAC049* (NZ_CP014960.1) | 14.49 | 3952609-3952564 | -1 | 1 | reductase (A3O05_RS19490) |
| | 10.24 | 4392665-4392617 | -1 | 2 | DNA-binding response regulator (A3O05_RS21530) |
| *Mycobacterium_abscessus_strain=FLAC054* (NZ_CP014961.1) | 10.42 | 4460783-4460830 | -1 | 1 | reductase (A3O06_RS22720) |
| | 11.72 | 4891352-4891304 | -1 | 2 | DNA-binding response regulator (A3O06_RS24710) |
| | 10.25 | 3791427-3791480 | 1 | 2 | cytochrome P450 (A3O06_RS19230) |
| *Mycobacterium_abscessus_strain=NOV0213* (NZ_CP013049.1) | 14.49 | 4103527-4103482 | -1 | 1 | reductase (AOY11_RS20460) |
| | 10.24 | 4583475-4583427 | -1 | 2 | DNA-binding response regulator (AOY11_RS22690) |
| *Mycobacterium_abscessus_subsp_Abscessus_1* (NZ_CP009615.1) | 10.25 | 1450868-1450815 | -1 | 2 | cytochrome P450 (NF92_RS07030) |
| | 10.19 | 851498-851545 | 1 | 2 | reductase (NF92_RS03990) |
| *Mycobacterium_abscessus_subsp_Abscessus_2* (NZ_CP009616.1) | 10.25 | 1451017-1450964 | -1 | 2 | cytochrome P450 (NF90_RS07030) |
| | 10.19 | 851598-851645 | 1 | 2 | reductase (NF90_RS03990) |
| *Mycobacterium_abscessus_subsp_bolletii_103* (NZ_CP009407.1) | 14.49 | 4082796-4082751 | -1 | 1 | reductase (LA61_RS20280) |
| | 13.68 | 1774108-1774164 | 1 | 1 | hypothetical protein (LA61_RS08890) |
| | 10.24 | 4559548-4559500 | -1 | 2 | DNA-binding response regulator (LA61_RS22490) |
| *Mycobacterium_abscessus_subsp_bolletii_50594* (NC_021282.1) | 11.72 | 4535117-4535069 | -1 | 2 | DNA-binding response regulator (MASS_RS22920) |
| | 10.44 | 3437738-3437670 | -1 | 2 | transcriptional regulator (MASS_RS17330) |
| | 10.25 | 3403861-3403914 | 1 | 2 | cytochrome P450 (MASS_RS17145) |
| | 10.19 | 4061201-4061154 | -1 | 2 | reductase (MASS_RS20610) |
| *Mycobacterium_abscessus_subsp_bolletii_CCUG_48898_=_JCM_15300* (NZ_AP014547.1) | 11.72 | 4549416-4549368 | -1 | 2 | DNA-binding response regulator (MMASJCM_RS22535) |
| | 10.42 | 4115542-4115495 | -1 | 2 | reductase (MMASJCM_RS20510) |
| | 10.25 | 3435184-3435237 | 1 | 2 | cytochrome P450 (MMASJCM_RS16990) |
| *Mycobacterium_abscessus_subsp_bolletii_MA_1948* (NZ_CP009408.1) | 14.49 | 4097642-4097597 | -1 | 1 | reductase (LA62_RS20365) |
| | 13.68 | 1789073-1789129 | 1 | 1 | hypothetical protein (LA62_RS08980) |
| | 10.24 | 4572395-4572347 | -1 | 2 | DNA-binding response regulator (LA62_RS22580) |
| *Mycobacterium_abscessus_subsp_Bolletii_MC1518* (NZ_CP009613.1) | 14.49 | 4080706-4080661 | -1 | 1 | reductase (NF82_RS20305) |
| | 13.68 | 1781082-1781138 | 1 | 1 | hypothetical protein (NF82_RS08935) |
| | 10.24 | 4557425-4557377 | -1 | 2 | DNA-binding response regulator (NF82_RS22515) |

| Organism | Bit Score | ANTAR RNA hit co-ordinates | Strand | RNA Category | Annotation of the downstream gene (Locus Tag) |
|---|---|---|---|---|---|
| *Mycobacterium_abscessus_subsp_Bolletii_MM1513* (NZ_CP009447.1) | 11.72 | 405303-405351 | 1 | 2 | DNA-binding response regulator (LH56_RS01865) |
| | 10.25 | 1435695-1435642 | -1 | 2 | cytochrome P450 (LH56_RS06925) |
| | 10.19 | 837674-837721 | 1 | 2 | reductase (LH56_RS03890) |
| *Mycobacterium_abscessus_subsp_bolletii_str_GO_06* (NC_018150.2) | 10.25 | 1451040-1450987 | -1 | 2 | cytochrome P450 (MYCMA_RS07030) |
| | 10.19 | 851623-851670 | 1 | 2 | reductase (MYCMA_RS03990) |
| *Mycobacterium_abscessus_UC22* (NZ_CP012044.1) | 14.49 | 4394004-4393959 | -1 | 1 | reductase (MAUC22_RS22025) |
| | 10.24 | 4843707-4843659 | -1 | 2 | DNA-binding response regulator (MAUC22_RS24155) |
| *Mycobacterium_africanum_GM041182* (NC_015758.1) | 17.37 | 2187796-2187846 | 1 | 1 | monooxygenase (MAF_RS10120) |
| | 13.76 | 3192504-3192558 | 1 | 1 | hypothetical protein (MAF_RS15125) |
| | 11.95 | 966641-966587 | -1 | 1 | hypothetical protein (MAF_RS04600) |
| | 11.67 | 3452031-3451978 | -1 | 1 | hypothetical protein (MAF_RS16190) |
| | 10.14 | 3485225-3485278 | 1 | 1 | AmmeMemoRadiSam system radical SAM enzyme (MAF_RS16385) |
| | 15.35 | 4317847-4317919 | 1 | 2 | ESX-1 secretion-associated protein EspE (MAF_RS20165) |
| | 10.74 | 3164610-3164675 | 1 | 2 | cell surface protein (MAF_RS14980) |
| | 10.74 | 3501741-3501787 | 1 | 2 | NADH-quinone oxidoreductase subunit I (MAF_RS16460) |
| | 10.04 | 2780855-2780806 | -1 | 2 | carboxylesterase LipQ (MAF_RS13005) |
| *Mycobacterium_avium_104* (NC_008595.1) | 12.37 | 2096449-2096515 | 1 | 2 | hypothetical protein (MAV_RS10005) |
| *Mycobacterium_avium_subsp_avium* (NZ_CP009614.1) | 12.37 | 1991098-1991164 | 1 | 2 | hypothetical protein (NF84_RS09385) |
| *Mycobacterium_avium_subsp_avium_2285_R* (NZ_CP009493.1) | 12.37 | 1986373-1986439 | 1 | 2 | hypothetical protein (LA63_RS09380) |
| *Mycobacterium_avium_subsp_avium_2285_S* (NZ_CP009482.1) | 12.37 | 1985968-1986034 | 1 | 2 | hypothetical protein (LA64_RS09395) |
| *Mycobacterium_avium_subsp_hominissuis_TH135* (NZ_AP012555.1) | 11.05 | 2832453-2832394 | -1 | 1 | TetR family transcriptional regulator (MAH_RS12745) |
| | 12.37 | 1815297-1815363 | 1 | 2 | hypothetical protein (MAH_RS08425) |
| *Mycobacterium_avium_subsp_paratuberculosis_E1* (NZ_CP010113.1) | 12.37 | 1882988-1883054 | 1 | 2 | hypothetical protein (RC58_RS08685) |
| *Mycobacterium_avium_subsp_paratuberculosis_E93* (NZ_CP010114.1) | 12.37 | 1885164-1885230 | 1 | 2 | hypothetical protein (RE97_RS08685) |
| *Mycobacterium_avium_subsp_paratuberculosis_K-10* (NC_002944.2) | 12.37 | 2307090-2307024 | -1 | 2 | hypothetical protein (MAP_RS10605) |
| *Mycobacterium_avium_subsp_paratuberculosis_MAP4* (NC_021200.1) | 12.37 | 1902216-1902282 | 1 | 2 | hypothetical protein (MAP4_RS08705) |
| *Mycobacterium_avium_subsp_Paratuberculosis_strain=MAP_TANUVAS_TN_India2008* (NZ_CP015495.1) | 12.37 | 2307090-2307024 | -1 | 2 | hypothetical protein (MAP_RS10605) |
| *Mycobacterium_bovis_BCG_str_ATCC_35743* | 17.37 | 1889021-1889071 | 1 | 1 | monooxygenase (BCGT_RS08920) |

| Organism | Bit Score | ANTAR RNA hit co-ordinates | Strand | RNA Category | Annotation of the downstream gene (Locus Tag) |
|---|---|---|---|---|---|
| (NZ_CP003494.1) | 13.76 | 2878418-2878472 | 1 | 1 | hypothetical protein (BCGT_RS13865) |
| | 11.95 | 688333-688279 | -1 | 1 | hypothetical protein (BCGT_RS03405) |
| | 11.67 | 3137311-3137258 | -1 | 1 | hypothetical protein (BCGT_RS14930) |
| | 10.14 | 3167077-3167130 | 1 | 1 | AmmeMemoRadiSam system radical SAM enzyme (BCGT_RS15100) |
| | 15.35 | 3987382-3987454 | 1 | 2 | ESX-1 secretion-associated protein EspE (BCGT_RS18845) |
| | 10.74 | 2850691-2850756 | 1 | 2 | cell surface protein (BCGT_RS13720) |
| | 10.74 | 3183593-3183639 | 1 | 2 | NADH-quinone oxidoreductase subunit I (BCGT_RS15175) |
| | 10.04 | 2467129-2467080 | -1 | 2 | carboxylesterase LipQ (BCGT_RS11710) |
| *Mycobacterium_bovis_BCG_str_Korea_1168P* (NC_020245.2) | 17.37 | 2159066-2159116 | 1 | 1 | monooxygenase (K60_RS09975) |
| | 13.76 | 3138116-3138170 | 1 | 1 | hypothetical protein (K60_RS14900) |
| | 11.95 | 971077-971023 | -1 | 1 | hypothetical protein (K60_RS04600) |
| | 11.67 | 3396818-3396765 | -1 | 1 | hypothetical protein (K60_RS15960) |
| | 10.14 | 3426471-3426524 | 1 | 1 | AmmeMemoRadiSam system radical SAM enzyme (K60_RS16125) |
| | 15.35 | 4314752-4314824 | 1 | 2 | ESX-1 secretion-associated protein EspE (K60_RS201650) |
| | 10.74 | 3110279-3110344 | 1 | 2 | cell surface protein (K60_RS14755) |
| | 10.74 | 3442987-3443033 | 1 | 2 | NADH-quinone oxidoreductase subunit I (K60_RS16200) |
| | 10.04 | 2727512-2727463 | -1 | 2 | carboxylesterase LipQ (K60_RS12755) |
| *Mycobacterium_bovis_BCG_str_Mexico* (NC_016804.1) | 17.37 | 2168786-2168836 | 1 | 1 | monooxygenase (BCGMEX_RS10025) |
| | 13.76 | 3147833-3147887 | 1 | 1 | hypothetical protein (BCGMEX_RS14940) |
| | 11.95 | 971023-970969 | -1 | 1 | hypothetical protein (BCGMEX_RS04600) |
| | 11.67 | 3406888-3406835 | -1 | 1 | hypothetical protein (BCGMEX_RS16005) |
| | 10.14 | 3436541-3436594 | 1 | 1 | AmmeMemoRadiSam system radical SAM enzyme (BCGMEX_RS16170) |
| | 15.35 | 4288427-4288499 | 1 | 2 | ESX-1 secretion-associated protein EspE (BCGMEX_RS20025) |
| | 10.74 | 3119996-3120061 | 1 | 2 | cell surface protein (BCGMEX_RS14795) |
| | 10.74 | 3453057-3453103 | 1 | 2 | NADH-quinone oxidoreductase subunit I (BCGMEX_RS16245) |
| | 10.04 | 2737232-2737183 | -1 | 2 | carboxylesterase LipQ (BCGMEX_RS12800) |
| *Mycobacterium_bovis_BCG_str_Moreau_RDJ* (NZ_AM412059.1) | 17.37 | 2171338-2171388 | 1 | 1 | monooxygenase (BCGM_RS10045) |
| | 13.76 | 3161017-3161071 | 1 | 1 | hypothetical protein (BCGM_RS15025) |
| | 11.95 | 972380-972326 | -1 | 1 | hypothetical protein (BCGM_RS04610) |
| | 11.67 | 3418970-3418917 | -1 | 1 | hypothetical protein (BCGM_RS16085) |
| | 10.14 | 3448623-3448676 | 1 | 1 | AmmeMemoRadiSam system radical SAM enzyme (BCGM_RS16250) |

| Organism | Bit Score | ANTAR RNA hit co-ordinates | Strand | RNA Category | Annotation of the downstream gene (Locus Tag) |
|---|---|---|---|---|---|
| | 15.35 | 4279033-4279105 | 1 | 2 | ESX-1 secretion-associated protein EspE (BCGM_RS20030) |
| | 10.74 | 3133180-3133245 | 1 | 2 | cell surface protein (BCGM_RS14880) |
| | 10.74 | 3465139-3465185 | 1 | 2 | NADH-quinone oxidoreductase subunit I (BCGM_RS16325) |
| | 10.04 | 2750361-2750312 | -1 | 2 | carboxylesterase LipQ (BCGM_RS12880) |
| *Mycobacterium_bovis_BCG_str_Pasteur_1173P2* (NC_008769.1) | 17.37 | 2189421-2189471 | 1 | 1 | monooxygenase (BCG_RS10150) |
| | 13.76 | 3168468-3168522 | 1 | 1 | hypothetical protein (BCG_RS15065) |
| | 11.95 | 1000744-1000690 | -1 | 1 | hypothetical protein (BCG_RS04760) |
| | 11.67 | 3427523-3427470 | -1 | 1 | hypothetical protein (BCG_RS16125) |
| | 10.14 | 3457176-3457229 | 1 | 1 | AmmeMemoRadiSam system radical SAM enzyme (BCG_RS16290) |
| | 15.35 | 4312563-4312635 | 1 | 2 | ESX-1 secretion-associated protein EspE (BCG_RS20165) |
| | 10.74 | 3140631-3140696 | 1 | 2 | cell surface protein (BCG_RS14920) |
| | 10.74 | 3473692-3473738 | 1 | 2 | NADH-quinone oxidoreductase subunit I (BCG_RS16365) |
| | 10.04 | 2757867-2757818 | -1 | 2 | carboxylesterase LipQ (BCG_RS12930) |
| *Mycobacterium_bovis_BCG_str_Tokyo_172* (NC_012207.1) | 17.37 | 2171412-2171462 | 1 | 1 | monooxygenase (JTY_RS10050) |
| | 13.76 | 3161295-3161349 | 1 | 1 | hypothetical protein (JTY_RS15030) |
| | 11.95 | 972461-972407 | -1 | 1 | hypothetical protein (JTY_RS04620) |
| | 11.67 | 3420173-3420120 | -1 | 1 | hypothetical protein (JTY_RS16090) |
| | 10.14 | 3449826-3449879 | 1 | 1 | AmmeMemoRadiSam system radical SAM enzyme (JTY_RS16255) |
| | 15.35 | 4309752-4309824 | 1 | 2 | ESX-1 secretion-associated protein EspE (JTY_RS20165) |
| | 10.74 | 3133458-3133523 | 1 | 2 | cell surface protein (JTY_RS14885) |
| | 10.74 | 3466342-3466388 | 1 | 2 | NADH-quinone oxidoreductase subunit I (JTY_RS16330) |
| | 10.04 | 2750638-2750589 | -1 | 2 | carboxylesterase LipQ (JTY_RS12890) |
| *Mycobacterium_bovis_BCG_str_Tokyo_172_substrain_TRCS* (NZ_CP014566.1) | 17.37 | 2171410-2171460 | 1 | 1 | monooxygenase (AZH48_RS10030) |
| | 13.76 | 3161295-3161349 | 1 | 1 | hypothetical protein (JTY_RS15030) |
| | 11.95 | 972461-972407 | -1 | 1 | hypothetical protein (JTY_RS04620) |
| | 11.67 | 3420173-3420120 | -1 | 1 | hypothetical protein (JTY_RS16090) |
| | 10.14 | 3449826-3449879 | 1 | 1 | AmmeMemoRadiSam system radical SAM enzyme (JTY_RS16255) |
| | 15.35 | 4309748-4309820 | 1 | 2 | ESX-1 secretion-associated protein EspE (AZH48_RS20125) |
| | 10.74 | 3133458-3133523 | 1 | 2 | cell surface protein (JTY_RS14885) |
| | 10.74 | 3466342-3466388 | 1 | 2 | NADH-quinone oxidoreductase subunit I (JTY_RS16330) |
| | 10.04 | 2750637-2750588 | -1 | 2 | carboxylesterase LipQ (AZH48_RS12860) |
| *Mycobacterium_bovis_BCG_strain=3281* (NZ_CP008744.1) | 17.37 | 2169564-2169614 | 1 | 1 | monooxygenase (GS11_RS10040) |
| | 13.76 | 3147951-3148005 | 1 | 1 | hypothetical protein (GS11_RS14945) |

| Organism | Bit Score | ANTAR RNA hit co-ordinates | Strand | RNA Category | Annotation of the downstream gene (Locus Tag) |
|---|---|---|---|---|---|
| | 11.95 | 971089-971035 | -1 | 1 | hypothetical protein (GS11_RS04605) |
| | 11.67 | 3406829-3406776 | -1 | 1 | hypothetical protein (GS11_RS16005) |
| | 10.14 | 3436482-3436535 | 1 | 1 | AmmeMemoRadiSam system radical SAM enzyme (GS11_RS16170) |
| | 15.35 | 4334126-4334198 | 1 | 2 | ESX-1 secretion-associated protein EspE (GS11_RS20240) |
| | 10.74 | 3120114-3120179 | 1 | 2 | cell surface protein (GS11_RS14805) |
| | 10.74 | 3452998-3453044 | 1 | 2 | NADH-quinone oxidoreductase subunit I (GS11_RS16245) |
| | 10.04 | 2738001-2737952 | -1 | 2 | carboxylesterase LipQ (GS11_RS12810) |
| *Mycobacterium_bovis_BCG_strain=Russia_368* (NZ_CP009243.1) | 17.37 | 2171415-2171465 | 1 | 1 | monooxygenase (BCGR_RS10045) |
| | 13.76 | 3161300-3161354 | 1 | 1 | hypothetical protein (BCGR_RS15010) |
| | 11.95 | 972458-972404 | -1 | 1 | hypothetical protein (BCGR_RS04610) |
| | 11.67 | 3420178-3420125 | -1 | 1 | hypothetical protein (BCGR_RS16065) |
| | 10.14 | 3449831-3449884 | 1 | 1 | AmmeMemoRadiSam system radical SAM enzyme (BCGR_RS16230) |
| | 15.35 | 4308179-4308251 | 1 | 2 | ESX-1 secretion-associated protein EspE (BCGR_RS20135) |
| | 10.74 | 3133463-3133528 | 1 | 2 | cell surface protein (BCGR_RS14870) |
| | 10.74 | 3466347-3466393 | 1 | 2 | NADH-quinone oxidoreductase subunit I (BCGR_RS16305) |
| | 10.04 | 2750642-2750593 | -1 | 2 | carboxylesterase LipQ (BCGR_RS12870) |
| *Mycobacterium_bovis_strain=1595* (NZ_CP012095.1) | 17.37 | 2183860-2183910 | 1 | 1 | monooxygenase (Mb1595_RS10130) |
| | 13.76 | 3175544-3175598 | 1 | 1 | hypothetical protein (Mb1595_RS15080) |
| | 11.95 | 970804-970750 | -1 | 1 | hypothetical protein (Mb1595_RS04610) |
| | 11.67 | 3434518-3434465 | -1 | 1 | hypothetical protein (Mb1595_RS16140) |
| | 10.14 | 3464172-3464225 | 1 | 1 | AmmeMemoRadiSam system radical SAM enzyme (Mb1595_RS16305) |
| | 15.35 | 4282663-4282735 | 1 | 2 | ESX-1 secretion-associated protein EspE (Mb1595_RS20010) |
| | 10.74 | 3147760-3147825 | 1 | 2 | cell surface protein (Mb1595_RS14940) |
| | 10.74 | 3480688-3480734 | 1 | 2 | NADH-quinone oxidoreductase subunit I (Mb1595_RS16380) |
| | 10.04 | 2765960-2765911 | -1 | 2 | carboxylesterase LipQ (Mb1595_RS12965) |
| *Mycobacterium_bovis_strain=BAA-935* (NZ_CP009449.1) | 17.37 | 2147621-2147671 | 1 | 1 | monooxygenase (LH58_RS10065) |
| | 13.76 | 3123485-3123539 | 1 | 1 | hypothetical protein (LH58_RS15005) |
| | 11.95 | 964167-964113 | -1 | 1 | PE-PGRS family protein (LH58_RS04630) |
| | 11.67 | 3382156-3382103 | -1 | 1 | hypothetical protein (LH58_RS16065) |
| | 10.14 | 3411810-3411863 | 1 | 1 | AmmeMemoRadiSam system radical SAM enzyme (LH58_RS16230) |
| | 15.35 | 4298241-4298313 | 1 | 2 | ESX-1 secretion-associated protein EspE |

| Organism | Bit Score | ANTAR RNA hit co-ordinates | Strand | RNA Category | Annotation of the downstream gene (Locus Tag) |
|---|---|---|---|---|---|
| | | | | | (LH58_RS20320) |
| | 10.74 | 3095648-3095713 | 1 | 2 | cell surface protein (LH58_RS14860) |
| | 10.74 | 3428326-3428372 | 1 | 2 | NADH-quinone oxidoreductase subunit I (LH58_RS16305 |
| | 10.04 | 2715443-2715394 | -1 | 2 | carboxylesterase LipQ (LH58_RS12860) |
| *Mycobacterium_bovis_strain=BCG-1_Russia* (NZ_CP013741.1) | 17.37 | 2171711-2171761 | 1 | 1 | monooxygenase (BOVR_RS10050) |
| | 13.76 | 3161523-3161577 | 1 | 1 | hypothetical protein (BOVR_RS15025) |
| | 11.95 | 972383-972329 | -1 | 1 | hypothetical protein (BOVR_RS04610) |
| | 11.67 | 3420578-3420525 | -1 | 1 | hypothetical protein (BOVR_RS16080) |
| | 10.14 | 3450231-3450284 | 1 | 1 | AmmeMemoRadiSam system radical SAM enzyme (BOVR_RS16245) |
| | 15.35 | 4308746-4308818 | 1 | 2 | ESX-1 secretion-associated protein EspE (BOVR_RS20130) |
| | 10.74 | 3133686-3133751 | 1 | 2 | cell surface protein (BOVR_RS14880) |
| | 10.74 | 3466747-3466793 | 1 | 2 | NADH-quinone oxidoreductase subunit I (BOVR_RS16320) |
| | 10.04 | 2750937-2750888 | -1 | 2 | carboxylesterase LipQ (BOVR_RS12880) |
| *Mycobacterium_bovis_strain=SP38* (NZ_CP015773.1) | 13.76 | 2077959-2078013 | 1 | 1 | hypothetical protein (SZ58_RS09910) |
| | 13.75 | 1085811-1085858 | 1 | 1 | monooxygenase (SZ58_RS04940) |
| | 11.95 | 4221344-4221290 | -1 | 1 | PE-PGRS family protein (SZ58_RS19735) |
| | 11.67 | 2336612-2336559 | -1 | 1 | hypothetical protein (SZ58_RS10965) |
| | 10.14 | 2366266-2366319 | 1 | 1 | AmmeMemoRadiSam system radical SAM enzyme (SZ58_RS11130) |
| | 15.35 | 3181083-3181155 | 1 | 2 | ESX-1 secretion-associated protein EspE (SZ58_RS14840) |
| | 10.74 | 2050122-2050187 | 1 | 2 | cell surface protein (SZ58_RS09770) |
| | 10.74 | 2382782-2382828 | 1 | 2 | NADH-quinone oxidoreductase subunit I (SZ58_RS11205) |
| | 10.04 | 1667522-1667473 | -1 | 2 | carboxylesterase LipQ (SZ58_RS07795) |
| *Mycobacterium_canettii_CIPT_140010059* (NC_015848.1) | 13.76 | 3273444-3273498 | 1 | 1 | hypothetical protein (MCAN_RS15385) |
| | 11.67 | 3537197-3537144 | -1 | 1 | hypothetical protein (MCAN_RS16475) |
| | 10.14 | 3559166-3559219 | 1 | 1 | AmmeMemoRadiSam system radical SAM enzyme (MCAN_RS16595) |
| | 15.35 | 4410435-4410507 | 1 | 2 | ESX-1 secretion-associated protein EspE (MCAN_RS20435) |
| | 10.74 | 3245334-3245399 | 1 | 2 | cell surface protein (MCAN_RS15240) |
| | 10.74 | 3575680-3575726 | 1 | 2 | NADH-quinone oxidoreductase subunit I (MCAN_RS16670) |
| | 10.04 | 2855928-2855879 | -1 | 2 | carboxylesterase LipQ (MCAN_RS13235) |
| *Mycobacterium_chelonae_CCUG_4744* (NZ_CP007220.1) | 11.18 | 3471896-3471949 | 1 | 2 | cytochrome P450 (BB28_RS17045) |
| *Mycobacterium_fortuitum_strain=CT6* | 14.91 | 6251576-6251522 | -1 | 1 | 16S rRNA methyltransferase G (XA26_RS29760) |

| Organism | Bit Score | ANTAR RNA hit co-ordinates | Strand | RNA Category | Annotation of the downstream gene (Locus Tag) |
|---|---|---|---|---|---|
| (NZ_CP011269.1) | | | | | |
| *Mycobacterium_goodii_strain=X7B* (NZ_CP012150.1) | 16.71 | 5895932-5895882 | -1 | 1 | hypothetical protein (AFA91_RS27500) |
| *Mycobacterium_haemophilum_DSM_44634* (NZ_CP011883.2) | 11.12 | 174049-173986 | -1 | 1 | acyl-CoA dehydrogenase (B586_RS00815) |
| | 10.14 | 1112421-1112495 | 1 | 1 | hypothetical protein (B586_RS05315) |
| *Mycobacterium_immunogenum_strain=CCUG_47286* (NZ_CP011530.1) | 12.01 | 5249432-5249478 | 1 | 1 | GntR family transcriptional regulator (ABG82_RS25935) |
| | 11.54 | 14770-14818 | 1 | 1 | hypothetical protein (ABG82_RS00080) |
| | 14.84 | 4673950-4674040 | 1 | 2 | amino acid permease (ABG82_RS23295) |
| | 10.75 | 3685270-3685323 | 1 | 2 | cytochrome P450 (ABG82_RS18165) |
| *Mycobacterium_indicus_pranii_MTCC_9506* (NC_018612.1) | 11.46 | 2593675-2593620 | -1 | 1 | DNA-directed RNA polymerase sigma-70 factor (MIP_RS11885) |
| *Mycobacterium_leprae_Br4923* (NC_011896.1) | 12.71 | 567138-567198 | 1 | 1 | hypothetical protein (MLBR_RS02290) |
| | 12.29 | 3088433-3088494 | 1 | 1 | hypothetical protein (MLBR_RS13170) |
| | 15.97 | 465129-465198 | 1 | 2 | bifunctional ADP-dependent (S)-NAD(P)H-hydrate dehydratase/NAD(P)H-hydrate epimerase (MLBR_RS01830) |
| *Mycobacterium_liflandii_128FXT* (NC_020133.1) | 10.88 | 3552989-3553040 | 1 | 1 | hypothetical protein (MULP_RS15355) |
| | 11.11 | 3257155-3257207 | 1 | 2 | long-chain-fatty-acid--CoA ligase (MULP_RS14220) |
| *Mycobacterium_marinum_E11* (NZ_HG917972.2) | 10.88 | 2705668-2705617 | -1 | 1 | hypothetical protein (MMARE11_RS11245) |
| | 11.11 | 2438081-2438029 | -1 | 2 | long-chain-fatty-acid--CoA ligase (MMARE11_RS10150) |
| *Mycobacterium_marinum_M* (NC_010612.1) | 10.88 | 2842763-2842712 | -1 | 1 | hypothetical protein (MMAR_RS11735) |
| | 11.11 | 2553321-2553269 | -1 | 2 | long-chain-fatty-acid--CoA ligase (MMAR_RS10535) |
| *Mycobacterium_neoaurum_VKM_Ac-1815D* (NC_023036.2) | 14.17 | 4181172-4181233 | 1 | 1 | hypothetical protein (D174_RS19485) |
| | 10.32 | 1555181-1555233 | 1 | 2 | hypothetical protein (D174_RS07385) |
| *Mycobacterium_rhodesiae_NBB3* (NC_016604.1) | 15.67 | 761121-761068 | -1 | 1 | DNA-directed RNA polymerase subunit beta (MYCRHN_RS03650) |
| | 15 | 3909900-3909952 | 1 | 1 | 2-hydroxy-6-ketonona-2,4-dienedioic acid hydrolase (MYCRHN_RS19080) |
| | 11.01 | 5093059-5093112 | 1 | 1 | hypothetical protein (MYCRHN_RS24420) |
| *Mycobacterium_tuberculosis_49-02* (NZ_HG813240.1) | 17.37 | 2173566-2173616 | 1 | 1 | monooxygenase (MT49_RS10070) |
| | 13.76 | 3202634-3202688 | 1 | 1 | hypothetical protein (MT49_RS15270) |
| | 11.95 | 968728-968674 | -1 | 1 | hypothetical protein (MT49_RS04585) |
| | 11.67 | 3462244-3462191 | -1 | 1 | hypothetical protein (MT49_RS16330) |
| | 10.14 | 3495085-3495138 | 1 | 1 | AmmeMemoRadiSam system radical SAM enzyme (MT49_RS16530) |
| | 15.35 | 4340917-4340989 | 1 | 2 | ESX-1 secretion-associated protein EspE (MT49_RS20365) |
| | 10.74 | 3174691-3174756 | 1 | 2 | cell surface protein (MT49_RS15130) |
| | 10.74 | 3511601-3511647 | 1 | 2 | NADH-quinone oxidoreductase subunit I (MT49_RS16605) |

| Organism | Bit Score | ANTAR RNA hit co-ordinates | Strand | RNA Category | Annotation of the downstream gene (Locus Tag) |
|---|---|---|---|---|---|
| | 10.04 | 2789495-2789446 | -1 | 2 | carboxylesterase (MT49_RS13115) |
| *Mycobacterium_tuberculosis_7199-99* (NC_020089.1) | 17.37 | 2191699-2191749 | 1 | 1 | monooxygenase (MT7199_RS10100) |
| | 13.76 | 3211910-3211964 | 1 | 1 | hypothetical protein (MT7199_RS15215) |
| | 11.95 | 973598-973544 | -1 | 1 | hypothetical protein (MT7199_RS04590) |
| | 11.67 | 3468909-3468856 | -1 | 1 | hypothetical protein (MT7199_RS16270) |
| | 10.14 | 3501049-3501102 | 1 | 1 | AmmeMemoRadiSam system radical SAM enzyme (MT7199_RS16465) |
| | 15.35 | 4349871-4349943 | 1 | 2 | ESX-1 secretion-associated protein EspE (MT7199_RS20340) |
| | 10.74 | 3184160-3184225 | 1 | 2 | cell surface protein (MT7199_RS15075) |
| | 10.74 | 3517565-3517611 | 1 | 2 | NADH-quinone oxidoreductase subunit I (MT7199_RS16540) |
| | 10.04 | 2794672-2794623 | -1 | 2 | carboxylesterase LipQ (MT7199_RS13040) |
| *Mycobacterium_tuberculosis_BT1* (NZ_CP002883.1) | 17.37 | 2173137-2173187 | 1 | 1 | monooxygenase (HKBT1_RS10085) |
| | 13.76 | 3196721-3196775 | 1 | 1 | hypothetical protein (HKBT1_RS15245) |
| | 11.95 | 969042-968988 | -1 | 1 | hypothetical protein (HKBT1_RS04585) |
| | 11.67 | 3456830-3456777 | -1 | 1 | hypothetical protein (HKBT1_RS16310) |
| | 10.14 | 3489671-3489724 | 1 | 1 | AmmeMemoRadiSam system radical SAM enzyme (HKBT1_RS16510) |
| | 15.35 | 4328038-4328110 | 1 | 2 | ESX-1 secretion-associated protein EspE (HKBT1_RS20335) |
| | 10.74 | 3168827-3168892 | 1 | 2 | cell surface protein (HKBT1_RS15105) |
| | 10.74 | 3506187-3506233 | 1 | 2 | NADH-quinone oxidoreductase subunit I (HKBT1_RS16585) |
| | 10.04 | 2783665-2783616 | -1 | 2 | carboxylesterase (HKBT1_RS13090) |
| *Mycobacterium_tuberculosis_BT2* (NZ_CP002882.1) | 17.37 | 2166350-2166400 | 1 | 1 | monooxygenase (HKBT2_RS10060) |
| | 13.76 | 3190985-3191039 | 1 | 1 | hypothetical protein (HKBT2_RS15230) |
| | 11.95 | 968987-968933 | -1 | 1 | hypothetical protein (HKBT2_RS04585) |
| | 11.67 | 3451102-3451049 | -1 | 1 | hypothetical protein (HKBT2_RS16300) |
| | 10.14 | 3483943-3483996 | 1 | 1 | AmmeMemoRadiSam system radical SAM enzyme (HKBT2_RS16500) |
| | 15.35 | 4330472-4330544 | 1 | 2 | ESX-1 secretion-associated protein EspE (HKBT2_RS20350) |
| | 10.74 | 3163042-3163107 | 1 | 2 | cell surface protein (HKBT2_RS15090) |
| | 10.74 | 3500459-3500505 | 1 | 2 | NADH-quinone oxidoreductase subunit I (HKBT2_RS16575) |
| | 10.04 | 2776270-2776221 | -1 | 2 | carboxylesterase (HKBT2_RS13070) |
| *Mycobacterium_tuberculosis_CCDC5079* (NC_021251.1) | 15.31 | 2174974-2175038 | 1 | 1 | monooxygenase (CFBS_RS10115) |
| | 13.76 | 3203172-3203226 | 1 | 1 | hypothetical protein (CFBS_RS15325) |
| | 11.95 | 968849-968795 | -1 | 1 | hypothetical protein (CFBS_RS04600) |

| Organism | Bit Score | ANTAR RNA hit co-ordinates | Strand | RNA Category | Annotation of the downstream gene (Locus Tag) |
|---|---|---|---|---|---|
| | 11.67 | 3463287-3463234 | -1 | 1 | hypothetical protein (CFBS_RS16400) |
| | 10.14 | 3496128-3496181 | 1 | 1 | AmmeMemoRadiSam system radical SAM enzyme (CFBS_RS16600) |
| | 15.35 | 4342863-4342935 | 1 | 2 | ESX-1 secretion-associated protein EspE (CFBS_RS20430) |
| | 10.74 | 3175229-3175294 | 1 | 2 | cell surface protein (CFBS_RS15180) |
| | 10.74 | 3512644-3512690 | 1 | 2 | NADH-quinone oxidoreductase subunit I (CFBS_RS16675) |
| | 10.04 | 2788597-2788548 | -1 | 2 | carboxylesterase (CFBS_RS13145) |
| *Mycobacterium_tuberculosis_CCDC5180_1* (NC_017522.1) | 17.37 | 2168763-2168813 | 1 | 1 | monooxygenase (CCDC5180_RS10115) |
| | 13.76 | 3195483-3195537 | 1 | 1 | hypothetical protein (CCDC5180_RS15305) |
| | 11.95 | 966435-966381 | -1 | 1 | PE-PGRS family protein (CCDC5180_RS04600) |
| | 11.67 | 3456873-3456820 | -1 | 1 | hypothetical protein (CCDC5180_RS16385) |
| | 10.14 | 3489714-3489767 | 1 | 1 | AmmeMemoRadiSam system radical SAM enzyme (CCDC5180_RS16585) |
| | 15.35 | 4334516-4334588 | 1 | 2 | ESX-1 secretion-associated protein EspE (CCDC5180_RS20410) |
| | 10.74 | 3167540-3167605 | 1 | 2 | cell surface protein (CCDC5180_RS15165) |
| | 10.74 | 3506230-3506276 | 1 | 2 | NADH-quinone oxidoreductase subunit I (CCDC5180_RS16660) |
| | 10.04 | 2782266-2782217 | -1 | 2 | carboxylesterase (CCDC5180_RS13135) |
| *Mycobacterium_tuberculosis_CCDC5180_2* (NZ_CP002885.1) | 17.37 | 2173333-2173383 | 1 | 1 | monooxygenase (CFBR_RS10085) |
| | 13.76 | 3200058-3200112 | 1 | 1 | hypothetical protein (CFBR_RS15260) |
| | 11.95 | 968803-968749 | -1 | 1 | hypothetical protein (CFBR_RS04585) |
| | 11.67 | 3462061-3462008 | -1 | 1 | hypothetical protein (CFBR_RS16355) |
| | 10.14 | 3494902-3494955 | 1 | 1 | AmmeMemoRadiSam system radical SAM enzyme (CFBR_RS16555) |
| | 15.35 | 4342884-4342956 | 1 | 2 | ESX-1 secretion-associated protein EspE (CFBR_RS20395) |
| | 10.74 | 3172115-3172180 | 1 | 2 | cell surface protein (CFBR_RS15120) |
| | 10.74 | 3511418-3511464 | 1 | 2 | NADH-quinone oxidoreductase subunit I (CFBR_RS16630) |
| | 10.04 | 2786843-2786794 | -1 | 2 | carboxylesterase (CFBR_RS13105) |
| *Mycobacterium_tuberculosis_CDC1551* (NC_002755.2) | 15.31 | 2184611-2184675 | 1 | 1 | monooxygenase (MT_RS10100) |
| | 13.76 | 3208690-3208744 | 1 | 1 | hypothetical protein (MT_RS15230) |
| | 11.95 | 970297-970243 | -1 | 1 | hypothetical protein (MT_RS04590) |
| | 11.6 | 3468832-3468780 | -1 | 1 | hypothetical protein (MT_RS16300) |
| | 10.14 | 3499614-3499667 | 1 | 1 | AmmeMemoRadiSam system radical SAM enzyme (MT_RS16480) |
| | 15.35 | 4332485-4332557 | 1 | 2 | ESX-1 secretion-associated protein EspE (MT_RS20285) |
| | 10.74 | 3180854-3180919 | 1 | 2 | cell surface protein (MT_RS15090) |

| Organism | Bit Score | ANTAR RNA hit co-ordinates | Strand | RNA Category | Annotation of the downstream gene (Locus Tag) |
|---|---|---|---|---|---|
| | 10.74 | 3516130-3516176 | 1 | 2 | NADH-quinone oxidoreductase subunit I (MT_RS16555) |
| | 10.04 | 2789546-2789497 | -1 | 2 | carboxylesterase LipQ (MT_RS13045) |
| *Mycobacterium_tuberculosis_CTRI-2* (NC_017524.1) | 17.37 | 2180824-2180874 | 1 | 1 | monooxygenase (MTCTRI2_RS10125) |
| | 13.76 | 3210750-3210804 | 1 | 1 | hypothetical protein (MTCTRI2_RS15310) |
| | 11.95 | 970700-970646 | -1 | 1 | hypothetical protein (MTCTRI2_RS04615) |
| | 11.67 | 3468822-3468769 | -1 | 1 | hypothetical protein (MTCTRI2_RS16380) |
| | 10.14 | 3501273-3501326 | 1 | 1 | AmmeMemoRadiSam system radical SAM enzyme (MTCTRI2_RS16580) |
| | 15.35 | 4327151-4327223 | 1 | 2 | ESX-1 secretion-associated protein EspE (MTCTRI2_RS20340) |
| | 10.74 | 3182967-3183032 | 1 | 2 | cell surface protein (MTCTRI2_RS15170) |
| | 10.74 | 3517789-3517835 | 1 | 2 | NADH-quinone oxidoreductase subunit I (MTCTRI2_RS16655) |
| | 10.04 | 2789010-2788961 | -1 | 2 | carboxylesterase LipQ (MTCTRI2_RS13105) |
| *Mycobacterium_tuberculosis_EAI5* (NC_021740.1) | 17.37 | 2181844-2181894 | 1 | 1 | monooxygenase (M943_RS10115) |
| | 13.76 | 3200965-3201019 | 1 | 1 | hypothetical protein (M943_RS15230) |
| | 11.95 | 968935-968881 | -1 | 1 | hypothetical protein (M943_RS04590) |
| | 11.67 | 3459464-3459411 | -1 | 1 | hypothetical protein (M943_RS16295) |
| | 10.14 | 3490553-3490606 | 1 | 1 | AmmeMemoRadiSam system radical SAM enzyme (M943_RS16480) |
| | 15.35 | 4319795-4319867 | 1 | 2 | ESX-1 secretion-associated protein EspE (M943_RS20215) |
| | 10.74 | 3173129-3173194 | 1 | 2 | cell surface protein (M943_RS15085) |
| | 10.74 | 3507069-3507115 | 1 | 2 | NADH-quinone oxidoreductase subunit I (M943_RS16555) |
| | 10.04 | 2781850-2781801 | -1 | 2 | carboxylesterase LipQ (M943_RS13040) |
| *Mycobacterium_tuberculosis_EAI5NITR206* (NC_021194.1) | 17.37 | 2181480-2181530 | 1 | 1 | monooxygenase (J114_RS10135) |
| | 13.76 | 3200395-3200449 | 1 | 1 | hypothetical protein (J114_RS15275) |
| | 11.95 | 968752-968698 | -1 | 1 | hypothetical protein (J114_RS04595) |
| | 11.67 | 3458814-3458761 | -1 | 1 | hypothetical protein (J114_RS16335) |
| | 10.14 | 3489881-3489934 | 1 | 1 | AmmeMemoRadiSam system radical SAM enzyme (J114_RS16520) |
| | 15.35 | 4318936-4319008 | 1 | 2 | ESX-1 secretion-associated protein EspE (J114_RS20245) |
| | 10.74 | 3172580-3172645 | 1 | 2 | cell surface protein (J114_RS15125) |
| | 10.74 | 3506397-3506443 | 1 | 2 | NADH-quinone oxidoreductase subunit I (J114_RS16595) |
| | 10.04 | 2781362-2781313 | -1 | 2 | carboxylesterase LipQ (J114_RS13075) |
| *Mycobacterium_tuberculosis_F11* (NC_009565.1) | 17.37 | 2195516-2195566 | 1 | 1 | monooxygenase (TBFG_RS10180) |
| | 13.76 | 3226065-3226119 | 1 | 1 | hypothetical protein (TBFG_RS15375) |
| | 11.95 | 974108-974054 | -1 | 1 | hypothetical protein (TBFG_RS04605) |
| | 11.67 | 3484801-3484748 | -1 | 1 | hypothetical protein (TBFG_RS16430) |

| Organism | Bit Score | ANTAR RNA hit co-ordinates | Strand | RNA Category | Annotation of the downstream gene (Locus Tag) |
|---|---|---|---|---|---|
| | 10.14 | 3517252-3517305 | 1 | 1 | AmmeMemoRadiSam system radical SAM enzyme (TBFG_RS16630) |
| | 15.35 | 4353162-4353234 | 1 | 2 | ESX-1 secretion-associated protein EspE (TBFG_RS20420) |
| | 10.74 | 3198229-3198294 | 1 | 2 | cell surface protein (TBFG_RS15235) |
| | 10.74 | 3533768-3533814 | 1 | 2 | NADH-quinone oxidoreductase subunit I (TBFG_RS16705) |
| | 10.04 | 2807374-2807325 | -1 | 2 | carboxylesterase (TBFG_RS13185) |
| *Mycobacterium_tuberculosis_H37Ra* (NC_009525.1) | 17.37 | 2197160-2197210 | 1 | 1 | monooxygenase (MRA_RS10230) |
| | 13.76 | 3226528-3226582 | 1 | 1 | hypothetical protein (MRA_RS15435) |
| | 11.95 | 971590-971536 | -1 | 1 | hypothetical protein (MRA_RS04620) |
| | 11.67 | 3485193-3485140 | -1 | 1 | hypothetical protein (MRA_RS16500) |
| | 10.14 | 3516285-3516338 | 1 | 1 | AmmeMemoRadiSam system radical SAM enzyme (MRA_RS16685) |
| | 15.35 | 4348608-4348680 | 1 | 2 | ESX-1 secretion-associated protein EspE (MRA_RS20490) |
| | 10.74 | 3198691-3198756 | 1 | 2 | cell surface protein (MRA_RS15290) |
| | 10.74 | 3532801-3532847 | 1 | 2 | NADH-quinone oxidoreductase subunit I (MRA_RS16760) |
| | 10.04 | 2806026-2805977 | -1 | 2 | carboxylesterase LipQ (MRA_RS13235) |
| *Mycobacterium_tuberculosis_H37Rv_1* (NC_000962.3) | 17.37 | 2187261-2187311 | 1 | 1 | monooxygenase (Rv1936) |
| | 13.76 | 3214536-3214590 | 1 | 1 | lipoprotein LppW (Rv2905) |
| | 11.95 | 970284-970230 | -1 | 1 | PE-PGRS family protein PE_PGRS15 (Rv0872c) |
| | 11.67 | 3473037-3472984 | -1 | 1 | transmembrane protein (Rv3104c) |
| | 10.14 | 3504129-3504182 | 1 | 1 | pyruvate formate lyase activating protein PflA (Rv3138) |
| | 15.35 | 4340162-4340234 | 1 | 2 | ESX-1 secretion-associated protein EspE (Rv3864) |
| | 10.74 | 3186699-3186764 | 1 | 2 | major secreted immunogenic protein Mpt70 (Rv2875) |
| | 10.74 | 3520645-3520691 | 1 | 2 | NADH-quinone oxidoreductase subunit I (Rv3153) |
| | 10.04 | 2794058-2794009 | -1 | 2 | carboxylesterase LipQ (Rv2485c) |
| *Mycobacterium_tuberculosis_H37Rv_2* (NC_018143.2) | 17.37 | 2187268-2187318 | 1 | 1 | monooxygenase (RVBD_RS10160) |
| | 13.76 | 3214546-3214600 | 1 | 1 | hypothetical protein (RVBD_RS15335) |
| | 11.95 | 970287-970233 | -1 | 1 | hypothetical protein (RVBD_RS04610) |
| | 11.67 | 3473034-3472981 | -1 | 1 | hypothetical protein (RVBD_RS16395) |
| | 10.14 | 3504126-3504179 | 1 | 1 | AmmeMemoRadiSam system radical SAM enzyme (RVBD_RS16580) |
| | 15.35 | 4340340-4340412 | 1 | 2 | ESX-1 secretion-associated protein EspE (RVBD_RS20380) |
| | 10.74 | 3186709-3186774 | 1 | 2 | cell surface protein (RVBD_RS15190) |
| | 10.74 | 3520642-3520688 | 1 | 2 | NADH-quinone oxidoreductase subunit I (RVBD_RS16655) |
| | 10.04 | 2794068-2794019 | -1 | 2 | carboxylesterase LipQ (RVBD_RS13135) |
| *Mycobacterium_tuberculosis_H37Rv_3* | 17.37 | 2182565-2182615 | 1 | 1 | monooxygenase (LH57_RS10210) |

| Organism | Bit Score | ANTAR RNA hit co-ordinates | Strand | RNA Category | Annotation of the downstream gene (Locus Tag) |
|---|---|---|---|---|---|
| (NZ_CP009480.1) | 13.76 | 3207732-3207786 | 1 | 1 | hypothetical protein (LH57_RS15425) |
| | 11.95 | 967338-967284 | -1 | 1 | hypothetical protein (LH57_RS04615) |
| | 11.67 | 3466182-3466129 | -1 | 1 | hypothetical protein (LH57_RS16485) |
| | 10.14 | 3497274-3497327 | 1 | 1 | AmmeMemoRadiSam system radical SAM enzyme (LH57_RS16670) |
| | 15.35 | 4332571-4332643 | 1 | 2 | ESX-1 secretion-associated protein EspE (LH57_RS20485) |
| | 10.74 | 3179895-3179960 | 1 | 2 | cell surface protein (LH57_RS15280) |
| | 10.74 | 3513790-3513836 | 1 | 2 | NADH-quinone oxidoreductase subunit I (LH57_RS16745) |
| | 10.04 | 2788486-2788437 | -1 | 2 | carboxylesterase LipQ (LH57_RS13205) |
| *Mycobacterium_tuberculosis_H37RvSiena* (NZ_CP007027.1) | 17.37 | 2186649-2186699 | 1 | 1 | monooxygenase (Y980_RS10165) |
| | 13.76 | 3213924-3213978 | 1 | 1 | hypothetical protein (Y980_RS15345) |
| | 11.95 | 970287-970233 | -1 | 1 | hypothetical protein (RVBD_RS04610) |
| | 11.67 | 3472419-3472366 | -1 | 1 | hypothetical protein (Y980_RS16405) |
| | 10.14 | 3503511-3503564 | 1 | 1 | AmmeMemoRadiSam system radical SAM enzyme (Y980_RS16590) |
| | 15.35 | 4339542-4339614 | 1 | 2 | ESX-1 secretion-associated protein EspE (Y980_RS20390) |
| | 10.74 | 3186087-3186152 | 1 | 2 | cell surface protein (Y980_RS15200) |
| | 10.74 | 3520027-3520073 | 1 | 2 | NADH-quinone oxidoreductase subunit I (Y980_RS16665) |
| | 10.04 | 2793446-2793397 | -1 | 2 | carboxylesterase LipQ (Y980_RS13145) |
| *Mycobacterium_tuberculosis_HKBS1* (NZ_CP002871.1) | 17.37 | 2172407-2172457 | 1 | 1 | monooxygenase (HKBS1_RS10075) |
| | 13.76 | 3199246-3199300 | 1 | 1 | hypothetical protein (HKBS1_RS15250) |
| | 11.95 | 968808-968754 | -1 | 1 | hypothetical protein (HKBS1_RS04585) |
| | 11.67 | 3461076-3461023 | -1 | 1 | hypothetical protein (HKBS1_RS16335) |
| | 10.14 | 3493917-3493970 | 1 | 1 | AmmeMemoRadiSam system radical SAM enzyme (HKBS1_RS16535) |
| | 15.35 | 4336467-4336539 | 1 | 2 | ESX-1 secretion-associated protein EspE (HKBS1_RS20365) |
| | 10.74 | 3171303-3171368 | 1 | 2 | cell surface protein (HKBS1_RS15110) |
| | 10.74 | 3510433-3510479 | 1 | 2 | NADH-quinone oxidoreductase subunit I (HKBS1_RS16610) |
| | 10.04 | 2786032-2785983 | -1 | 2 | carboxylesterase (HKBS1_RS13095) |
| *Mycobacterium_tuberculosis_K* (NZ_CP007803.1) | 15.31 | 2159824-2159888 | 1 | 1 | monooxygenase (MTBK_RS09995) |
| | 13.76 | 3189638-3189692 | 1 | 1 | hypothetical protein (MTBK_RS15185) |
| | 11.95 | 963585-963531 | -1 | 1 | hypothetical protein (MTBK_RS04560) |
| | 11.67 | 3449942-3449889 | -1 | 1 | hypothetical protein (MTBK_RS16250) |
| | 10.14 | 3482372-3482425 | 1 | 1 | AmmeMemoRadiSam system radical SAM enzyme (MTBK_RS16435) |
| | 15.35 | 4314056-4314128 | 1 | 2 | ESX-1 secretion-associated protein EspE |

| Organism | Bit Score | ANTAR RNA hit co-ordinates | Strand | RNA Category | Annotation of the downstream gene (Locus Tag) |
|---|---|---|---|---|---|
| | | | | | (MTBK_RS20230) |
| | 10.74 | 3161696-3161761 | 1 | 2 | cell surface protein (MTBK_RS15045) |
| | 10.74 | 3498888-3498934 | 1 | 2 | NADH-quinone oxidoreductase subunit I (MTBK_RS16510) |
| | 10.04 | 2777386-2777337 | -1 | 2 | carboxylesterase (MTBK_RS13030) |
| *Mycobacterium_tuberculosis_KZN_1435_MDR* (NC_012943.1) | 17.37 | 2235812-2235762 | -1 | 1 | monooxygenase (TBMG_RS10750) |
| | 13.76 | 1199292-1199238 | -1 | 1 | hypothetical protein (TBMG_RS05535) |
| | 11.95 | 3441695-3441749 | 1 | 1 | hypothetical protein (TBMG_RS16250) |
| | 11.67 | 940739-940792 | 1 | 1 | hypothetical protein (TBMG_RS04470) |
| | 10.14 | 3504952-3505005 | 1 | 1 | AmmeMemoRadiSam system radical SAM enzyme (TBMG_RS16580) |
| | 15.35 | 4327108-4327180 | 1 | 2 | ESX-1 secretion-associated protein EspE (TBMG_RS20315) |
| | 10.74 | 1227128-1227063 | -1 | 2 | cell surface protein (TBMG_RS05675) |
| | 10.74 | 3521468-3521514 | 1 | 2 | NADH-quinone oxidoreductase subunit I (TBMG_RS16655) |
| | 10.04 | 1620982-1621031 | 1 | 2 | carboxylesterase LipQ (TBMG_RS07750) |
| *Mycobacterium_tuberculosis_KZN_4207* (NC_016768.1) | 17.37 | 2232543-2232493 | -1 | 1 | monooxygenase (TBSG_RS10740) |
| | 13.76 | 1199290-1199236 | -1 | 1 | hypothetical protein (TBSG_RS05545) |
| | 11.95 | 3438428-3438482 | 1 | 1 | hypothetical protein (TBSG_RS16240) |
| | 11.67 | 940742-940795 | 1 | 1 | hypothetical protein (TBSG_RS04485) |
| | 10.14 | 3501686-3501739 | 1 | 1 | AmmeMemoRadiSam system radical SAM enzyme (TBSG_RS16570) |
| | 15.35 | 4323712-4323784 | 1 | 2 | ESX-1 secretion-associated protein EspE (TBSG_RS20305) |
| | 10.74 | 1227126-1227061 | -1 | 2 | cell surface protein (TBSG_RS05685) |
| | 10.74 | 3518202-3518248 | 1 | 2 | NADH-quinone oxidoreductase subunit I (TBSG_RS16645) |
| | 10.04 | 1620926-1620975 | 1 | 2 | carboxylesterase LipQ (TBSG_RS07760) |
| *Mycobacterium_tuberculosis_KZN_605_XDR* (NC_018078.1) | 17.37 | 2235693-2235643 | -1 | 1 | monooxygenase (TBXG_RS10755) |
| | 13.76 | 1199404-1199350 | -1 | 1 | hypothetical protein (TBXG_RS05540) |
| | 11.95 | 3441449-3441503 | 1 | 1 | hypothetical protein (TBXG_RS16255) |
| | 11.67 | 940744-940797 | 1 | 1 | hypothetical protein (TBXG_RS04475) |
| | 10.14 | 3506065-3506118 | 1 | 1 | AmmeMemoRadiSam system radical SAM enzyme (TBXG_RS16595) |
| | 15.35 | 4327977-4328049 | 1 | 2 | ESX-1 secretion-associated protein EspE (TBXG_RS20330) |
| | 10.74 | 1227240-1227175 | -1 | 2 | cell surface protein (TBXG_RS05680) |
| | 10.74 | 3522581-3522627 | 1 | 2 | NADH-quinone oxidoreductase subunit I (TBXG_RS16670) |
| | 10.04 | 1621097-1621146 | 1 | 2 | carboxylesterase LipQ (TBXG_RS07755) |

| Organism | Bit Score | ANTAR RNA hit co-ordinates | Strand | RNA Category | Annotation of the downstream gene (Locus Tag) |
|---|---|---|---|---|---|
| *Mycobacterium_tuberculosis_str_BeijingNITR203* (NC_021054.1) | 19.26 | 2187067-2187117 | 1 | 1 | monooxygenase (J112_RS10210) |
| | 13.76 | 3214228-3214282 | 1 | 1 | hypothetical protein (J112_RS15385) |
| | 11.95 | 970192-970138 | -1 | 1 | PE-PGRS family protein (J112_RS04620) |
| | 11.67 | 3472698-3472645 | -1 | 1 | hypothetical protein (J112_RS16440) |
| | 10.14 | 3503789-3503842 | 1 | 1 | AmmeMemoRadiSam system radical SAM enzyme (J112_RS16625) |
| | 15.35 | 4339761-4339833 | 1 | 2 | ESX-1 secretion-associated protein EspE (J112_RS20450) |
| | 10.74 | 3186391-3186456 | 1 | 2 | cell surface protein (J112_RS15245) |
| | 10.74 | 3520296-3520342 | 1 | 2 | NADH-quinone oxidoreductase subunit I (J112_RS16700) |
| | 10.04 | 2793805-2793756 | -1 | 2 | carboxylesterase (J112_RS13200) |
| *Mycobacterium_tuberculosis_str_Erdman_=_ATCC _35801* (NC_020559.1) | 17.37 | 2178334-2178384 | 1 | 1 | monooxygenase (ERDMAN_RS10140) |
| | 13.76 | 3203020-3203074 | 1 | 1 | hypothetical protein (ERDMAN_RS15290) |
| | 11.95 | 967780-967726 | -1 | 1 | hypothetical protein (ERDMAN_RS04620) |
| | 11.67 | 3459301-3459248 | -1 | 1 | hypothetical protein (ERDMAN_RS16340) |
| | 10.14 | 3490082-3490135 | 1 | 1 | AmmeMemoRadiSam system radical SAM enzyme (ERDMAN_RS16520) |
| | 15.35 | 4321068-4321140 | 1 | 2 | ESX-1 secretion-associated protein EspE (ERDMAN_RS20375) |
| | 10.74 | 3175270-3175335 | 1 | 2 | cell surface protein (ERDMAN_RS15150) |
| | 10.74 | 3506597-3506643 | 1 | 2 | NADH-quinone oxidoreductase subunit I (ERDMAN_RS16595) |
| | 10.04 | 2781645-2781596 | -1 | 2 | carboxylesterase LipQ (ERDMAN_RS13075) |
| *Mycobacterium_tuberculosis_str_Haarlem* (NC_022350.1) | 17.37 | 2186892-2186942 | 1 | 1 | monooxygenase (TBHG_RS10090) |
| | 13.76 | 3209954-3210008 | 1 | 1 | hypothetical protein (TBHG_RS15220) |
| | 11.95 | 973863-973809 | -1 | 1 | hypothetical protein (TBHG_RS04590) |
| | 11.67 | 3466754-3466701 | -1 | 1 | mechanosensitive ion channel protein MscS (TBHG_RS16270) |
| | 10.14 | 3497535-3497588 | 1 | 1 | AmmeMemoRadiSam system radical SAM enzyme (TBHG_RS16450) |
| | 15.35 | 4337000-4337072 | 1 | 2 | ESX-1 secretion-associated protein EspE (TBHG_RS20290) |
| | 10.74 | 3182204-3182269 | 1 | 2 | cell surface protein (TBHG_RS15080) |
| | 10.74 | 3514051-3514097 | 1 | 2 | NADH-quinone oxidoreductase subunit I (TBHG_RS16525) |
| | 10.04 | 2788664-2788615 | -1 | 2 | carboxylesterase LipQ (TBHG_RS13020) |
| *Mycobacterium_tuberculosis_str_Kurono* (NZ_AP014573.1) | 17.37 | 2192698-2192748 | 1 | 1 | monooxygenase (KURONO_RS10185) |
| | 13.76 | 3220427-3220481 | 1 | 1 | hypothetical protein (KURONO_RS15365) |
| | 11.95 | 974384-974330 | -1 | 1 | PE family protein (KURONO_RS04615) |
| | 11.67 | 3479384-3479331 | -1 | 1 | hypothetical protein (KURONO_RS16425) |
| | 10.14 | 3511835-3511888 | 1 | 1 | AmmeMemoRadiSam system radical SAM enzyme |

| Organism | Bit Score | ANTAR RNA hit co-ordinates | Strand | RNA Category | Annotation of the downstream gene (Locus Tag) |
|---|---|---|---|---|---|
| | | | | | (KURONO_RS16620) |
| | 15.35 | 4343757-4343829 | 1 | 2 | ESX-1 secretion-associated protein EspE (KURONO_RS20405) |
| | 10.74 | 3192537-3192602 | 1 | 2 | cell surface protein (KURONO_RS15220) |
| | 10.74 | 3528351-3528397 | 1 | 2 | NADH-quinone oxidoreductase subunit I (KURONO_RS16695) |
| | 10.04 | 2798460-2798411 | -1 | 2 | carboxylesterase LipQ (KURONO_RS13160) |
| *Mycobacterium_tuberculosis_strain=22103* (NZ_CP010339.1) | 17.37 | 2185332-2185382 | 1 | 1 | monooxygenase (RN14_RS10075) |
| | 13.76 | 3212420-3212474 | 1 | 1 | hypothetical protein (RN14_RS15220) |
| | 11.95 | 972496-972442 | -1 | 1 | PE-PGRS family protein (RN14_RS04595) |
| | 11.67 | 3471484-3471431 | -1 | 1 | hypothetical protein (RN14_RS16300) |
| | 10.14 | 3503913-3503966 | 1 | 1 | AmmeMemoRadiSam system radical SAM enzyme (RN14_RS16485) |
| | 15.35 | 4328067-4328139 | 1 | 2 | ESX-1 secretion-associated protein EspE (RN14_RS20275) |
| | 10.74 | 3184584-3184649 | 1 | 2 | cell surface protein (RN14_RS15080) |
| | 10.74 | 3520429-3520475 | 1 | 2 | NADH-quinone oxidoreductase subunit I (RN14_RS16560) |
| | 10.04 | 2793261-2793212 | -1 | 2 | carboxylesterase LipQ (RN14_RS13030) |
| *Mycobacterium_tuberculosis_strain=22115* (NZ_CP010337.1) | 17.37 | 2190192-2190242 | 1 | 1 | monooxygenase (RN12_RS10150) |
| | 13.76 | 3221258-3221312 | 1 | 1 | hypothetical protein (RN12_RS15320) |
| | 11.95 | 972797-972743 | -1 | 1 | PE-PGRS family protein (RN12_RS04585) |
| | 11.67 | 3482005-3481952 | -1 | 1 | hypothetical protein (RN12_RS16400) |
| | 10.14 | 3513096-3513149 | 1 | 1 | AmmeMemoRadiSam system radical SAM enzyme (RN12_RS16585) |
| | 15.35 | 4330406-4330478 | 1 | 2 | ESX-1 secretion-associated protein EspE (RN12_RS20370) |
| | 10.74 | 3193423-3193488 | 1 | 2 | cell surface protein (RN12_RS15180) |
| | 10.74 | 3529612-3529658 | 1 | 2 | NADH-quinone oxidoreductase subunit I (RN12_RS16660) |
| | 10.04 | 2801925-2801876 | -1 | 2 | carboxylesterase LipQ (RN12_RS13125) |
| *Mycobacterium_tuberculosis_strain=26105* (NZ_CP010340.1) | 17.37 | 2199709-2199759 | 1 | 1 | monooxygenase (RN15_RS10215) |
| | 13.76 | 3233211-3233265 | 1 | 1 | hypothetical protein (RN15_RS15415) |
| | 11.95 | 977074-977020 | -1 | 1 | PE-PGRS family protein (RN15_RS04610) |
| | 11.67 | 3492272-3492219 | -1 | 1 | hypothetical protein (RN15_RS16475) |
| | 10.14 | 3521601-3521654 | 1 | 1 | AmmeMemoRadiSam system radical SAM enzyme (RN15_RS16655) |
| | 15.35 | 4355116-4355188 | 1 | 2 | ESX-1 secretion-associated protein EspE (RN15_RS20455) |
| | 10.74 | 3205322-3205387 | 1 | 2 | cell surface protein (RN15_RS15275) |
| | 10.74 | 3538115-3538161 | 1 | 2 | NADH-quinone oxidoreductase subunit I (RN15_RS16730) |
| | 10.04 | 2812543-2812494 | -1 | 2 | carboxylesterase LipQ (RN15_RS13215) |

| Organism | Bit Score | ANTAR RNA hit co-ordinates | Strand | RNA Category | Annotation of the downstream gene (Locus Tag) |
|---|---|---|---|---|---|
| *Mycobacterium_tuberculosis_strain=37004* (NZ_CP010338.1) | 17.37 | 2182762-2182812 | 1 | 1 | monooxygenase (RN13_RS10115) |
| | 13.76 | 3207358-3207412 | 1 | 1 | hypothetical protein (RN13_RS15290) |
| | 11.95 | 972245-972191 | -1 | 1 | PE-PGRS family protein (RN13_RS04585) |
| | 11.67 | 3466344-3466291 | -1 | 1 | hypothetical protein (RN13_RS16360) |
| | 10.14 | 3497435-3497488 | 1 | 1 | AmmeMemoRadiSam system radical SAM enzyme (RN13_RS16545) |
| | 15.35 | 4345713-4345785 | 1 | 2 | ESX-1 secretion-associated protein EspE (RN13_RS20450) |
| | 10.74 | 3179523-3179588 | 1 | 2 | cell surface protein (RN13_RS15150) |
| | 10.74 | 3513951-3513997 | 1 | 2 | NADH-quinone oxidoreductase subunit I (RN13_RS16620) |
| | 10.04 | 2793119-2793070 | -1 | 2 | carboxylesterase LipQ (RN13_RS13100) |
| *Mycobacterium_tuberculosis_strain=96075* (NZ_CP009426.1) | 17.37 | 2167819-2167869 | 1 | 1 | monooxygenase (LJ70_RS10085) |
| | 13.76 | 3184918-3184972 | 1 | 1 | hypothetical protein (LJ70_RS15255) |
| | 11.95 | 970202-970148 | -1 | 1 | hypothetical protein (LJ70_RS04590) |
| | 11.67 | 3440746-3440693 | -1 | 1 | hypothetical protein (LJ70_RS16290) |
| | 10.14 | 3473587-3473640 | 1 | 1 | AmmeMemoRadiSam system radical SAM enzyme (LJ70_RS16490) |
| | 15.35 | 4307966-4308038 | 1 | 2 | ESX-1 secretion-associated protein EspE (LJ70_RS20315) |
| | 10.74 | 3156975-3157040 | 1 | 2 | cell surface protein (LJ70_RS15115) |
| | 10.74 | 3490103-3490149 | 1 | 2 | NADH-quinone oxidoreductase subunit I (LJ70_RS16565) |
| | 10.04 | 2776849-2776800 | -1 | 2 | carboxylesterase (LJ70_RS13095) |
| *Mycobacterium_tuberculosis_strain=96121* (NZ_CP009427.1) | 17.37 | 2195453-2195503 | 1 | 1 | monooxygenase (HR37_RS10225) |
| | 13.76 | 3222862-3222916 | 1 | 1 | hypothetical protein (HR37_RS15420) |
| | 11.95 | 967101-967047 | -1 | 1 | hypothetical protein (HR37_RS04595) |
| | 11.67 | 3479189-3479136 | -1 | 1 | hypothetical protein (HR37_RS16465) |
| | 10.14 | 3509779-3509832 | 1 | 1 | AmmeMemoRadiSam system radical SAM enzyme (HR37_RS16645) |
| | 15.35 | 4339484-4339556 | 1 | 2 | ESX-1 secretion-associated protein EspE (HR37_RS20440) |
| | 10.74 | 3194915-3194980 | 1 | 2 | cell surface protein (HR37_RS15275) |
| | 10.74 | 3526295-3526341 | 1 | 2 | NADH-quinone oxidoreductase subunit I (HR37_RS16720) |
| | 10.04 | 2805515-2805466 | -1 | 2 | carboxylesterase LipQ (HR37_RS13230) |
| *Mycobacterium_tuberculosis_strain=F28* (NZ_CP010330.1) | 17.37 | 2194218-2194268 | 1 | 1 | monooxygenase (RN05_RS10215) |
| | 13.76 | 3223546-3223600 | 1 | 1 | hypothetical protein (RN05_RS15415) |
| | 11.95 | 970224-970170 | -1 | 1 | hypothetical protein (RN05_RS04600) |
| | 11.67 | 3482207-3482154 | -1 | 1 | hypothetical protein (RN05_RS16475) |
| | 10.14 | 3513299-3513352 | 1 | 1 | AmmeMemoRadiSam system radical SAM enzyme (RN05_RS16660) |
| | 15.35 | 4350537-4350609 | 1 | 2 | ESX-1 secretion-associated protein EspE (RN05_RS20465) |

| Organism | Bit Score | ANTAR RNA hit co-ordinates | Strand | RNA Category | Annotation of the downstream gene (Locus Tag) |
|---|---|---|---|---|---|
| | 10.74 | 3195709-3195774 | 1 | 2 | cell surface protein (RN05_RS15270) |
| | 10.74 | 3529815-3529861 | 1 | 2 | NADH-quinone oxidoreductase subunit I (RN05_RS16735) |
| | 10.04 | 2803074-2803025 | -1 | 2 | carboxylesterase LipQ (RN05_RS13210) |
| *Mycobacterium_tuberculosis_strain=KIT87190* (NZ_CP007809.1) | 17.37 | 2166335-2166385 | 1 | 1 | monooxygenase (ER17_RS10060) |
| | 13.76 | 3198994-3199048 | 1 | 1 | hypothetical protein (ER17_RS15265) |
| | 11.95 | 962146-962092 | -1 | 1 | hypothetical protein (ER17_RS04580) |
| | 11.67 | 3460996-3460943 | -1 | 1 | hypothetical protein (ER17_RS16355) |
| | 10.14 | 3492285-3492338 | 1 | 1 | AmmeMemoRadiSam system radical SAM enzyme (ER17_RS16530) |
| | 15.35 | 4339323-4339395 | 1 | 2 | ESX-1 secretion-associated protein EspE (ER17_RS20365) |
| | 10.74 | 3171105-3171170 | 1 | 2 | cell surface protein (ER17_RS15120) |
| | 10.74 | 3508801-3508847 | 1 | 2 | NADH-quinone oxidoreductase subunit I (ER17_RS16605) |
| | 10.04 | 2786510-2786461 | -1 | 2 | carboxylesterase (ER17_RS13105) |
| *Mycobacterium_tuberculosis_strain=SCAID_1870* (NZ_CP012506.1) | 17.37 | 2160992-2161042 | 1 | 1 | monooxygenase (AFL40_RS10020) |
| | 13.76 | 3184602-3184656 | 1 | 1 | hypothetical protein (AFL40_RS15160) |
| | 11.95 | 961490-961436 | -1 | 1 | hypothetical protein (AFL40_RS04560) |
| | 11.67 | 3442653-3442600 | -1 | 1 | hypothetical protein (AFL40_RS16220) |
| | 10.14 | 3473744-3473797 | 1 | 1 | AmmeMemoRadiSam system radical SAM enzyme (AFL40_RS16405) |
| | 15.35 | 4308106-4308178 | 1 | 2 | ESX-1 secretion-associated protein EspE (AFL40_RS20200) |
| | 10.74 | 3156765-3156830 | 1 | 2 | cell surface protein (AFL40_RS15020) |
| | 10.74 | 3490251-3490297 | 1 | 2 | NADH-quinone oxidoreductase subunit I (AFL40_RS16480) |
| | 10.04 | 2770281-2770232 | -1 | 2 | carboxylesterase (AFL40_RS12990) |
| *Mycobacterium_tuberculosis_strain=ZMC13-264* (NZ_CP009100.1) | 17.37 | 2187257-2187307 | 1 | 1 | monooxygenase (IZ84_RS10135) |
| | 13.76 | 3214554-3214608 | 1 | 1 | hypothetical protein (IZ84_RS15310) |
| | 11.95 | 970278-970224 | -1 | 1 | PE-PGRS family protein (IZ84_RS04580) |
| | 11.67 | 3473031-3472978 | -1 | 1 | hypothetical protein (IZ84_RS16370) |
| | 10.14 | 3504124-3504177 | 1 | 1 | AmmeMemoRadiSam system radical SAM enzyme (IZ84_RS16555) |
| | 15.35 | 4340137-4340209 | 1 | 2 | ESX-1 secretion-associated protein EspE (IZ84_RS20345) |
| | 10.74 | 3186717-3186782 | 1 | 2 | cell surface protein (IZ84_RS15165) |
| | 10.74 | 3520640-3520686 | 1 | 2 | NADH-quinone oxidoreductase subunit I (IZ84_RS16630) |
| | 10.04 | 2794094-2794045 | -1 | 2 | carboxylesterase (IZ84_RS13115) |
| *Mycobacterium_tuberculosis_strain=ZMC13-88* (NZ_CP009101.1) | 17.37 | 2187289-2187339 | 1 | 1 | monooxygenase (JE53_RS10145) |
| | 13.76 | 3214565-3214619 | 1 | 1 | hypothetical protein (JE53_RS15315) |
| | 11.95 | 970283-970229 | -1 | 1 | hypothetical protein (JE53_RS04595) |

| Organism | Bit Score | ANTAR RNA hit co-ordinates | Strand | RNA Category | Annotation of the downstream gene (Locus Tag) |
|---|---|---|---|---|---|
| | 11.67 | 3473042-3472989 | -1 | 1 | hypothetical protein (JE53_RS16375) |
| | 10.14 | 3504136-3504189 | 1 | 1 | AmmeMemoRadiSam system radical SAM enzyme (JE53_RS16560) |
| | 15.35 | 4340145-4340217 | 1 | 2 | ESX-1 secretion-associated protein EspE (JE53_RS20350) |
| | 10.74 | 3186728-3186793 | 1 | 2 | cell surface protein (JE53_RS15170) |
| | 10.74 | 3520652-3520698 | 1 | 2 | NADH-quinone oxidoreductase subunit I (JE53_RS16635) |
| | 10.04 | 2794105-2794056 | -1 | 2 | carboxylesterase (JE53_RS13120) |
| *Mycobacterium_tuberculosis_W-148* (NZ_CP012090.1) | 17.37 | 1954309-1954359 | 1 | 1 | monooxygenase (TBPG_RS08905) |
| | 13.76 | 2983060-2983114 | 1 | 1 | hypothetical protein (TBPG_RS14075) |
| | 11.95 | 3443793-3443847 | 1 | 1 | hypothetical protein (TBPG_RS16140) |
| | 11.67 | 946737-946790 | 1 | 1 | hypothetical protein (TBPG_RS04385) |
| | 10.14 | 913896-913843 | -1 | 1 | AmmeMemoRadiSam system radical SAM enzyme (TBPG_RS04185) |
| | 15.35 | 4347078-4347150 | 1 | 2 | ESX-1 secretion-associated protein EspE (TBPG_RS20360) |
| | 10.74 | 2955117-2955182 | 1 | 2 | cell surface protein (TBPG_RS13935) |
| | 10.74 | 897380-897334 | -1 | 2 | NADH-quinone oxidoreductase subunit I (TBPG_RS04110) |
| | 10.04 | 2568488-2568439 | -1 | 2 | carboxylesterase (TBPG_RS11910) |

*Table S3: Alignment of mycobacterial ANTAR RNA motifs predicted using Infernal.*
Structure based alignment of ANTAR RNAs obtained using Infernal. Alignment edited using RALEE v0.8 (PMID: 15377506). The stems are shaded in pink and blue.

| **Accession/Start-End** | **Sequence** |
|---|---|
| NC_008726/6033904-6033960 | .AGAGAGAUCACCAC.AAAGGU.GUGGUUG.......................ACUACACGGGCAAAGUGGCCGGUUUUCGGG |
| NZ_CP011491/5794851-5794907 | .AAAGAGAUCGCCAC.AAAGGU.GUGGUUG.......................ACUACACGGGCACAGUGGCCGUUCGUGGGA |
| NC_008726/4556520-4556572 | .UAACUAUAGUGCAC.AUGGAU.GUGCG--.......................-AUAAAUGAGCAAAGU-GCUUACAAAAUAC |
| NC_021054/2187067-2187117 | .ACAAAUUCCGAUGC.AAAGAA.GCACUU-.......................-----GUGUACAACGAAGUAUCUUGGUAGU |
| NC_009338/1209116-1209171 | .UGAGAGAUCACCAC.AAAGGU.GUGGUUG.......................-ACUACAGGGCAAUUGGGCCGGACUCGGGG |
| NC_014814/5213312-5213367 | .UGAGAGAUCACCAC.AAAGGU.GUGGUUG.......................-ACUACAGGGCAAUUGGGCCGGACUCGGGG |
| NC_015758/2187796-2187846 | .ACAAAUUCCCAUGC.AAAGAA.GCACUU-.......................-----GUGUACAACGAAGUAUCUUGGUAGU |
| NZ_CP003494/1889021-1889071 | .ACAAAUUCCCAUGC.AAAGAA.GCACUU-.......................-----GUGUACAACGAAGUAUCUUGGUAGU |
| NC_020245/2159066-2159116 | .ACAAAUUCCCAUGC.AAAGAA.GCACUU-.......................-----GUGUACAACGAAGUAUCUUGGUAGU |
| NC_016804/2168786-2168836 | .ACAAAUUCCCAUGC.AAAGAA.GCACUU-.......................-----GUGUACAACGAAGUAUCUUGGUAGU |
| NZ_AM412059/2171338-2171388 | .ACAAAUUCCCAUGC.AAAGAA.GCACUU-.......................-----GUGUACAACGAAGUAUCUUGGUAGU |
| NC_008769/2189421-2189471 | .ACAAAUUCCCAUGC.AAAGAA.GCACUU-.......................-----GUGUACAACGAAGUAUCUUGGUAGU |
| NC_012207/2171412-2171462 | .ACAAAUUCCCAUGC.AAAGAA.GCACUU-.......................-----GUGUACAACGAAGUAUCUUGGUAGU |
| NZ_CP014566/2171410-2171460 | .ACAAAUUCCCAUGC.AAAGAA.GCACUU-.......................-----GUGUACAACGAAGUAUCUUGGUAGU |
| NZ_CP008744/2169564-2169614 | .ACAAAUUCCCAUGC.AAAGAA.GCACUU-.......................-----GUGUACAACGAAGUAUCUUGGUAGU |
| NZ_CP009243/2171415-2171465 | .ACAAAUUCCCAUGC.AAAGAA.GCACUU-.......................-----GUGUACAACGAAGUAUCUUGGUAGU |
| NZ_CP012095/2183860-2183910 | .ACAAAUUCCCAUGC.AAAGAA.GCACUU-.......................-----GUGUACAACGAAGUAUCUUGGUAGU |
| NZ_CP009449/2147621-2147671 | .ACAAAUUCCCAUGC.AAAGAA.GCACUU-.......................-----GUGUACAACGAAGUAUCUUGGUAGU |
| NZ_CP013741/2171711-2171761 | .ACAAAUUCCCAUGC.AAAGAA.GCACUU-.......................-----GUGUACAACGAAGUAUCUUGGUAGU |
| NZ_HG813240/2173566-2173616 | .ACAAAUUCCCAUGC.AAAGAA.GCACUU-.......................-----GUGUACAACGAAGUAUCUUGGUAGU |
| NC_020089/2191699-2191749 | .ACAAAUUCCCAUGC.AAAGAA.GCACUU-.......................-----GUGUACAACGAAGUAUCUUGGUAGU |
| NZ_CP002883/2173137-2173187 | .ACAAAUUCCCAUGC.AAAGAA.GCACUU-.......................-----GUGUACAACGAAGUAUCUUGGUAGU |
| NZ_CP002882/2166350-2166400 | .ACAAAUUCCCAUGC.AAAGAA.GCACUU-.......................-----GUGUACAACGAAGUAUCUUGGUAGU |
| NC_017522/2168763-2168813 | .ACAAAUUCCCAUGC.AAAGAA.GCACUU-.......................-----GUGUACAACGAAGUAUCUUGGUAGU |
| NZ_CP002885/2173333-2173383 | .ACAAAUUCCCAUGC.AAAGAA.GCACUU-.......................-----GUGUACAACGAAGUAUCUUGGUAGU |
| NC_017524/2180824-2180874 | .ACAAAUUCCCAUGC.AAAGAA.GCACUU-.......................-----GUGUACAACGAAGUAUCUUGGUAGU |
| NC_021740/2181844-2181894 | .ACAAAUUCCCAUGC.AAAGAA.GCACUU-.......................-----GUGUACAACGAAGUAUCUUGGUAGU |
| NC_021194/2181480-2181530 | .ACAAAUUCCCAUGC.AAAGAA.GCACUU-.......................-----GUGUACAACGAAGUAUCUUGGUAGU |
| NC_009565/2195516-2195566 | .ACAAAUUCCCAUGC.AAAGAA.GCACUU-.......................-----GUGUACAACGAAGUAUCUUGGUAGU |
| NC_009525/2197160-2197210 | .ACAAAUUCCCAUGC.AAAGAA.GCACUU-.......................-----GUGUACAACGAAGUAUCUUGGUAGU |
| NC_000962/2187261-2187311 | .ACAAAUUCCCAUGC.AAAGAA.GCACUU-.......................-----GUGUACAACGAAGUAUCUUGGUAGU |
| NC_018143/2187268-2187318 | .ACAAAUUCCCAUGC.AAAGAA.GCACUU-.......................-----GUGUACAACGAAGUAUCUUGGUAGU |
| NZ_CP009480/2182565-2182615 | .ACAAAUUCCCAUGC.AAAGAA.GCACUU-.......................-----GUGUACAACGAAGUAUCUUGGUAGU |
| NZ_CP007027/2186649-2186699 | .ACAAAUUCCCAUGC.AAAGAA.GCACUU-.......................-----GUGUACAACGAAGUAUCUUGGUAGU |
| NZ_CP002871/2172407-2172457 | .ACAAAUUCCCAUGC.AAAGAA.GCACUU-.......................-----GUGUACAACGAAGUAUCUUGGUAGU |
| NC_012943/2235762-2235812 | .ACAAAUUCCCAUGC.AAAGAA.GCACUU-.......................-----GUGUACAACGAAGUAUCUUGGUAGU |
| NC_016768/2232493-2232543 | .ACAAAUUCCCAUGC.AAAGAA.GCACUU-.......................-----GUGUACAACGAAGUAUCUUGGUAGU |
| NC_018078/2235643-2235693 | .ACAAAUUCCCAUGC.AAAGAA.GCACUU-.......................-----GUGUACAACGAAGUAUCUUGGUAGU |
| NC_020559/2178334-2178384 | .ACAAAUUCCCAUGC.AAAGAA.GCACUU-.......................-----GUGUACAACGAAGUAUCUUGGUAGU |
| NC_022350/2186892-2186942 | .ACAAAUUCCCAUGC.AAAGAA.GCACUU-.......................-----GUGUACAACGAAGUAUCUUGGUAGU |
| NZ_AP014573/2192698-2192748 | .ACAAAUUCCCAUGC.AAAGAA.GCACUU-.......................-----GUGUACAACGAAGUAUCUUGGUAGU |
| NZ_CP010339/2185332-2185382 | .ACAAAUUCCCAUGC.AAAGAA.GCACUU-.......................-----GUGUACAACGAAGUAUCUUGGUAGU |
| NZ_CP010337/2190192-2190242 | .ACAAAUUCCCAUGC.AAAGAA.GCACUU-.......................-----GUGUACAACGAAGUAUCUUGGUAGU |
| NZ_CP010340/2199709-2199759 | .ACAAAUUCCCAUGC.AAAGAA.GCACUU-.......................-----GUGUACAACGAAGUAUCUUGGUAGU |
| NZ_CP010338/2182762-2182812 | .ACAAAUUCCCAUGC.AAAGAA.GCACUU-.......................-----GUGUACAACGAAGUAUCUUGGUAGU |
| NZ_CP009426/2167819-2167869 | .ACAAAUUCCCAUGC.AAAGAA.GCACUU-.......................-----GUGUACAACGAAGUAUCUUGGUAGU |
| NZ_CP009427/2195453-2195503 | .ACAAAUUCCCAUGC.AAAGAA.GCACUU-.......................-----GUGUACAACGAAGUAUCUUGGUAGU |
| NZ_CP010330/2194218-2194268 | .ACAAAUUCCCAUGC.AAAGAA.GCACUU-.......................-----GUGUACAACGAAGUAUCUUGGUAGU |
| NZ_CP007809/2166335-2166385 | .ACAAAUUCCCAUGC.AAAGAA.GCACUU-.......................-----GUGUACAACGAAGUAUCUUGGUAGU |
| NZ_CP012506/2160992-2161042 | .ACAAAUUCCCAUGC.AAAGAA.GCACUU-.......................-----GUGUACAACGAAGUAUCUUGGUAGU |
| NZ_CP009100/2187257-2187307 | .ACAAAUUCCCAUGC.AAAGAA.GCACUU-.......................-----GUGUACAACGAAGUAUCUUGGUAGU |
| NZ_CP009101/2187289-2187339 | .ACAAAUUCCCAUGC.AAAGAA.GCACUU-.......................-----GUGUACAACGAAGUAUCUUGGUAGU |
| NZ_CP012090/1954309-1954359 | .ACAAAUUCCCAUGC.AAAGAA.GCACUU-.......................-----GUGUACAACGAAGUAUCUUGGUAGU |
| NZ_CP012150/5895882-5895932 | .ACAGUGCGCAGCGC.AUCGGC.GCGCU--.......................----GCUGCGCAACGACGCGCAGAUGAUGA |
| NC_016604/761068-761121 | .AGUCGCUGUGCCUC.AACGUC.GAGGU--.......................-GUUGUCGAGCGACGGUGCUGCGAUCGAAA |
| NC_021251/2174974-2175038 | .AAGCACUUGUGUAC.AACGAA.GUAUCUUggua..............guacuGUGAUAUACGCAAAGG-GCGCCACCGCAGC |
| NC_002755/2184611-2184675 | .AAGCACUUGUGUAC.AACGAA.GUAUCUUggua..............guacuGUGAUAUACGCAAAGG-GCGCCACCGCAGC |
| NZ_CP007803/2159824-2159888 | .AAGCACUUGUGUAC.AACGAA.GUAUCUUggua..............guacuGUGAUAUACGCAAAGG-GCGCCACCGCAGC |
| NZ_CP011773/2186394-2186461 | .AAGGUUCGAUGCGC.AACGGC.GCGUACCcccug............accgcgGCCCAGAAAGCGAUGGUGCUCGGCAUGAAA |
| NC_016604/3909900-3909952 | .UGACUAUAGUGCAC.AUGGAC.GUGCG--.......................-AUAAAUGAGCAUAGU-GCUUACAAAAUAC |
| NZ_CP011269/6251522-6251576 | .GAGUUCUUAGUUAC.AAAGGU.GUAGUU-.......................-CGAUUGCGGCGACGGUGCCGUAGGGAGUG |
| NZ_CP009914/6321128-6321182 | .GAGUUCUUAGUUAC.AAAGGU.GUAGUU-.......................-CGAUUGCGGCGACGGUGCCGUAGGGAGUG |
| NC_010397/4098456-4098501 | .UAAGACAGUUCUAC.AAGGAG.GAAG---.......................--------AGCAAUGGCGCUCGUACUGGAA |
| NZ_CP014952/3992049-3992094 | .UAAGACAGUUCUAC.AAGGAG.GAAG---.......................--------AGCAAUGGCGCUCGUACUGGAA |
| NZ_CP014955/4105776-4105821 | .UAAGACAGUUCUAC.AAGGAG.GAAG---.......................--------AGCAAUGGCGCUCGUACUGGAA |
| NZ_CP014956/4220765-4220810 | .UAAGACAGUUCUAC.AAGGAG.GAAG---.......................--------AGCAAUGGCGCUCGUACUGGAA |
| NZ_CP014957/4178190-4178235 | .UAAGACAGUUCUAC.AAGGAG.GAAG---.......................--------AGCAAUGGCGCUCGUACUGGAA |

NZ_CP014960/3952564-3952609 .UAAGACAGUUCUAC.AAGGAG.GAAG---.......................--------AGCAAUGGCGCUCGUACUGGAA
NZ_CP013049/4103482-4103527 .UAAGACAGUUCUAC.AAGGAG.GAAG---.......................--------AGCAAUGGCGCUCGUACUGGAA
NZ_CP009407/4082751-4082796 .UAAGACAGUUCUAC.AAGGAG.GAAG---.......................--------AGCAAUGGCGCUCGUACUGGAA
NZ_CP009408/4097597-4097642 .UAAGACAGUUCUAC.AAGGAG.GAAG---.......................--------AGCAAUGGCGCUCGUACUGGAA
NZ_CP009613/4080661-4080706 .UAAGACAGUUCUAC.AAGGAG.GAAG---.......................--------AGCAAUGGCGCUCGUACUGGAA
NZ_CP012044/4393959-4394004 .UAAGACAGUUCUAC.AAGGAG.GAAG---.......................--------AGCAAUGGCGCUCGUACUGGAA
NC_023036/4181172-4181233 .AGCUUGCACGGAUC.AAUGGC.GAUCCUGuu..................agcCUUUUGAUCACAAAGGUGUGUAACAAAUCA
NZ_CP011022/4181243-4181304 .AGCUUGCACGGAUC.AAUGGC.GAUCCUGuu..................agcCUUUUGAUCACAAAGGUGUGUAACAAAUCA
NC_015758/3192504-3192558 .UUGGCGUGACACCC.AACAGA.GGGCAC.........................UUAAGAGGGCAAUGCGGCCGCCUACCUGC
NZ_CP003494/2878418-2878472 .UUGGCGUGACACCC.AACAGA.GGGCAC.........................UUAAGAGGGCAAUGCGGCCGCCUACCUGC
NC_020245/3138116-3138170 .UUGGCGUGACACCC.AACAGA.GGGCAC.........................UUAAGAGGGCAAUGCGGCCGCCUACCUGC
NC_016804/3147833-3147887 .UUGGCGUGACACCC.AACAGA.GGGCAC.........................UUAAGAGGGCAAUGCGGCCGCCUACCUGC
NZ_AM412059/3161017-3161071 .UUGGCGUGACACCC.AACAGA.GGGCAC.........................UUAAGAGGGCAAUGCGGCCGCCUACCUGC
NC_008769/3168468-3168522 .UUGGCGUGACACCC.AACAGA.GGGCAC.........................UUAAGAGGGCAAUGCGGCCGCCUACCUGC
NC_012207/3161295-3161349 .UUGGCGUGACACCC.AACAGA.GGGCAC.........................UUAAGAGGGCAAUGCGGCCGCCUACCUGC
NZ_CP014566/3161295-3161349 .UUGGCGUGACACCC.AACAGA.GGGCAC.........................UUAAGAGGGCAAUGCGGCCGCCUACCUGC
NZ_CP008744/3147951-3148005 .UUGGCGUGACACCC.AACAGA.GGGCAC.........................UUAAGAGGGCAAUGCGGCCGCCUACCUGC
NZ_CP009243/3161300-3161354 .UUGGCGUGACACCC.AACAGA.GGGCAC.........................UUAAGAGGGCAAUGCGGCCGCCUACCUGC
NZ_CP012095/3175544-3175598 .UUGGCGUGACACCC.AACAGA.GGGCAC.........................UUAAGAGGGCAAUGCGGCCGCCUACCUGC
NZ_CP009449/3123485-3123539 .UUGGCGUGACACCC.AACAGA.GGGCAC.........................UUAAGAGGGCAAUGCGGCCGCCUACCUGC
NZ_CP013741/3161523-3161577 .UUGGCGUGACACCC.AACAGA.GGGCAC.........................UUAAGAGGGCAAUGCGGCCGCCUACCUGC
NZ_CP015773/2077959-2078013 .UUGGCGUGACACCC.AACAGA.GGGCAC.........................UUAAGAGGGCAAUGCGGCCGCCUACCUGC
NC_015848/3273444-3273498 .UUGGCGUGACACCC.AACAGA.GGGCAC.........................UUAAGAGGGCAAUGCGGCCGCCUACCUGC
NZ_HG813240/3202634-3202688 .UUGGCGUGACACCC.AACAGA.GGGCAC.........................UUAAGAGGGCAAUGCGGCCGCCUACCUGC
NC_020089/3211910-3211964 .UUGGCGUGACACCC.AACAGA.GGGCAC.........................UUAAGAGGGCAAUGCGGCCGCCUACCUGC
NZ_CP002883/3196721-3196775 .UUGGCGUGACACCC.AACAGA.GGGCAC.........................UUAAGAGGGCAAUGCGGCCGCCUACCUGC
NZ_CP002882/3190985-3191039 .UUGGCGUGACACCC.AACAGA.GGGCAC.........................UUAAGAGGGCAAUGCGGCCGCCUACCUGC
NC_021251/3203172-3203226 .UUGGCGUGACACCC.AACAGA.GGGCAC.........................UUAAGAGGGCAAUGCGGCCGCCUACCUGC
NC_017522/3195483-3195537 .UUGGCGUGACACCC.AACAGA.GGGCAC.........................UUAAGAGGGCAAUGCGGCCGCCUACCUGC
NZ_CP002885/3200058-3200112 .UUGGCGUGACACCC.AACAGA.GGGCAC.........................UUAAGAGGGCAAUGCGGCCGCCUACCUGC
NC_002755/3208690-3208744 .UUGGCGUGACACCC.AACAGA.GGGCAC.........................UUAAGAGGGCAAUGCGGCCGCCUACCUGC
NC_017524/3210750-3210804 .UUGGCGUGACACCC.AACAGA.GGGCAC.........................UUAAGAGGGCAAUGCGGCCGCCUACCUGC
NC_021740/3200965-3201019 .UUGGCGUGACACCC.AACAGA.GGGCAC.........................UUAAGAGGGCAAUGCGGCCGCCUACCUGC
NC_021194/3200395-3200449 .UUGGCGUGACACCC.AACAGA.GGGCAC.........................UUAAGAGGGCAAUGCGGCCGCCUACCUGC
NC_009565/3226065-3226119 .UUGGCGUGACACCC.AACAGA.GGGCAC.........................UUAAGAGGGCAAUGCGGCCGCCUACCUGC
NC_009525/3226528-3226582 .UUGGCGUGACACCC.AACAGA.GGGCAC.........................UUAAGAGGGCAAUGCGGCCGCCUACCUGC
NC_018143/3214546-3214600 .UUGGCGUGACACCC.AACAGA.GGGCAC.........................UUAAGAGGGCAAUGCGGCCGCCUACCUGC
NZ_CP009480/3207732-3207786 .UUGGCGUGACACCC.AACAGA.GGGCAC.........................UUAAGAGGGCAAUGCGGCCGCCUACCUGC
NZ_CP007027/3213924-3213978 .UUGGCGUGACACCC.AACAGA.GGGCAC.........................UUAAGAGGGCAAUGCGGCCGCCUACCUGC
NZ_CP002871/3199246-3199300 .UUGGCGUGACACCC.AACAGA.GGGCAC.........................UUAAGAGGGCAAUGCGGCCGCCUACCUGC
NZ_CP007803/3189638-3189692 .UUGGCGUGACACCC.AACAGA.GGGCAC.........................UUAAGAGGGCAAUGCGGCCGCCUACCUGC
NC_012943/1199238-1199292 .UUGGCGUGACACCC.AACAGA.GGGCAC.........................UUAAGAGGGCAAUGCGGCCGCCUACCUGC
NC_016768/1199236-1199290 .UUGGCGUGACACCC.AACAGA.GGGCAC.........................UUAAGAGGGCAAUGCGGCCGCCUACCUGC
NC_018078/1199350-1199404 .UUGGCGUGACACCC.AACAGA.GGGCAC.........................UUAAGAGGGCAAUGCGGCCGCCUACCUGC
NC_021054/3214228-3214282 .UUGGCGUGACACCC.AACAGA.GGGCAC.........................UUAAGAGGGCAAUGCGGCCGCCUACCUGC
NC_020559/3203020-3203074 .UUGGCGUGACACCC.AACAGA.GGGCAC.........................UUAAGAGGGCAAUGCGGCCGCCUACCUGC
NC_022350/3209954-3210008 .UUGGCGUGACACCC.AACAGA.GGGCAC.........................UUAAGAGGGCAAUGCGGCCGCCUACCUGC
NZ_AP014573/3220427-3220481 .UUGGCGUGACACCC.AACAGA.GGGCAC.........................UUAAGAGGGCAAUGCGGCCGCCUACCUGC
NZ_CP010339/3212420-3212474 .UUGGCGUGACACCC.AACAGA.GGGCAC.........................UUAAGAGGGCAAUGCGGCCGCCUACCUGC
NZ_CP010337/3221258-3221312 .UUGGCGUGACACCC.AACAGA.GGGCAC.........................UUAAGAGGGCAAUGCGGCCGCCUACCUGC
NZ_CP010340/3233211-3233265 .UUGGCGUGACACCC.AACAGA.GGGCAC.........................UUAAGAGGGCAAUGCGGCCGCCUACCUGC
NZ_CP010338/3207358-3207412 .UUGGCGUGACACCC.AACAGA.GGGCAC.........................UUAAGAGGGCAAUGCGGCCGCCUACCUGC
NZ_CP009426/3184918-3184972 .UUGGCGUGACACCC.AACAGA.GGGCAC.........................UUAAGAGGGCAAUGCGGCCGCCUACCUGC
NZ_CP009427/3222862-3222916 .UUGGCGUGACACCC.AACAGA.GGGCAC.........................UUAAGAGGGCAAUGCGGCCGCCUACCUGC
NZ_CP010330/3223546-3223600 .UUGGCGUGACACCC.AACAGA.GGGCAC.........................UUAAGAGGGCAAUGCGGCCGCCUACCUGC
NZ_CP007809/3198994-3199048 .UUGGCGUGACACCC.AACAGA.GGGCAC.........................UUAAGAGGGCAAUGCGGCCGCCUACCUGC
NZ_CP012506/3184602-3184656 .UUGGCGUGACACCC.AACAGA.GGGCAC.........................UUAAGAGGGCAAUGCGGCCGCCUACCUGC
NZ_CP009100/3214554-3214608 .UUGGCGUGACACCC.AACAGA.GGGCAC.........................UUAAGAGGGCAAUGCGGCCGCCUACCUGC
NZ_CP009101/3214565-3214619 .UUGGCGUGACACCC.AACAGA.GGGCAC.........................UUAAGAGGGCAAUGCGGCCGCCUACCUGC
NZ_CP012090/2983060-2983114 .UUGGCGUGACACCC.AACAGA.GGGCAC.........................UUAAGAGGGCAAUGCGGCCGCCUACCUGC
NC_000962/3214536-3214590 .UUGGCGUGACACCC.AACAGA.GGGCAC.........................UUAAGAGGGCAAUGCGGCCGCCUACCUGC
NZ_CP015773/1085811-1085858 .----------GUAC.AACGAA.GUAUUUUggua..............guacuGUGAUAUACGCAAAGG-GCGCCA-------
NC_010397/1789584-1789640 .AUUGCACAAGUCGA.AAAGGA.UCGGGCC.......................AAGGAAGAGGCGAUGGCGCCACAAGUAGCG
NZ_CP014955/1772773-1772829 .AUUGCACAAGUCGA.AAAGGA.UCGGGCC.......................AAGGAAGAGGCGAUGGCGCCACAAGUAGCG
NZ_CP009407/1774108-1774164 .AUUGCACAAGUCGA.AAAGGA.UCGGGCC.......................AAGGAAGAGGCGAUGGCGCCACAAGUAGCG
NZ_CP009408/1789073-1789129 .AUUGCACAAGUCGA.AAAGGA.UCGGGCC.......................AAGGAAGAGGCGAUGGCGCCACAAGUAGCG
NZ_CP009613/1781082-1781138 .AUUGCACAAGUCGA.AAAGGA.UCGGGCC.......................AAGGAAGAGGCGAUGGCGCCACAAGUAGCG
NZ_CP014475/1931132-1931198 .AAAGACCCUAGGUC.AACGGU.GCUCUUCggcuu.............acgguGGAACCGUCGCAACGGAGCGCAAAGGAGAA
NC_011896/567138-567198 .CGUGUCUUGGCGCC.AUCGAU.GGCGU-ggu..................ggugGGUAUUGGGCAACCGAGCCUCAAAAACUC
NC_009077/2700344-2700396 .AACAUCUGUUCCGC.AUUAGG.GUGG---.......................-AUCGGACCGCAAGGGUGCGGACGACGAAA
NC_008705/2714603-2714655 .AACAUCUGUUCCGC.AUUAGG.GUGG---.......................-AUCGGACCGCAAGGGUGCGGACGACGAAA
NC_008146/2696715-2696767 .AACAUCUGUUCCGC.AUUAGG.GUGG---.......................-AUCGGACCGCAAGGGUGCGGACGACGAAA

NC_019958/29757-29809 .UCAACGAAGAAGAC.AUCGAC.GUCUU--......................--CUUGAACGCAUUGUCGCGACAAGGCAGC

```
[illegible]
NC_011896/3088433-3088494 .ACACAACCAGCUAU.AGAGCC.GUGGCAGac..................cauAAGAGAAGAGCCAAGGUGCUUUGUGAGGCA
NZ_CP011530/5249432-5249478 .UAUGCAGUUAGUAC.AACGCU.GUGCA--.......................--------UGCAAAGAUGCACGGGAGCGCA
NC_015758/966587-966641 .ACAGCCGAUCGGUC.AACGAA.GACCCU.........................GCGGGAGGACAGCGAUGUCGUAUGUAUUG
NZ_CP003494/688279-688333 .ACAGCCGAUCGGUC.AACGAA.GACCCU.........................GCGGGAGGACAGCGAUGUCGUAUGUAUUG
NC_020245/971023-971077 .ACAGCCGAUCGGUC.AACGAA.GACCCU.........................GCGGGAGGACAGCGAUGUCGUAUGUAUUG
NC_016804/970969-971023 .ACAGCCGAUCGGUC.AACGAA.GACCCU.........................GCGGGAGGACAGCGAUGUCGUAUGUAUUG
NZ_AM412059/972326-972380 .ACAGCCGAUCGGUC.AACGAA.GACCCU.........................GCGGGAGGACAGCGAUGUCGUAUGUAUUG
NC_008769/1000690-1000744 .ACAGCCGAUCGGUC.AACGAA.GACCCU.........................GCGGGAGGACAGCGAUGUCGUAUGUAUUG
NC_012207/972407-972461 .ACAGCCGAUCGGUC.AACGAA.GACCCU.........................GCGGGAGGACAGCGAUGUCGUAUGUAUUG
NZ_CP014566/972407-972461 .ACAGCCGAUCGGUC.AACGAA.GACCCU.........................GCGGGAGGACAGCGAUGUCGUAUGUAUUG
NZ_CP008744/971035-971089 .ACAGCCGAUCGGUC.AACGAA.GACCCU.........................GCGGGAGGACAGCGAUGUCGUAUGUAUUG
NZ_CP009243/972404-972458 .ACAGCCGAUCGGUC.AACGAA.GACCCU.........................GCGGGAGGACAGCGAUGUCGUAUGUAUUG
NZ_CP012095/970750-970804 .ACAGCCGAUCGGUC.AACGAA.GACCCU.........................GCGGGAGGACAGCGAUGUCGUAUGUAUUG
NZ_CP013741/972329-972383 .ACAGCCGAUCGGUC.AACGAA.GACCCU.........................GCGGGAGGACAGCGAUGUCGUAUGUAUUG
NZ_HG813240/968674-968728 .ACAGCCGAUCGGUC.AACGAA.GACCCU.........................GCGGGAGGACAGCGAUGUCGUAUGUAUUG
NC_020089/973544-973598 .ACAGCCGAUCGGUC.AACGAA.GACCCU.........................GCGGGAGGACAGCGAUGUCGUAUGUAUUG
NZ_CP002883/968988-969042 .ACAGCCGAUCGGUC.AACGAA.GACCCU.........................GCGGGAGGACAGCGAUGUCGUAUGUAUUG
NZ_CP002882/968933-968987 .ACAGCCGAUCGGUC.AACGAA.GACCCU.........................GCGGGAGGACAGCGAUGUCGUAUGUAUUG
NC_021251/968795-968849 .ACAGCCGAUCGGUC.AACGAA.GACCCU.........................GCGGGAGGACAGCGAUGUCGUAUGUAUUG
NZ_CP002885/968749-968803 .ACAGCCGAUCGGUC.AACGAA.GACCCU.........................GCGGGAGGACAGCGAUGUCGUAUGUAUUG
NC_002755/970243-970297 .ACAGCCGAUCGGUC.AACGAA.GACCCU.........................GCGGGAGGACAGCGAUGUCGUAUGUAUUG
NC_017524/970646-970700 .ACAGCCGAUCGGUC.AACGAA.GACCCU.........................GCGGGAGGACAGCGAUGUCGUAUGUAUUG
NC_021740/968881-968935 .ACAGCCGAUCGGUC.AACGAA.GACCCU.........................GCGGGAGGACAGCGAUGUCGUAUGUAUUG
NC_021194/968698-968752 .ACAGCCGAUCGGUC.AACGAA.GACCCU.........................GCGGGAGGACAGCGAUGUCGUAUGUAUUG
NC_009565/974054-974108 .ACAGCCGAUCGGUC.AACGAA.GACCCU.........................GCGGGAGGACAGCGAUGUCGUAUGUAUUG
NC_009525/971536-971590 .ACAGCCGAUCGGUC.AACGAA.GACCCU.........................GCGGGAGGACAGCGAUGUCGUAUGUAUUG
NC_018143/970233-970287 .ACAGCCGAUCGGUC.AACGAA.GACCCU.........................GCGGGAGGACAGCGAUGUCGUAUGUAUUG
NZ_CP009480/967284-967338 .ACAGCCGAUCGGUC.AACGAA.GACCCU.........................GCGGGAGGACAGCGAUGUCGUAUGUAUUG
NZ_CP007027/970233-970287 .ACAGCCGAUCGGUC.AACGAA.GACCCU.........................GCGGGAGGACAGCGAUGUCGUAUGUAUUG
NZ_CP002871/968754-968808 .ACAGCCGAUCGGUC.AACGAA.GACCCU.........................GCGGGAGGACAGCGAUGUCGUAUGUAUUG
NZ_CP007803/963531-963585 .ACAGCCGAUCGGUC.AACGAA.GACCCU.........................GCGGGAGGACAGCGAUGUCGUAUGUAUUG
NC_012943/3441695-3441749 .ACAGCCGAUCGGUC.AACGAA.GACCCU.........................GCGGGAGGACAGCGAUGUCGUAUGUAUUG
NC_016768/3438428-3438482 .ACAGCCGAUCGGUC.AACGAA.GACCCU.........................GCGGGAGGACAGCGAUGUCGUAUGUAUUG
NC_018078/3441449-3441503 .ACAGCCGAUCGGUC.AACGAA.GACCCU.........................GCGGGAGGACAGCGAUGUCGUAUGUAUUG
NC_020559/967726-967780 .ACAGCCGAUCGGUC.AACGAA.GACCCU.........................GCGGGAGGACAGCGAUGUCGUAUGUAUUG
NC_022350/973809-973863 .ACAGCCGAUCGGUC.AACGAA.GACCCU.........................GCGGGAGGACAGCGAUGUCGUAUGUAUUG
NZ_CP009426/970148-970202 .ACAGCCGAUCGGUC.AACGAA.GACCCU.........................GCGGGAGGACAGCGAUGUCGUAUGUAUUG
NZ_CP009427/967047-967101 .ACAGCCGAUCGGUC.AACGAA.GACCCU.........................GCGGGAGGACAGCGAUGUCGUAUGUAUUG
NZ_CP010330/970170-970224 .ACAGCCGAUCGGUC.AACGAA.GACCCU.........................GCGGGAGGACAGCGAUGUCGUAUGUAUUG
NZ_CP007809/962092-962146 .ACAGCCGAUCGGUC.AACGAA.GACCCU.........................GCGGGAGGACAGCGAUGUCGUAUGUAUUG
NZ_CP012506/961436-961490 .ACAGCCGAUCGGUC.AACGAA.GACCCU.........................GCGGGAGGACAGCGAUGUCGUAUGUAUUG
NZ_CP009101/970229-970283 .ACAGCCGAUCGGUC.AACGAA.GACCCU.........................GCGGGAGGACAGCGAUGUCGUAUGUAUUG
NZ_CP012090/3443793-3443847 .ACAGCCGAUCGGUC.AACGAA.GACCCU.........................GCGGGAGGACAGCGAUGUCGUAUGUAUUG
NZ_AP014573/974330-974384 .ACAGCCGAUCGGUC.AACGAA.GACCCU.........................GCGGGAGGACAGCGAUGUCGUAUGUAUUG
NZ_CP009449/964113-964167 .ACAGCCGAUCGGUC.AACGAA.GACCCU.........................GCGGGAGGACAGCGAUGUCGUAUGUAUUG
NZ_CP015773/4221290-4221344 .ACAGCCGAUCGGUC.AACGAA.GACCCU.........................GCGGGAGGACAGCGAUGUCGUAUGUAUUG
NC_017522/966381-966435 .ACAGCCGAUCGGUC.AACGAA.GACCCU.........................GCGGGAGGACAGCGAUGUCGUAUGUAUUG
NC_021054/970138-970192 .ACAGCCGAUCGGUC.AACGAA.GACCCU.........................GCGGGAGGACAGCGAUGUCGUAUGUAUUG
NZ_CP010339/972442-972496 .ACAGCCGAUCGGUC.AACGAA.GACCCU.........................GCGGGAGGACAGCGAUGUCGUAUGUAUUG
NZ_CP010337/972743-972797 .ACAGCCGAUCGGUC.AACGAA.GACCCU.........................GCGGGAGGACAGCGAUGUCGUAUGUAUUG
NZ_CP010340/977020-977074 .ACAGCCGAUCGGUC.AACGAA.GACCCU.........................GCGGGAGGACAGCGAUGUCGUAUGUAUUG
NZ_CP010338/972191-972245 .ACAGCCGAUCGGUC.AACGAA.GACCCU.........................GCGGGAGGACAGCGAUGUCGUAUGUAUUG
NZ_CP009100/970224-970278 .ACAGCCGAUCGGUC.AACGAA.GACCCU.........................GCGGGAGGACAGCGAUGUCGUAUGUAUUG
NC_000962/970230-970284 .ACAGCCGAUCGGUC.AACGAA.GACCCU.........................GCGGGAGGACAGCGAUGUCGUAUGUAUUG
NZ_CP011491/3023516-3023552 .----------GUAC.AGUGGC.GUACCG-.......................-GAACCCGAGCGAAGGAGCUUC--------
NC_015758/3451978-3452031 .CAGAACAAGAUCGC.GGCGAU.GCGGGU.........................UCUGC-AAGCAAAGUUGUUGGAACGCAAG
NZ_CP003494/3137258-3137311 .CAGAACAAGAUCGC.GGCGAU.GCGGGU.........................UCUGC-AAGCAAAGUUGUUGGAACGCAAG
NC_020245/3396765-3396818 .CAGAACAAGAUCGC.GGCGAU.GCGGGU.........................UCUGC-AAGCAAAGUUGUUGGAACGCAAG
NC_016804/3406835-3406888 .CAGAACAAGAUCGC.GGCGAU.GCGGGU.........................UCUGC-AAGCAAAGUUGUUGGAACGCAAG
NZ_AM412059/3418917-3418970 .CAGAACAAGAUCGC.GGCGAU.GCGGGU.........................UCUGC-AAGCAAAGUUGUUGGAACGCAAG
NC_008769/3427470-3427523 .CAGAACAAGAUCGC.GGCGAU.GCGGGU.........................UCUGC-AAGCAAAGUUGUUGGAACGCAAG
NC_012207/3420120-3420173 .CAGAACAAGAUCGC.GGCGAU.GCGGGU.........................UCUGC-AAGCAAAGUUGUUGGAACGCAAG
NZ_CP014566/3420120-3420173 .CAGAACAAGAUCGC.GGCGAU.GCGGGU.........................UCUGC-AAGCAAAGUUGUUGGAACGCAAG
NZ_CP008744/3406776-3406829 .CAGAACAAGAUCGC.GGCGAU.GCGGGU.........................UCUGC-AAGCAAAGUUGUUGGAACGCAAG
NZ_CP009243/3420125-3420178 .CAGAACAAGAUCGC.GGCGAU.GCGGGU.........................UCUGC-AAGCAAAGUUGUUGGAACGCAAG
NZ_CP012095/3434465-3434518 .CAGAACAAGAUCGC.GGCGAU.GCGGGU.........................UCUGC-AAGCAAAGUUGUUGGAACGCAAG
NZ_CP009449/3382103-3382156 .CAGAACAAGAUCGC.GGCGAU.GCGGGU.........................UCUGC-AAGCAAAGUUGUUGGAACGCAAG
NZ_CP013741/3420525-3420578 .CAGAACAAGAUCGC.GGCGAU.GCGGGU.........................UCUGC-AAGCAAAGUUGUUGGAACGCAAG
NZ_CP015773/2336559-2336612 .CAGAACAAGAUCGC.GGCGAU.GCGGGU.........................UCUGC-AAGCAAAGUUGUUGGAACGCAAG
NC_015848/3537144-3537197 .CAGAACAAGAUCGC.GGCGAU.GCGGGU.........................UCUGC-AAGCAAAGUUGUUGGAACGCAAG
NZ_HG813240/3462191-3462244 .CAGAACAAGAUCGC.GGCGAU.GCGGGU.........................UCUGC-AAGCAAAGUUGUUGGAACGCAAG
NC_020089/3468856-3468909 .CAGAACAAGAUCGC.GGCGAU.GCGGGU.........................UCUGC-AAGCAAAGUUGUUGGAACGCAAG
```

| | |
|---|---|
| [illegible] | [illegible] |
| NZ_CP002883/3456777-3456830 | .CAGAACAAGAUCGC.GGCGAU.GCGGGU.........................UCUGC-AAGCAAAGUUGUUGGAACGCAAG |
| NZ_CP002882/3451049-3451102 | .CAGAACAAGAUCGC.GGCGAU.GCGGGU.........................UCUGC-AAGCAAAGUUGUUGGAACGCAAG |
| NC_021251/3463234-3463287 | .CAGAACAAGAUCGC.GGCGAU.GCGGGU.........................UCUGC-AAGCAAAGUUGUUGGAACGCAAG |
| NC_017522/3456820-3456873 | .CAGAACAAGAUCGC.GGCGAU.GCGGGU.........................UCUGC-AAGCAAAGUUGUUGGAACGCAAG |
| NZ_CP002885/3462008-3462061 | .CAGAACAAGAUCGC.GGCGAU.GCGGGU.........................UCUGC-AAGCAAAGUUGUUGGAACGCAAG |
| NC_017524/3468769-3468822 | .CAGAACAAGAUCGC.GGCGAU.GCGGGU.........................UCUGC-AAGCAAAGUUGUUGGAACGCAAG |
| NC_021740/3459411-3459464 | .CAGAACAAGAUCGC.GGCGAU.GCGGGU.........................UCUGC-AAGCAAAGUUGUUGGAACGCAAG |
| NC_021194/3458761-3458814 | .CAGAACAAGAUCGC.GGCGAU.GCGGGU.........................UCUGC-AAGCAAAGUUGUUGGAACGCAAG |
| NC_009565/3484748-3484801 | .CAGAACAAGAUCGC.GGCGAU.GCGGGU.........................UCUGC-AAGCAAAGUUGUUGGAACGCAAG |
| NC_009525/3485140-3485193 | .CAGAACAAGAUCGC.GGCGAU.GCGGGU.........................UCUGC-AAGCAAAGUUGUUGGAACGCAAG |
| NC_018143/3472981-3473034 | .CAGAACAAGAUCGC.GGCGAU.GCGGGU.........................UCUGC-AAGCAAAGUUGUUGGAACGCAAG |
| NZ_CP009480/3466129-3466182 | .CAGAACAAGAUCGC.GGCGAU.GCGGGU.........................UCUGC-AAGCAAAGUUGUUGGAACGCAAG |
| NZ_CP007027/3472366-3472419 | .CAGAACAAGAUCGC.GGCGAU.GCGGGU.........................UCUGC-AAGCAAAGUUGUUGGAACGCAAG |
| NZ_CP002871/3461023-3461076 | .CAGAACAAGAUCGC.GGCGAU.GCGGGU.........................UCUGC-AAGCAAAGUUGUUGGAACGCAAG |
| NZ_CP007803/3449889-3449942 | .CAGAACAAGAUCGC.GGCGAU.GCGGGU.........................UCUGC-AAGCAAAGUUGUUGGAACGCAAG |
| NC_012943/940739-940792 | .CAGAACAAGAUCGC.GGCGAU.GCGGGU.........................UCUGC-AAGCAAAGUUGUUGGAACGCAAG |
| NC_016768/940742-940795 | .CAGAACAAGAUCGC.GGCGAU.GCGGGU.........................UCUGC-AAGCAAAGUUGUUGGAACGCAAG |
| NC_018078/940744-940797 | .CAGAACAAGAUCGC.GGCGAU.GCGGGU.........................UCUGC-AAGCAAAGUUGUUGGAACGCAAG |
| NC_021054/3472645-3472698 | .CAGAACAAGAUCGC.GGCGAU.GCGGGU.........................UCUGC-AAGCAAAGUUGUUGGAACGCAAG |
| NC_020559/3459248-3459301 | .CAGAACAAGAUCGC.GGCGAU.GCGGGU.........................UCUGC-AAGCAAAGUUGUUGGAACGCAAG |
| NZ_AP014573/3479331-3479384 | .CAGAACAAGAUCGC.GGCGAU.GCGGGU.........................UCUGC-AAGCAAAGUUGUUGGAACGCAAG |
| NZ_CP010339/3471431-3471484 | .CAGAACAAGAUCGC.GGCGAU.GCGGGU.........................UCUGC-AAGCAAAGUUGUUGGAACGCAAG |
| NZ_CP010337/3481952-3482005 | .CAGAACAAGAUCGC.GGCGAU.GCGGGU.........................UCUGC-AAGCAAAGUUGUUGGAACGCAAG |
| NZ_CP010340/3492219-3492272 | .CAGAACAAGAUCGC.GGCGAU.GCGGGU.........................UCUGC-AAGCAAAGUUGUUGGAACGCAAG |
| NZ_CP010338/3466291-3466344 | .CAGAACAAGAUCGC.GGCGAU.GCGGGU.........................UCUGC-AAGCAAAGUUGUUGGAACGCAAG |
| NZ_CP009426/3440693-3440746 | .CAGAACAAGAUCGC.GGCGAU.GCGGGU.........................UCUGC-AAGCAAAGUUGUUGGAACGCAAG |
| NZ_CP009427/3479136-3479189 | .CAGAACAAGAUCGC.GGCGAU.GCGGGU.........................UCUGC-AAGCAAAGUUGUUGGAACGCAAG |
| NZ_CP010330/3482154-3482207 | .CAGAACAAGAUCGC.GGCGAU.GCGGGU.........................UCUGC-AAGCAAAGUUGUUGGAACGCAAG |
| NZ_CP007809/3460943-3460996 | .CAGAACAAGAUCGC.GGCGAU.GCGGGU.........................UCUGC-AAGCAAAGUUGUUGGAACGCAAG |
| NZ_CP012506/3442600-3442653 | .CAGAACAAGAUCGC.GGCGAU.GCGGGU.........................UCUGC-AAGCAAAGUUGUUGGAACGCAAG |
| NZ_CP009100/3472978-3473031 | .CAGAACAAGAUCGC.GGCGAU.GCGGGU.........................UCUGC-AAGCAAAGUUGUUGGAACGCAAG |
| NZ_CP009101/3472989-3473042 | .CAGAACAAGAUCGC.GGCGAU.GCGGGU.........................UCUGC-AAGCAAAGUUGUUGGAACGCAAG |
| NZ_CP012090/946737-946790 | .CAGAACAAGAUCGC.GGCGAU.GCGGGU.........................UCUGC-AAGCAAAGUUGUUGGAACGCAAG |
| NC_022350/3466701-3466754 | .CAGAACAAGAUCGC.GGCGAU.GCGGGU.........................UCUGC-AAGCAAAGUUGUUGGAACGCAAG |
| NC_000962/3472984-3473037 | .CAGAACAAGAUCGC.GGCGAU.GCGGGU.........................UCUGC-AAGCAAAGUUGUUGGAACGCAAG |
| NC_002755/3468780-3468832 | .CAGAACAAGAUCGC.GGCGAU.GCGGGU.........................UCUGC-AAGCAAAGUUGUUGGAACG-CAA |
| NZ_CP011530/14770-14818 | .AAAGGGUGGAACCG.AAUGGC.CGGUC--.......................------CAUGCGUCGGAGCACAAUAAGAAG |
| NZ_CP014957/4646038-4646086 | .AUAGAUCG...CGGCCAGAUGCUCG..............................GUCGGCGCAAUGGUGCGCAGAAAGAUG |
| NC_018612/2593620-2593675 | .GAUUCGAGUAGCCC.AACGUC.GCGCUU-.......................CCAGGUUCAGCAACGCCGCUCUUCAACGUG |
| NC_009338/2744015-2744062 | .----------GCAC.ACUGGU.GAGCUCGagag...............caggACUCAUAGGGCAAUGACGCCUUU------- |
| NC_014814/2104142-2104189 | .----------GCAC.ACUGGU.GAGCUCGagag...............caggACUCAUAGGGCAAUGACGCCUUU------- |
| NZ_CP011883/173986-174049 | .AUUCCCAGAUCUAC.GAUGGU.GUGGCUUcau.................caacGAUGCCGGUGCAACGGUGCGCCGGGAACG |
| NZ_CP009914/2975165-2975225 | .UCCUGGUACcgCGC.AGUGU-.GCG..................gguacguggACUUGAAAUGCAUUGUGGCGCAGUAUGAUU |
| NZ_AP012555/2832394-2832453 | .UAGAGCCAGGCGGC.ACAGGU.GCCGUCGg.....................ccGAGGUAAAAGUGAUGUCGCUUCGGGUCAUG |
| NC_008596/3682246-3682309 | .GUGAUCCCCGGUAC.AUGGAG.GUACAUcaug................gcggaGGUGCUGGUCACCGGCGGCGACACCGAAC |
| NC_018289/3683859-3683922 | .GUGAUCCCCGGUAC.AUGGAG.GUACAUcaug................gcggaGGUGCUGGUCACCGGCGGCGACACCGAAC |
| NZ_CP009494/3682286-3682349 | .GUGAUCCCCGGUAC.AUGGAG.GUACAUcaug................gcggaGGUGCUGGUCACCGGCGGCGACACCGAAC |
| NZ_CP009495/3682320-3682383 | .GUGAUCCCCGGUAC.AUGGAG.GUACAUcaug................gcggaGGUGCUGGUCACCGGCGGCGACACCGAAC |
| NZ_CP009496/3682308-3682371 | .GUGAUCCCCGGUAC.AUGGAG.GUACAUcaug................gcggaGGUGCUGGUCACCGGCGGCGACACCGAAC |
| NZ_LN831039/3744254-3744317 | .GUGAUCCCCGGUAC.AUGGAG.GUACAUcaug................gcggaGGUGCUGGUCACCGGCGGCGACACCGAAC |
| NC_016604/5093059-5093112 | .UCUUCAACGAGCGC.AAAGUC.GCGGA--.......................-GCUGAAGGUCAAAGAAGCCAGCUAUUGGA |
| NC_020133/3552989-3553040 | .GUUAUACCGACGGC.AGAGUG.GUCGU--.......................---GCCUUUGCAAAGGCGCAGCCAACUAAG |
| NZ_HG917972/2705617-2705668 | .GUUAUACCGACGGC.AGAGUG.GUCGU--.......................---GCCUUUGCAAAGGCGCAGCCAACUAAG |
| NC_010612/2842712-2842763 | .GUUAUACCGACGGC.AGAGUG.GUCGU--.......................---GCCUUUGCAAAGGCGCAGCCAACUAAG |
| NC_009077/1434237-1434280 | .----------CCAC.AUCGAA.GUGGUUCgcc...............gaagAUAUGUUUGGCAAUGACGUC---------- |
| NC_008705/1416437-1416480 | .----------CCAC.AUCGAA.GUGGUUCgcc...............gaagAUAUGUUUGGCAAUGACGUC---------- |
| NC_008146/1412128-1412171 | .----------CCAC.AUCGAA.GUGGUUCgcc...............gaagAUAUGUUUGGCAAUGACGUC---------- |
| NZ_CP014950/3909717-3909764 | .--AUAAGACAGUUC.GUACAA.GGA..............................GGAAGAGCAAUGGCGCUCGUACUGGAA |
| NZ_CP014959/4078623-4078670 | .--AUAAGACAGUUC.GUACAA.GGA..............................GGAAGAGCAAUGGCGCUCGUACUGGAA |
| NZ_CP014961/4460783-4460830 | .--AUAAGACAGUUC.GUACAA.GGA..............................GGAAGAGCAAUGGCGCUCGUACUGGAA |
| NZ_CP009914/5424084-5424140 | .AACUUCUACCAGAC.AGCGGU.GUCGUGC.......................AUGGACACGGCGUGGAAGCCCCUGAUCACU |
| NC_015758/3485225-3485278 | .ACGGUCUGUUGCAC.GACGAG.GUGCU-..........................GACACGGCUCAACGCCGGGUAACCUGGCG |
| NZ_CP003494/3167077-3167130 | .ACGGUCUGUUGCAC.GACGAG.GUGCU-..........................GACACGGCUCAACGCCGGGUAACCUGGCG |
| NC_020245/3426471-3426524 | .ACGGUCUGUUGCAC.GACGAG.GUGCU-..........................GACACGGCUCAACGCCGGGUAACCUGGCG |
| NC_016804/3436541-3436594 | .ACGGUCUGUUGCAC.GACGAG.GUGCU-..........................GACACGGCUCAACGCCGGGUAACCUGGCG |
| NZ_AM412059/3448623-3448676 | .ACGGUCUGUUGCAC.GACGAG.GUGCU-..........................GACACGGCUCAACGCCGGGUAACCUGGCG |
| NC_008769/3457176-3457229 | .ACGGUCUGUUGCAC.GACGAG.GUGCU-..........................GACACGGCUCAACGCCGGGUAACCUGGCG |
| NC_012207/3449826-3449879 | .ACGGUCUGUUGCAC.GACGAG.GUGCU-..........................GACACGGCUCAACGCCGGGUAACCUGGCG |
| NZ_CP014566/3449826-3449879 | .ACGGUCUGUUGCAC.GACGAG.GUGCU-..........................GACACGGCUCAACGCCGGGUAACCUGGCG |
| NZ_CP008744/3436482-3436535 | .ACGGUCUGUUGCAC.GACGAG.GUGCU-..........................GACACGGCUCAACGCCGGGUAACCUGGCG |
| NZ_CP009243/3449831-3449884 | .ACGGUCUGUUGCAC.GACGAG.GUGCU-..........................GACACGGCUCAACGCCGGGUAACCUGGCG |
| NZ_CP012095/3464172-3464225 | .ACGGUCUGUUGCAC.GACGAG.GUGCU-..........................GACACGGCUCAACGCCGGGUAACCUGGCG |

NZ_CP009449/3411810-3411863 .ACGGUCUGUUGCAC.GACGAG.GUGCU-........................GACACGGCUCAACGCCGGGUAACCUGGCG
NZ_CP013741/3450231-3450284 .ACGGUCUGUUGCAC.GACGAG.GUGCU-........................GACACGGCUCAACGCCGGGUAACCUGGCG
NZ_CP015773/2366266-2366319 .ACGGUCUGUUGCAC.GACGAG.GUGCU-........................GACACGGCUCAACGCCGGGUAACCUGGCG
NC_015848/3559166-3559219 .ACGGUCUGUUGCAC.GACGAG.GUGCU-........................GACACGGCUCAACGCCGGGUAACCUGGCG
NZ_HG813240/3495085-3495138 .ACGGUCUGUUGCAC.GACGAG.GUGCU-........................GACACGGCUCAACGCCGGGUAACCUGGCG
NC_020089/3501049-3501102 .ACGGUCUGUUGCAC.GACGAG.GUGCU-........................GACACGGCUCAACGCCGGGUAACCUGGCG
NZ_CP002883/3489671-3489724 .ACGGUCUGUUGCAC.GACGAG.GUGCU-........................GACACGGCUCAACGCCGGGUAACCUGGCG
NZ_CP002882/3483943-3483996 .ACGGUCUGUUGCAC.GACGAG.GUGCU-........................GACACGGCUCAACGCCGGGUAACCUGGCG
NC_021251/3496128-3496181 .ACGGUCUGUUGCAC.GACGAG.GUGCU-........................GACACGGCUCAACGCCGGGUAACCUGGCG
NC_017522/3489714-3489767 .ACGGUCUGUUGCAC.GACGAG.GUGCU-........................GACACGGCUCAACGCCGGGUAACCUGGCG
NZ_CP002885/3494902-3494955 .ACGGUCUGUUGCAC.GACGAG.GUGCU-........................GACACGGCUCAACGCCGGGUAACCUGGCG
NC_002755/3499614-3499667 .ACGGUCUGUUGCAC.GACGAG.GUGCU-........................GACACGGCUCAACGCCGGGUAACCUGGCG
NC_017524/3501273-3501326 .ACGGUCUGUUGCAC.GACGAG.GUGCU-........................GACACGGCUCAACGCCGGGUAACCUGGCG
NC_021740/3490553-3490606 .ACGGUCUGUUGCAC.GACGAG.GUGCU-........................GACACGGCUCAACGCCGGGUAACCUGGCG
NC_021194/3489881-3489934 .ACGGUCUGUUGCAC.GACGAG.GUGCU-........................GACACGGCUCAACGCCGGGUAACCUGGCG
NC_009565/3517252-3517305 .ACGGUCUGUUGCAC.GACGAG.GUGCU-........................GACACGGCUCAACGCCGGGUAACCUGGCG
NC_009525/3516285-3516338 .ACGGUCUGUUGCAC.GACGAG.GUGCU-........................GACACGGCUCAACGCCGGGUAACCUGGCG
NC_018143/3504126-3504179 .ACGGUCUGUUGCAC.GACGAG.GUGCU-........................GACACGGCUCAACGCCGGGUAACCUGGCG
NZ_CP009480/3497274-3497327 .ACGGUCUGUUGCAC.GACGAG.GUGCU-........................GACACGGCUCAACGCCGGGUAACCUGGCG
NZ_CP007027/3503511-3503564 .ACGGUCUGUUGCAC.GACGAG.GUGCU-........................GACACGGCUCAACGCCGGGUAACCUGGCG
NZ_CP002871/3493917-3493970 .ACGGUCUGUUGCAC.GACGAG.GUGCU-........................GACACGGCUCAACGCCGGGUAACCUGGCG
NZ_CP007803/3482372-3482425 .ACGGUCUGUUGCAC.GACGAG.GUGCU-........................GACACGGCUCAACGCCGGGUAACCUGGCG
NC_012943/3504952-3505005 .ACGGUCUGUUGCAC.GACGAG.GUGCU-........................GACACGGCUCAACGCCGGGUAACCUGGCG
NC_016768/3501686-3501739 .ACGGUCUGUUGCAC.GACGAG.GUGCU-........................GACACGGCUCAACGCCGGGUAACCUGGCG
NC_018078/3506065-3506118 .ACGGUCUGUUGCAC.GACGAG.GUGCU-........................GACACGGCUCAACGCCGGGUAACCUGGCG
NC_021054/3503789-3503842 .ACGGUCUGUUGCAC.GACGAG.GUGCU-........................GACACGGCUCAACGCCGGGUAACCUGGCG
NC_020559/3490082-3490135 .ACGGUCUGUUGCAC.GACGAG.GUGCU-........................GACACGGCUCAACGCCGGGUAACCUGGCG
NC_022350/3497535-3497588 .ACGGUCUGUUGCAC.GACGAG.GUGCU-........................GACACGGCUCAACGCCGGGUAACCUGGCG
NZ_AP014573/3511835-3511888 .ACGGUCUGUUGCAC.GACGAG.GUGCU-........................GACACGGCUCAACGCCGGGUAACCUGGCG
NZ_CP010339/3503913-3503966 .ACGGUCUGUUGCAC.GACGAG.GUGCU-........................GACACGGCUCAACGCCGGGUAACCUGGCG
NZ_CP010337/3513096-3513149 .ACGGUCUGUUGCAC.GACGAG.GUGCU-........................GACACGGCUCAACGCCGGGUAACCUGGCG
NZ_CP010340/3521601-3521654 .ACGGUCUGUUGCAC.GACGAG.GUGCU-........................GACACGGCUCAACGCCGGGUAACCUGGCG
NZ_CP010338/3497435-3497488 .ACGGUCUGUUGCAC.GACGAG.GUGCU-........................GACACGGCUCAACGCCGGGUAACCUGGCG
NZ_CP009426/3473587-3473640 .ACGGUCUGUUGCAC.GACGAG.GUGCU-........................GACACGGCUCAACGCCGGGUAACCUGGCG
NZ_CP009427/3509779-3509832 .ACGGUCUGUUGCAC.GACGAG.GUGCU-........................GACACGGCUCAACGCCGGGUAACCUGGCG
NZ_CP010330/3513299-3513352 .ACGGUCUGUUGCAC.GACGAG.GUGCU-........................GACACGGCUCAACGCCGGGUAACCUGGCG
NZ_CP007809/3492285-3492338 .ACGGUCUGUUGCAC.GACGAG.GUGCU-........................GACACGGCUCAACGCCGGGUAACCUGGCG
NZ_CP012506/3473744-3473797 .ACGGUCUGUUGCAC.GACGAG.GUGCU-........................GACACGGCUCAACGCCGGGUAACCUGGCG
NZ_CP009100/3504124-3504177 .ACGGUCUGUUGCAC.GACGAG.GUGCU-........................GACACGGCUCAACGCCGGGUAACCUGGCG
NZ_CP009101/3504136-3504189 .ACGGUCUGUUGCAC.GACGAG.GUGCU-........................GACACGGCUCAACGCCGGGUAACCUGGCG
NZ_CP012090/913843-913896 .ACGGUCUGUUGCAC.GACGAG.GUGCU-........................GACACGGCUCAACGCCGGGUAACCUGGCG
NZ_CP011883/1112421-1112495 .AUUAGUUUUGCGGU.AAGGGC.GCUGAGGuguuagugu....aguugacacGUGUCAAGUGCAAGGGUGCACGUUCGCAUA
NC_000962/3504129-3504182 .ACGGUCUGUUGCAC.GACGAG.GUGCU-........................GACACGGCUCAACGCCGGGUAACCUGGCG
NC_011896/465129-465188 .ACUUCAAUCCGGCGCAACUGUACGUCU.....................guaGUUAAAUGCGCAACGACGCGAGCAUGGAAU
NC_015758/4317847-4317904 AACCCCCCUCCGAGCGCUGCUAAGCUC........................aAUGUGCAGUGCAAAGGUGCAGAUAAUGAUG
NZ_CP003494/3987382-3987439 AACCCCCCUCCGAGCGCUGCUAAGCUC........................aAUGUGCAGUGCAAAGGUGCAGAUAAUGAUG
NC_020245/4314752-4314809 AACCCCCCUCCGAGCGCUGCUAAGCUC........................aAUGUGCAGUGCAAAGGUGCAGAUAAUGAUG
NC_016804/4288427-4288484 AACCCCCCUCCGAGCGCUGCUAAGCUC........................aAUGUGCAGUGCAAAGGUGCAGAUAAUGAUG
NZ_AM412059/4279033-4279090 AACCCCCCUCCGAGCGCUGCUAAGCUC........................aAUGUGCAGUGCAAAGGUGCAGAUAAUGAUG
NC_008769/4312563-4312620 AACCCCCCUCCGAGCGCUGCUAAGCUC........................aAUGUGCAGUGCAAAGGUGCAGAUAAUGAUG
NC_012207/4309752-4309809 AACCCCCCUCCGAGCGCUGCUAAGCUC........................aAUGUGCAGUGCAAAGGUGCAGAUAAUGAUG
NZ_CP014566/4309748-4309805 AACCCCCCUCCGAGCGCUGCUAAGCUC........................aAUGUGCAGUGCAAAGGUGCAGAUAAUGAUG
NZ_CP008744/4334126-4334183 AACCCCCCUCCGAGCGCUGCUAAGCUC........................aAUGUGCAGUGCAAAGGUGCAGAUAAUGAUG
NZ_CP009243/4308179-4308236 AACCCCCCUCCGAGCGCUGCUAAGCUC........................aAUGUGCAGUGCAAAGGUGCAGAUAAUGAUG
NZ_CP012095/4282663-4282720 AACCCCCCUCCGAGCGCUGCUAAGCUC........................aAUGUGCAGUGCAAAGGUGCAGAUAAUGAUG
NZ_CP009449/4298241-4298298 AACCCCCCUCCGAGCGCUGCUAAGCUC........................aAUGUGCAGUGCAAAGGUGCAGAUAAUGAUG
NZ_CP013741/4308746-4308803 AACCCCCCUCCGAGCGCUGCUAAGCUC........................aAUGUGCAGUGCAAAGGUGCAGAUAAUGAUG
NZ_CP015773/3181083-3181140 AACCCCCCUCCGAGCGCUGCUAAGCUC........................aAUGUGCAGUGCAAAGGUGCAGAUAAUGAUG
NC_015848/4410435-4410492 AACCCCCCUCCGAGCGCUGCUAAGCUC........................aAUGUGCAGUGCAAAGGUGCAGAUAAUGAUG
NZ_HG813240/4340917-4340974 AACCCCCCUCCGAGCGCUGCUAAGCUC........................aAUGUGCAGUGCAAAGGUGCAGAUAAUGAUG
NC_020089/4349871-4349928 AACCCCCCUCCGAGCGCUGCUAAGCUC........................aAUGUGCAGUGCAAAGGUGCAGAUAAUGAUG
NZ_CP002883/4328038-4328095 AACCCCCCUCCGAGCGCUGCUAAGCUC........................aAUGUGCAGUGCAAAGGUGCAGAUAAUGAUG
NZ_CP002882/4330472-4330529 AACCCCCCUCCGAGCGCUGCUAAGCUC........................aAUGUGCAGUGCAAAGGUGCAGAUAAUGAUG
NC_021251/4342863-4342920 AACCCCCCUCCGAGCGCUGCUAAGCUC........................aAUGUGCAGUGCAAAGGUGCAGAUAAUGAUG
NC_017522/4334516-4334573 AACCCCCCUCCGAGCGCUGCUAAGCUC........................aAUGUGCAGUGCAAAGGUGCAGAUAAUGAUG
NZ_CP002885/4342884-4342941 AACCCCCCUCCGAGCGCUGCUAAGCUC........................aAUGUGCAGUGCAAAGGUGCAGAUAAUGAUG
NC_002755/4332485-4332542 AACCCCCCUCCGAGCGCUGCUAAGCUC........................aAUGUGCAGUGCAAAGGUGCAGAUAAUGAUG
NC_017524/4327151-4327208 AACCCCCCUCCGAGCGCUGCUAAGCUC........................aAUGUGCAGUGCAAAGGUGCAGAUAAUGAUG
NC_021740/4319795-4319852 AACCCCCCUCCGAGCGCUGCUAAGCUC........................aAUGUGCAGUGCAAAGGUGCAGAUAAUGAUG
NC_021194/4318936-4318993 AACCCCCCUCCGAGCGCUGCUAAGCUC........................aAUGUGCAGUGCAAAGGUGCAGAUAAUGAUG
NC_009565/4353162-4353219 AACCCCCCUCCGAGCGCUGCUAAGCUC........................aAUGUGCAGUGCAAAGGUGCAGAUAAUGAUG
NC_009525/4348608-4348665 AACCCCCCUCCGAGCGCUGCUAAGCUC........................aAUGUGCAGUGCAAAGGUGCAGAUAAUGAUG

| | |
|---|---|
| NC_000962/4340162-4340219 | AACCCCCCUCCGAGCGCUGCUAAGCUC.........................aAUGUGCAGUGCAAAGGUGCAGAUAAUGAUG |
| NC_018143/4340340-4340397 | AACCCCCCUCCGAGCGCUGCUAAGCUC.........................aAUGUGCAGUGCAAAGGUGCAGAUAAUGAUG |
| NZ_CP009480/4332571-4332628 | AACCCCCCUCCGAGCGCUGCUAAGCUC.........................aAUGUGCAGUGCAAAGGUGCAGAUAAUGAUG |
| NZ_CP007027/4339542-4339599 | AACCCCCCUCCGAGCGCUGCUAAGCUC.........................aAUGUGCAGUGCAAAGGUGCAGAUAAUGAUG |
| NZ_CP002871/4336467-4336524 | AACCCCCCUCCGAGCGCUGCUAAGCUC.........................aAUGUGCAGUGCAAAGGUGCAGAUAAUGAUG |
| NZ_CP007803/4314056-4314113 | AACCCCCCUCCGAGCGCUGCUAAGCUC.........................aAUGUGCAGUGCAAAGGUGCAGAUAAUGAUG |
| NC_012943/4327108-4327165 | AACCCCCCUCCGAGCGCUGCUAAGCUC.........................aAUGUGCAGUGCAAAGGUGCAGAUAAUGAUG |
| NC_016768/4323712-4323769 | AACCCCCCUCCGAGCGCUGCUAAGCUC.........................aAUGUGCAGUGCAAAGGUGCAGAUAAUGAUG |
| NC_018078/4327977-4328034 | AACCCCCCUCCGAGCGCUGCUAAGCUC.........................aAUGUGCAGUGCAAAGGUGCAGAUAAUGAUG |
| NC_021054/4339761-4339818 | AACCCCCCUCCGAGCGCUGCUAAGCUC.........................aAUGUGCAGUGCAAAGGUGCAGAUAAUGAUG |
| NC_020559/4321068-4321125 | AACCCCCCUCCGAGCGCUGCUAAGCUC.........................aAUGUGCAGUGCAAAGGUGCAGAUAAUGAUG |
| NC_022350/4337000-4337057 | AACCCCCCUCCGAGCGCUGCUAAGCUC.........................aAUGUGCAGUGCAAAGGUGCAGAUAAUGAUG |
| NZ_AP014573/4343757-4343814 | AACCCCCCUCCGAGCGCUGCUAAGCUC.........................aAUGUGCAGUGCAAAGGUGCAGAUAAUGAUG |
| NZ_CP010339/4328067-4328124 | AACCCCCCUCCGAGCGCUGCUAAGCUC.........................aAUGUGCAGUGCAAAGGUGCAGAUAAUGAUG |
| NZ_CP010337/4330406-4330463 | AACCCCCCUCCGAGCGCUGCUAAGCUC.........................aAUGUGCAGUGCAAAGGUGCAGAUAAUGAUG |
| NZ_CP010340/4355116-4355173 | AACCCCCCUCCGAGCGCUGCUAAGCUC.........................aAUGUGCAGUGCAAAGGUGCAGAUAAUGAUG |
| NZ_CP010338/4345713-4345770 | AACCCCCCUCCGAGCGCUGCUAAGCUC.........................aAUGUGCAGUGCAAAGGUGCAGAUAAUGAUG |
| NZ_CP009426/4307966-4308023 | AACCCCCCUCCGAGCGCUGCUAAGCUC.........................aAUGUGCAGUGCAAAGGUGCAGAUAAUGAUG |
| NZ_CP009427/4339484-4339541 | AACCCCCCUCCGAGCGCUGCUAAGCUC.........................aAUGUGCAGUGCAAAGGUGCAGAUAAUGAUG |
| NZ_CP010330/4350537-4350594 | AACCCCCCUCCGAGCGCUGCUAAGCUC.........................aAUGUGCAGUGCAAAGGUGCAGAUAAUGAUG |
| NZ_CP007809/4339323-4339380 | AACCCCCCUCCGAGCGCUGCUAAGCUC.........................aAUGUGCAGUGCAAAGGUGCAGAUAAUGAUG |
| NZ_CP012506/4308106-4308163 | AACCCCCCUCCGAGCGCUGCUAAGCUC.........................aAUGUGCAGUGCAAAGGUGCAGAUAAUGAUG |
| NZ_CP009100/4340137-4340194 | AACCCCCCUCCGAGCGCUGCUAAGCUC.........................aAUGUGCAGUGCAAAGGUGCAGAUAAUGAUG |
| NZ_CP009101/4340145-4340202 | AACCCCCCUCCGAGCGCUGCUAAGCUC.........................aAUGUGCAGUGCAAAGGUGCAGAUAAUGAUG |
| NZ_CP012090/4347078-4347135 | AACCCCCCUCCGAGCGCUGCUAAGCUC.........................aAUGUGCAGUGCAAAGGUGCAGAUAAUGAUG |
| NZ_CP011530/4673950-4674016 | .AUAUUUCUUCCCAC.AUAGAU.GUGGUUU.............uugacgauguguagggcGCGAAAAGCGUUGCccgacgacc |
| NZ_CP011491/1708817-1708870 | .AC---GAUCAGCGG.CAGCGC.CUGCUGG.......................AUGUGGUGCGCAAGGACGCGCCCUGGAAAU |
| NC_008595/2096449-2096505 | .CGACGAUUUgaCGC.CGGGC-.GUG......................cgucgCAAGCAAUCGCAUUGGUGCGCAGAGGGAGG |
| NZ_CP009614/1991098-1991154 | .CGACGAUUUgaCGC.CGGGC-.GUG......................cgucgCAAGCAAUCGCAUUGGUGCGCAGAGGGAGG |
| NZ_CP009493/1986373-1986429 | .CGACGAUUUgaCGC.CGGGC-.GUG......................cgucgCAAGCAAUCGCAUUGGUGCGCAGAGGGAGG |
| NZ_CP009482/1985968-1986024 | .CGACGAUUUgaCGC.CGGGC-.GUG......................cgucgCAAGCAAUCGCAUUGGUGCGCAGAGGGAGG |
| NZ_AP012555/1815297-1815353 | .CGACGAUUUgaCGC.CGGGC-.GUG......................cgucgCAAGCAAUCGCAUUGGUGCGCAGAGGGAGG |
| NZ_CP010113/1882988-1883044 | .CGACGAUUUgaCGC.CGGGC-.GUG......................cgucgCAAGCAAUCGCAUUGGUGCGCAGAGGGAGG |
| NZ_CP010114/1885164-1885220 | .CGACGAUUUgaCGC.CGGGC-.GUG......................cgucgCAAGCAAUCGCAUUGGUGCGCAGAGGGAGG |
| NC_021200/1902216-1902272 | .CGACGAUUUgaCGC.CGGGC-.GUG......................cgucgCAAGCAAUCGCAUUGGUGCGCAGAGGGAGG |
| NC_002944/2307024-2307080 | .CGACGAUUUgaCGC.CGGGC-.GUG......................cgucgCAAGCAAUCGCAUUGGUGCGCAGAGGGAGG |
| NZ_CP015495/2307024-2307080 | .CGACGAUUUgaCGC.CGGGC-.GUG......................cgucgCAAGCAAUCGCAUUGGUGCGCAGAGGGAGG |
| NC_014814/4314648-4314715 | .UCAGUUAAGACUAC.AAGGGU.GUAGUUUguugguaugcc............ucgcgcGGGUUGGCGGAAGCCUGAAUCGGA |
| NZ_CP014959/4510746-4510794 | .AUAGAUCG...CGGCCAGAUGCUCG..............................GUCGGCGCAAUGGUGCGCAGAAAGAUG |
| NZ_CP014961/4891304-4891352 | .AUAGAUCG...CGGCCAGAUGCUCG..............................GUCGGCGCAAUGGUGCGCAGAAAGAUG |
| NC_021282/4535069-4535117 | .AUAGAUCG...CGGCCAGAUGCUCG..............................GUCGGCGCAAUGGUGCGCAGAAAGAUG |
| NZ_AP014547/4549368-4549416 | .AUAGAUCG...CGGCCAGAUGCUCG..............................GUCGGCGCAAUGGUGCGCAGAAAGAUG |
| NZ_CP009447/405303-405351 | .AUAGAUCG...CGGCCAGAUGCUCG..............................GUCGGCGCAAUGGUGCGCAGAAAGAUG |
| NZ_CP014950/4379334-4379382 | .AUAGAUCG...CGGCCAGAUGCUCG..............................GUCGGCGCAAUGGUGCGCAGAAAGAUG |
| NC_018027/2095742-2095801 | GGACAAUAGaGCGCUAUUGGCCUGGCGU.......................ccUACCUUUGAACAUGGCAGUUUUGUAAAAAG |
| NZ_CP007220/3471896-3471949 | .AUGAUCUGGAGCAC.ACCGAU.GUGCU--.......................-CGUCGCGAGCGAAGGAGCCACGCCCCAUG |
| NZ_CP010071/3306365-3306418 | .AUGAUCUGGAGCAC.ACCGAU.GUGCU--.......................-CGUCGCGAGCGAAGGAGCCACGCCCCAUG |
| NC_020133/3257155-3257207 | .--AUUUACUUUAGC.UAACGU.UCUAGU-.......................-UGUCUAGCGCAACGUCGCGCGGUCAAAUU |
| NZ_HG917972/2438029-2438081 | .--AUUUACUUUAGC.UAACGU.UCUAGU-.......................-UGUCUAGCGCAACGUCGCGCGGUCAAAUU |
| NC_010612/2553269-2553321 | .--AUUUACUUUAGC.UAACGU.UCUAGU-.......................-UGUCUAGCGCAACGUCGCGCGGUCAAAUU |
| NC_008596/6497108-6497169 | .GAAUUCAUCGGCAC.AUCGAG.GUGCC-gcggacaggcc.................aacaCCGAUGUCCUGGAGGCAAAAUG |
| NC_018289/6498717-6498778 | .GAAUUCAUCGGCAC.AUCGAG.GUGCC-gcggacaggcc.................aacaCCGAUGUCCUGGAGGCAAAAUG |
| NZ_CP009494/6497148-6497209 | .GAAUUCAUCGGCAC.AUCGAG.GUGCC-gcggacaggcc.................aacaCCGAUGUCCUGGAGGCAAAAUG |
| NZ_CP009495/6497210-6497271 | .GAAUUCAUCGGCAC.AUCGAG.GUGCC-gcggacaggcc.................aacaCCGAUGUCCUGGAGGCAAAAUG |
| NZ_CP009496/6497179-6497240 | .GAAUUCAUCGGCAC.AUCGAG.GUGCC-gcggacaggcc.................aacaCCGAUGUCCUGGAGGCAAAAUG |
| NZ_LN831039/6514456-6514517 | .GAAUUCAUCGGCAC.AUCGAG.GUGCC-gcggacaggcc.................aacaCCGAUGUCCUGGAGGCAAAAUG |
| NZ_CP011530/3685270-3685323 | .AUGAUCUGGAGCAC.AUCGAC.GUGCU--.......................-UGCCGCGAGCGAAGGAGCGACGCCCCAUG |
| NC_018289/6012010-6012063 | .UCACUUUCGUGGGA.GGCCGG.UCC..........................ggUCAGACGGGCAAUGAGGCCGAGCAUCAGC |
| NZ_CP009494/6010442-6010495 | .UCACUUUCGUGGGA.GGCCGG.UCC..........................ggUCAGACGGGCAAUGAGGCCGAGCAUCAGC |
| NZ_CP009495/6010512-6010565 | .UCACUUUCGUGGGA.GGCCGG.UCC..........................ggUCAGACGGGCAAUGAGGCCGAGCAUCAGC |
| NZ_CP009496/6010480-6010533 | .UCACUUUCGUGGGA.GGCCGG.UCC..........................ggUCAGACGGGCAAUGAGGCCGAGCAUCAGC |
| NC_008596/6010399-6010452 | .UCACUUUCGUGGGA.GGCCGG.UCC..........................ggUCAGACGGGCAAUGAGGCCGAGCAUCAGC |
| NC_015758/3164610-3164675 | .CCACGUCCGACGGC.AACGAC.GCCGCUaucaa..............auugaaUUACCACGCCAAAGACGUCUACAUCGUUG |
| NZ_CP003494/2850691-2850756 | .CCACGUCCGACGGC.AACGAC.GCCGCUaucaa..............auugaaUUACCACGCCAAAGACGUCUACAUCGUUG |
| NC_020245/3110279-3110344 | .CCACGUCCGACGGC.AACGAC.GCCGCUaucaa..............auugaaUUACCACGCCAAAGACGUCUACAUCGUUG |
| NC_016804/3119996-3120061 | .CCACGUCCGACGGC.AACGAC.GCCGCUaucaa..............auugaaUUACCACGCCAAAGACGUCUACAUCGUUG |
| NZ_AM412059/3133180-3133245 | .CCACGUCCGACGGC.AACGAC.GCCGCUaucaa..............auugaaUUACCACGCCAAAGACGUCUACAUCGUUG |
| NC_008769/3140631-3140696 | .CCACGUCCGACGGC.AACGAC.GCCGCUaucaa..............auugaaUUACCACGCCAAAGACGUCUACAUCGUUG |
| NC_012207/3133458-3133523 | .CCACGUCCGACGGC.AACGAC.GCCGCUaucaa..............auugaaUUACCACGCCAAAGACGUCUACAUCGUUG |
| NZ_CP014566/3133458-3133523 | .CCACGUCCGACGGC.AACGAC.GCCGCUaucaa..............auugaaUUACCACGCCAAAGACGUCUACAUCGUUG |
| NZ_CP008744/3120114-3120179 | .CCACGUCCGACGGC.AACGAC.GCCGCUaucaa..............auugaaUUACCACGCCAAAGACGUCUACAUCGUUG |
| NZ_CP009243/3133463-3133528 | .CCACGUCCGACGGC.AACGAC.GCCGCUaucaa..............auugaaUUACCACGCCAAAGACGUCUACAUCGUUG |
| [illegible] | [illegible] |

| | |
|---|---|
| NZ_CP012095/3147760-3147825 | .CCACGUCCGACGGC.AACGAC.GCCGCUaucaa..............auugaaUUACCACGCCAAAGACGUCUACAUCGUUG |
| NZ_CP009449/3095648-3095713 | .CCACGUCCGACGGC.AACGAC.GCCGCUaucaa..............auugaaUUACCACGCCAAAGACGUCUACAUCGUUG |
| NZ_CP013741/3133686-3133751 | .CCACGUCCGACGGC.AACGAC.GCCGCUaucaa..............auugaaUUACCACGCCAAAGACGUCUACAUCGUUG |
| NZ_CP015773/2050122-2050187 | .CCACGUCCGACGGC.AACGAC.GCCGCUaucaa..............auugaaUUACCACGCCAAAGACGUCUACAUCGUUG |
| NC_015848/3245334-3245399 | .CCACGUCCGACGGC.AACGAC.GCCGCUaucaa..............auugaaUUACCACGCCAAAGACGUCUACAUCGUUG |
| NZ_HG813240/3174691-3174756 | .CCACGUCCGACGGC.AACGAC.GCCGCUaucaa..............auugaaUUACCACGCCAAAGACGUCUACAUCGUUG |
| NC_020089/3184160-3184225 | .CCACGUCCGACGGC.AACGAC.GCCGCUaucaa..............auugaaUUACCACGCCAAAGACGUCUACAUCGUUG |
| NZ_CP002883/3168827-3168892 | .CCACGUCCGACGGC.AACGAC.GCCGCUaucaa..............auugaaUUACCACGCCAAAGACGUCUACAUCGUUG |
| NZ_CP002882/3163042-3163107 | .CCACGUCCGACGGC.AACGAC.GCCGCUaucaa..............auugaaUUACCACGCCAAAGACGUCUACAUCGUUG |
| NC_021251/3175229-3175294 | .CCACGUCCGACGGC.AACGAC.GCCGCUaucaa..............auugaaUUACCACGCCAAAGACGUCUACAUCGUUG |
| NC_017522/3167540-3167605 | .CCACGUCCGACGGC.AACGAC.GCCGCUaucaa..............auugaaUUACCACGCCAAAGACGUCUACAUCGUUG |
| NZ_CP002885/3172115-3172180 | .CCACGUCCGACGGC.AACGAC.GCCGCUaucaa..............auugaaUUACCACGCCAAAGACGUCUACAUCGUUG |
| NC_002755/3180854-3180919 | .CCACGUCCGACGGC.AACGAC.GCCGCUaucaa..............auugaaUUACCACGCCAAAGACGUCUACAUCGUUG |
| NC_017524/3182967-3183032 | .CCACGUCCGACGGC.AACGAC.GCCGCUaucaa..............auugaaUUACCACGCCAAAGACGUCUACAUCGUUG |
| NC_021740/3173129-3173194 | .CCACGUCCGACGGC.AACGAC.GCCGCUaucaa..............auugaaUUACCACGCCAAAGACGUCUACAUCGUUG |
| NC_021194/3172580-3172645 | .CCACGUCCGACGGC.AACGAC.GCCGCUaucaa..............auugaaUUACCACGCCAAAGACGUCUACAUCGUUG |
| NC_009565/3198229-3198294 | .CCACGUCCGACGGC.AACGAC.GCCGCUaucaa..............auugaaUUACCACGCCAAAGACGUCUACAUCGUUG |
| NC_009525/3198691-3198756 | .CCACGUCCGACGGC.AACGAC.GCCGCUaucaa..............auugaaUUACCACGCCAAAGACGUCUACAUCGUUG |
| NC_018143/3186709-3186774 | .CCACGUCCGACGGC.AACGAC.GCCGCUaucaa..............auugaaUUACCACGCCAAAGACGUCUACAUCGUUG |
| NZ_CP009480/3179895-3179960 | .CCACGUCCGACGGC.AACGAC.GCCGCUaucaa..............auugaaUUACCACGCCAAAGACGUCUACAUCGUUG |
| NZ_CP007027/3186087-3186152 | .CCACGUCCGACGGC.AACGAC.GCCGCUaucaa..............auugaaUUACCACGCCAAAGACGUCUACAUCGUUG |
| NZ_CP002871/3171303-3171368 | .CCACGUCCGACGGC.AACGAC.GCCGCUaucaa..............auugaaUUACCACGCCAAAGACGUCUACAUCGUUG |
| NZ_CP007803/3161696-3161761 | .CCACGUCCGACGGC.AACGAC.GCCGCUaucaa..............auugaaUUACCACGCCAAAGACGUCUACAUCGUUG |
| NC_012943/1227063-1227128 | .CCACGUCCGACGGC.AACGAC.GCCGCUaucaa..............auugaaUUACCACGCCAAAGACGUCUACAUCGUUG |
| NC_016768/1227061-1227126 | .CCACGUCCGACGGC.AACGAC.GCCGCUaucaa..............auugaaUUACCACGCCAAAGACGUCUACAUCGUUG |
| NC_018078/1227175-1227240 | .CCACGUCCGACGGC.AACGAC.GCCGCUaucaa..............auugaaUUACCACGCCAAAGACGUCUACAUCGUUG |
| NC_021054/3186391-3186456 | .CCACGUCCGACGGC.AACGAC.GCCGCUaucaa..............auugaaUUACCACGCCAAAGACGUCUACAUCGUUG |
| NC_020559/3175270-3175335 | .CCACGUCCGACGGC.AACGAC.GCCGCUaucaa..............auugaaUUACCACGCCAAAGACGUCUACAUCGUUG |
| NC_022350/3182204-3182269 | .CCACGUCCGACGGC.AACGAC.GCCGCUaucaa..............auugaaUUACCACGCCAAAGACGUCUACAUCGUUG |
| NZ_AP014573/3192537-3192602 | .CCACGUCCGACGGC.AACGAC.GCCGCUaucaa..............auugaaUUACCACGCCAAAGACGUCUACAUCGUUG |
| NZ_CP010339/3184584-3184649 | .CCACGUCCGACGGC.AACGAC.GCCGCUaucaa..............auugaaUUACCACGCCAAAGACGUCUACAUCGUUG |
| NZ_CP010337/3193423-3193488 | .CCACGUCCGACGGC.AACGAC.GCCGCUaucaa..............auugaaUUACCACGCCAAAGACGUCUACAUCGUUG |
| NZ_CP010340/3205322-3205387 | .CCACGUCCGACGGC.AACGAC.GCCGCUaucaa..............auugaaUUACCACGCCAAAGACGUCUACAUCGUUG |
| NZ_CP010338/3179523-3179588 | .CCACGUCCGACGGC.AACGAC.GCCGCUaucaa..............auugaaUUACCACGCCAAAGACGUCUACAUCGUUG |
| NZ_CP009426/3156975-3157040 | .CCACGUCCGACGGC.AACGAC.GCCGCUaucaa..............auugaaUUACCACGCCAAAGACGUCUACAUCGUUG |
| NZ_CP009427/3194915-3194980 | .CCACGUCCGACGGC.AACGAC.GCCGCUaucaa..............auugaaUUACCACGCCAAAGACGUCUACAUCGUUG |
| NZ_CP010330/3195709-3195774 | .CCACGUCCGACGGC.AACGAC.GCCGCUaucaa..............auugaaUUACCACGCCAAAGACGUCUACAUCGUUG |
| NZ_CP007809/3171105-3171170 | .CCACGUCCGACGGC.AACGAC.GCCGCUaucaa..............auugaaUUACCACGCCAAAGACGUCUACAUCGUUG |
| NZ_CP012506/3156765-3156830 | .CCACGUCCGACGGC.AACGAC.GCCGCUaucaa..............auugaaUUACCACGCCAAAGACGUCUACAUCGUUG |
| NZ_CP009100/3186717-3186782 | .CCACGUCCGACGGC.AACGAC.GCCGCUaucaa..............auugaaUUACCACGCCAAAGACGUCUACAUCGUUG |
| NZ_CP009101/3186728-3186793 | .CCACGUCCGACGGC.AACGAC.GCCGCUaucaa..............auugaaUUACCACGCCAAAGACGUCUACAUCGUUG |
| NZ_CP012090/2955117-2955182 | .CCACGUCCGACGGC.AACGAC.GCCGCUaucaa..............auugaaUUACCACGCCAAAGACGUCUACAUCGUUG |
| NC_000962/3186699-3186764 | .CCACGUCCGACGGC.AACGAC.GCCGCUaucaa..............auugaaUUACCACGCCAAAGACGUCUACAUCGUUG |
| NC_015758/3501741-3501787 | .AAAAAGACGG--UC.ACCGAG.GA--GU.........................----AUCCGGAAAGGCCCGGUCCGGUAGC |
| NZ_CP003494/3183593-3183639 | .AAAAAGACGG--UC.ACCGAG.GA--GU.........................----AUCCGGAAAGGCCCGGUCCGGUAGC |
| NC_020245/3442987-3443033 | .AAAAAGACGG--UC.ACCGAG.GA--GU.........................----AUCCGGAAAGGCCCGGUCCGGUAGC |
| NC_016804/3453057-3453103 | .AAAAAGACGG--UC.ACCGAG.GA--GU.........................----AUCCGGAAAGGCCCGGUCCGGUAGC |
| NZ_AM412059/3465139-3465185 | .AAAAAGACGG--UC.ACCGAG.GA--GU.........................----AUCCGGAAAGGCCCGGUCCGGUAGC |
| NC_008769/3473692-3473738 | .AAAAAGACGG--UC.ACCGAG.GA--GU.........................----AUCCGGAAAGGCCCGGUCCGGUAGC |
| NC_012207/3466342-3466388 | .AAAAAGACGG--UC.ACCGAG.GA--GU.........................----AUCCGGAAAGGCCCGGUCCGGUAGC |
| NZ_CP014566/3466342-3466388 | .AAAAAGACGG--UC.ACCGAG.GA--GU.........................----AUCCGGAAAGGCCCGGUCCGGUAGC |
| NZ_CP008744/3452998-3453044 | .AAAAAGACGG--UC.ACCGAG.GA--GU.........................----AUCCGGAAAGGCCCGGUCCGGUAGC |
| NZ_CP009243/3466347-3466393 | .AAAAAGACGG--UC.ACCGAG.GA--GU.........................----AUCCGGAAAGGCCCGGUCCGGUAGC |
| NZ_CP012095/3480688-3480734 | .AAAAAGACGG--UC.ACCGAG.GA--GU.........................----AUCCGGAAAGGCCCGGUCCGGUAGC |
| NZ_CP009449/3428326-3428372 | .AAAAAGACGG--UC.ACCGAG.GA--GU.........................----AUCCGGAAAGGCCCGGUCCGGUAGC |
| NZ_CP013741/3466747-3466793 | .AAAAAGACGG--UC.ACCGAG.GA--GU.........................----AUCCGGAAAGGCCCGGUCCGGUAGC |
| NZ_CP015773/2382782-2382828 | .AAAAAGACGG--UC.ACCGAG.GA--GU.........................----AUCCGGAAAGGCCCGGUCCGGUAGC |
| NC_015848/3575680-3575726 | .AAAAAGACGG--UC.ACCGAG.GA--GU.........................----AUCCGGAAAGGCCCGGUCCGGUAGC |
| NZ_HG813240/3511601-3511647 | .AAAAAGACGG--UC.ACCGAG.GA--GU.........................----AUCCGGAAAGGCCCGGUCCGGUAGC |
| NC_020089/3517565-3517611 | .AAAAAGACGG--UC.ACCGAG.GA--GU.........................----AUCCGGAAAGGCCCGGUCCGGUAGC |
| NZ_CP002883/3506187-3506233 | .AAAAAGACGG--UC.ACCGAG.GA--GU.........................----AUCCGGAAAGGCCCGGUCCGGUAGC |
| NZ_CP002882/3500459-3500505 | .AAAAAGACGG--UC.ACCGAG.GA--GU.........................----AUCCGGAAAGGCCCGGUCCGGUAGC |
| NC_021251/3512644-3512690 | .AAAAAGACGG--UC.ACCGAG.GA--GU.........................----AUCCGGAAAGGCCCGGUCCGGUAGC |
| NC_017522/3506230-3506276 | .AAAAAGACGG--UC.ACCGAG.GA--GU.........................----AUCCGGAAAGGCCCGGUCCGGUAGC |
| NZ_CP002885/3511418-3511464 | .AAAAAGACGG--UC.ACCGAG.GA--GU.........................----AUCCGGAAAGGCCCGGUCCGGUAGC |
| NC_002755/3516130-3516176 | .AAAAAGACGG--UC.ACCGAG.GA--GU.........................----AUCCGGAAAGGCCCGGUCCGGUAGC |
| NC_017524/3517789-3517835 | .AAAAAGACGG--UC.ACCGAG.GA--GU.........................----AUCCGGAAAGGCCCGGUCCGGUAGC |
| NC_021740/3507069-3507115 | .AAAAAGACGG--UC.ACCGAG.GA--GU.........................----AUCCGGAAAGGCCCGGUCCGGUAGC |
| NC_021194/3506397-3506443 | .AAAAAGACGG--UC.ACCGAG.GA--GU.........................----AUCCGGAAAGGCCCGGUCCGGUAGC |
| NC_009565/3533768-3533814 | .AAAAAGACGG--UC.ACCGAG.GA--GU.........................----AUCCGGAAAGGCCCGGUCCGGUAGC |
| NC_009525/3532801-3532847 | .AAAAAGACGG--UC.ACCGAG.GA--GU.........................----AUCCGGAAAGGCCCGGUCCGGUAGC |
| NC_000962/3520645-3520691 | .AAAAAGACGG--UC.ACCGAG.GA--GU.........................----AUCCGGAAAGGCCCGGUCCGGUAGC |
| [illegible] | [illegible] |

| | |
|---|---|
| NC_018143/3520642-3520688 | .AAAAAGACGG--UC.ACCGAG.GA--GU.........................----AUCCGGAAAGGCCCGGUCCGGUAGC |
| NZ_CP009480/3513790-3513836 | .AAAAAGACGG--UC.ACCGAG.GA--GU.........................----AUCCGGAAAGGCCCGGUCCGGUAGC |
| NZ_CP007027/3520027-3520073 | .AAAAAGACGG--UC.ACCGAG.GA--GU.........................----AUCCGGAAAGGCCCGGUCCGGUAGC |
| NZ_CP002871/3510433-3510479 | .AAAAAGACGG--UC.ACCGAG.GA--GU.........................----AUCCGGAAAGGCCCGGUCCGGUAGC |
| NZ_CP007803/3498888-3498934 | .AAAAAGACGG--UC.ACCGAG.GA--GU.........................----AUCCGGAAAGGCCCGGUCCGGUAGC |
| NC_012943/3521468-3521514 | .AAAAAGACGG--UC.ACCGAG.GA--GU.........................----AUCCGGAAAGGCCCGGUCCGGUAGC |
| NC_016768/3518202-3518248 | .AAAAAGACGG--UC.ACCGAG.GA--GU.........................----AUCCGGAAAGGCCCGGUCCGGUAGC |
| NC_018078/3522581-3522627 | .AAAAAGACGG--UC.ACCGAG.GA--GU.........................----AUCCGGAAAGGCCCGGUCCGGUAGC |
| NC_021054/3520296-3520342 | .AAAAAGACGG--UC.ACCGAG.GA--GU.........................----AUCCGGAAAGGCCCGGUCCGGUAGC |
| NC_020559/3506597-3506643 | .AAAAAGACGG--UC.ACCGAG.GA--GU.........................----AUCCGGAAAGGCCCGGUCCGGUAGC |
| NC_022350/3514051-3514097 | .AAAAAGACGG--UC.ACCGAG.GA--GU.........................----AUCCGGAAAGGCCCGGUCCGGUAGC |
| NZ_AP014573/3528351-3528397 | .AAAAAGACGG--UC.ACCGAG.GA--GU.........................----AUCCGGAAAGGCCCGGUCCGGUAGC |
| NZ_CP010339/3520429-3520475 | .AAAAAGACGG--UC.ACCGAG.GA--GU.........................----AUCCGGAAAGGCCCGGUCCGGUAGC |
| NZ_CP010337/3529612-3529658 | .AAAAAGACGG--UC.ACCGAG.GA--GU.........................----AUCCGGAAAGGCCCGGUCCGGUAGC |
| NZ_CP010340/3538115-3538161 | .AAAAAGACGG--UC.ACCGAG.GA--GU.........................----AUCCGGAAAGGCCCGGUCCGGUAGC |
| NZ_CP010338/3513951-3513997 | .AAAAAGACGG--UC.ACCGAG.GA--GU.........................----AUCCGGAAAGGCCCGGUCCGGUAGC |
| NZ_CP009426/3490103-3490149 | .AAAAAGACGG--UC.ACCGAG.GA--GU.........................----AUCCGGAAAGGCCCGGUCCGGUAGC |
| NZ_CP009427/3526295-3526341 | .AAAAAGACGG--UC.ACCGAG.GA--GU.........................----AUCCGGAAAGGCCCGGUCCGGUAGC |
| NZ_CP010330/3529815-3529861 | .AAAAAGACGG--UC.ACCGAG.GA--GU.........................----AUCCGGAAAGGCCCGGUCCGGUAGC |
| NZ_CP007809/3508801-3508847 | .AAAAAGACGG--UC.ACCGAG.GA--GU.........................----AUCCGGAAAGGCCCGGUCCGGUAGC |
| NZ_CP012506/3490251-3490297 | .AAAAAGACGG--UC.ACCGAG.GA--GU.........................----AUCCGGAAAGGCCCGGUCCGGUAGC |
| NZ_CP009100/3520640-3520686 | .AAAAAGACGG--UC.ACCGAG.GA--GU.........................----AUCCGGAAAGGCCCGGUCCGGUAGC |
| NZ_CP009101/3520652-3520698 | .AAAAAGACGG--UC.ACCGAG.GA--GU.........................----AUCCGGAAAGGCCCGGUCCGGUAGC |
| NZ_CP012090/897334-897380 | .AAAAAGACGG--UC.ACCGAG.GA--GU.........................----AUCCGGAAAGGCCCGGUCCGGUAGC |
| NC_019966/3367677-3367738 | .AUUACUAAUGUCGG.GGACA-.CCGG.................acacugacuAUGGGGGGCGCAAUGAAGCGUCAGCGAAUU |
| NC_008596/5536087-5536152 | .cggugcaagGCUCG.UUGUU-.CGAGC.............augccgaacgucgCCCGCUGGGCAAUGGCGCCUGGAUCGAGC |
| NC_018289/5537699-5537764 | .cggugcaagGCUCG.UUGUU-.CGAGC.............augccgaacgucgCCCGCUGGGCAAUGGCGCCUGGAUCGAGC |
| NZ_CP009494/5536133-5536198 | .cggugcaagGCUCG.UUGUU-.CGAGC.............augccgaacgucgCCCGCUGGGCAAUGGCGCCUGGAUCGAGC |
| NZ_CP009495/5536187-5536252 | .cggugcaagGCUCG.UUGUU-.CGAGC.............augccgaacgucgCCCGCUGGGCAAUGGCGCCUGGAUCGAGC |
| NZ_CP009496/5536168-5536233 | .cggugcaagGCUCG.UUGUU-.CGAGC.............augccgaacgucgCCCGCUGGGCAAUGGCGCCUGGAUCGAGC |
| NZ_LN831039/5543050-5543115 | .cggugcaagGCUCG.UUGUU-.CGAGC.............augccgaacgucgCCCGCUGGGCAAUGGCGCCUGGAUCGAGC |
| NC_021282/3437670-3437736 | .AAACGCGUACGGAA.UGCAU-.AUUCUGAcg............uuugcgcacGUGGAGUGCGCAACGAUGCGUCGGAUUAUG |
| NZ_AP014547/4115495-4115542 | .--AUAAGACAGUUC.GUACAA.GGA..............................GGAAGAGCAAUGGCGCUCGUACUGGAA |
| NC_023036/1555181-1555233 | .ACGAUGCGAGGGAC.AAAGCA.GUCUC--.......................--UGGUUGCGCGAUGGUGUUCUGCGAAGGA |
| NZ_CP011022/1555180-1555232 | .ACGAUGCGAGGGAC.AAAGCA.GUCUC--.......................--UGGUUGCGCGAUGGUGUUCUGCGAAGGA |
| NZ_CP014950/3287342-3287395 | .AUGAGCUAGAGCAC.AUUGAC.GUGCU--.......................-UGCCGCGAGCGAAGGAGCGACGCCCCAUG |
| NZ_CP014951/3721005-3721058 | .AUGAGCUAGAGCAC.AUUGAC.GUGCU--.......................-UGCCGCGAGCGAAGGAGCGACGCCCCAUG |
| NZ_CP014952/3364829-3364882 | .AUGAGCUAGAGCAC.AUUGAC.GUGCU--.......................-UGCCGCGAGCGAAGGAGCGACGCCCCAUG |
| NZ_CP014953/3586037-3586090 | .AUGAGCUAGAGCAC.AUUGAC.GUGCU--.......................-UGCCGCGAGCGAAGGAGCGACGCCCCAUG |
| NZ_CP014954/3686526-3686579 | .AUGAGCUAGAGCAC.AUUGAC.GUGCU--.......................-UGCCGCGAGCGAAGGAGCGACGCCCCAUG |
| NZ_CP014957/3490346-3490399 | .AUGAGCUAGAGCAC.AUUGAC.GUGCU--.......................-UGCCGCGAGCGAAGGAGCGACGCCCCAUG |
| NZ_CP014958/3693121-3693174 | .AUGAGCUAGAGCAC.AUUGAC.GUGCU--.......................-UGCCGCGAGCGAAGGAGCGACGCCCCAUG |
| NZ_CP014959/3453708-3453761 | .AUGAGCUAGAGCAC.AUUGAC.GUGCU--.......................-UGCCGCGAGCGAAGGAGCGACGCCCCAUG |
| NZ_CP014961/3791427-3791480 | .AUGAGCUAGAGCAC.AUUGAC.GUGCU--.......................-UGCCGCGAGCGAAGGAGCGACGCCCCAUG |
| NZ_CP009615/1450815-1450868 | .AUGAGCUAGAGCAC.AUUGAC.GUGCU--.......................-UGCCGCGAGCGAAGGAGCGACGCCCCAUG |
| NZ_CP009616/1450964-1451017 | .AUGAGCUAGAGCAC.AUUGAC.GUGCU--.......................-UGCCGCGAGCGAAGGAGCGACGCCCCAUG |
| NC_021282/3403861-3403914 | .AUGAGCUAGAGCAC.AUUGAC.GUGCU--.......................-UGCCGCGAGCGAAGGAGCGACGCCCCAUG |
| NZ_AP014547/3435184-3435237 | .AUGAGCUAGAGCAC.AUUGAC.GUGCU--.......................-UGCCGCGAGCGAAGGAGCGACGCCCCAUG |
| NZ_CP009447/1435642-1435695 | .AUGAGCUAGAGCAC.AUUGAC.GUGCU--.......................-UGCCGCGAGCGAAGGAGCGACGCCCCAUG |
| NC_018150/1450987-1451040 | .AUGAGCUAGAGCAC.AUUGAC.GUGCU--.......................-UGCCGCGAGCGAAGGAGCGACGCCCCAUG |
| NZ_CP014952/4425526-4425574 | .AUAGACC....GCG.GCCAGA.UGC...........................UCAGUCGGCGCAAUGGUGCGCAGAAAGAUG |
| NZ_CP014955/4582395-4582443 | .AUAGACC....GCG.GCCAGA.UGC...........................UCAGUCGGCGCAAUGGUGCGCAGAAAGAUG |
| NZ_CP014956/4697458-4697506 | .AUAGACC....GCG.GCCAGA.UGC...........................UCAGUCGGCGCAAUGGUGCGCAGAAAGAUG |
| NZ_CP014960/4392617-4392665 | .AUAGACC....GCG.GCCAGA.UGC...........................UCAGUCGGCGCAAUGGUGCGCAGAAAGAUG |
| NZ_CP013049/4583427-4583475 | .AUAGACC....GCG.GCCAGA.UGC...........................UCAGUCGGCGCAAUGGUGCGCAGAAAGAUG |
| NZ_CP009407/4559500-4559548 | .AUAGACC....GCG.GCCAGA.UGC...........................UCAGUCGGCGCAAUGGUGCGCAGAAAGAUG |
| NZ_CP009408/4572347-4572395 | .AUAGACC....GCG.GCCAGA.UGC...........................UCAGUCGGCGCAAUGGUGCGCAGAAAGAUG |
| NZ_CP009613/4557377-4557425 | .AUAGACC....GCG.GCCAGA.UGC...........................UCAGUCGGCGCAAUGGUGCGCAGAAAGAUG |
| NZ_CP012044/4843659-4843707 | .AUAGACC....GCG.GCCAGA.UGC...........................UCAGUCGGCGCAAUGGUGCGCAGAAAGAUG |
| NC_010397/4575201-4575249 | .AUAGACC....GCG.GCCAGA.UGC...........................UCAGUCGGCGCAAUGGUGCGCAGAAAGAUG |
| NZ_CP014951/4374135-4374182 | .--AUAAGACAGUUC.GUACAA.GGA..............................GGAAGAGCAAUGGCGCUCGUACUGGAA |
| NZ_CP014953/4197267-4197314 | .--AUAAGACAGUUC.GUACAA.GGA..............................GGAAGAGCAAUGGCGCUCGUACUGGAA |
| NZ_CP014954/4297756-4297803 | .--AUAAGACAGUUC.GUACAA.GGA..............................GGAAGAGCAAUGGCGCUCGUACUGGAA |
| NZ_CP014958/4304351-4304398 | .--AUAAGACAGUUC.GUACAA.GGA..............................GGAAGAGCAAUGGCGCUCGUACUGGAA |
| NZ_CP009615/851498-851545 | .--AUAAGACAGUUC.GUACAA.GGA..............................GGAAGAGCAAUGGCGCUCGUACUGGAA |
| NZ_CP009616/851598-851645 | .--AUAAGACAGUUC.GUACAA.GGA..............................GGAAGAGCAAUGGCGCUCGUACUGGAA |
| NC_021282/4061154-4061201 | .--AUAAGACAGUUC.GUACAA.GGA..............................GGAAGAGCAAUGGCGCUCGUACUGGAA |
| NZ_CP009447/837674-837721 | .--AUAAGACAGUUC.GUACAA.GGA..............................GGAAGAGCAAUGGCGCUCGUACUGGAA |
| NC_018150/851623-851670 | .--AUAAGACAGUUC.GUACAA.GGA..............................GGAAGAGCAAUGGCGCUCGUACUGGAA |
| NZ_HG813240/2789446-2789495 | .CAAAACCGGCUGGC.AACCA-.GCCGUU.........................----UACCGCAACGAUGCGUGUGCCUGAC |
| NZ_CP002883/2783616-2783665 | .CAAAACCGGCUGGC.AACCA-.GCCGUU.........................----UACCGCAACGAUGCGUGUGCCUGAC |
| NZ_CP002882/2776221-2776270 | .CAAAACCGGCUGGC.AACCA-.GCCGUU.........................----UACCGCAACGAUGCGUGUGCCUGAC |
| NC_021251/2788548-2788597 | .CAAAACCGGCUGGC.AACCA-.GCCGUU.........................----UACCGCAACGAUGCGUGUGCCUGAC |

```
NC_021251/2788548-2788597     .CAAAACCGGCUGGC.AACCA-.GCCGUU.........................----UACCGCAACGAUGCGUGUGCCUGAC
NC_017522/2782217-2782266     .CAAAACCGGCUGGC.AACCA-.GCCGUU.........................----UACCGCAACGAUGCGUGUGCCUGAC
NZ_CP002885/2786794-2786843   .CAAAACCGGCUGGC.AACCA-.GCCGUU.........................----UACCGCAACGAUGCGUGUGCCUGAC
NC_009565/2807325-2807374     .CAAAACCGGCUGGC.AACCA-.GCCGUU.........................----UACCGCAACGAUGCGUGUGCCUGAC
NZ_CP002871/2785983-2786032   .CAAAACCGGCUGGC.AACCA-.GCCGUU.........................----UACCGCAACGAUGCGUGUGCCUGAC
NZ_CP007803/2777337-2777386   .CAAAACCGGCUGGC.AACCA-.GCCGUU.........................----UACCGCAACGAUGCGUGUGCCUGAC
NC_021054/2793756-2793805     .CAAAACCGGCUGGC.AACCA-.GCCGUU.........................----UACCGCAACGAUGCGUGUGCCUGAC
NZ_CP009426/2776800-2776849   .CAAAACCGGCUGGC.AACCA-.GCCGUU.........................----UACCGCAACGAUGCGUGUGCCUGAC
NZ_CP007809/2786461-2786510   .CAAAACCGGCUGGC.AACCA-.GCCGUU.........................----UACCGCAACGAUGCGUGUGCCUGAC
NZ_CP012506/2770232-2770281   .CAAAACCGGCUGGC.AACCA-.GCCGUU.........................----UACCGCAACGAUGCGUGUGCCUGAC
NZ_CP009100/2794045-2794094   .CAAAACCGGCUGGC.AACCA-.GCCGUU.........................----UACCGCAACGAUGCGUGUGCCUGAC
NZ_CP009101/2794056-2794105   .CAAAACCGGCUGGC.AACCA-.GCCGUU.........................----UACCGCAACGAUGCGUGUGCCUGAC
NZ_CP012090/2568439-2568488   .CAAAACCGGCUGGC.AACCA-.GCCGUU.........................----UACCGCAACGAUGCGUGUGCCUGAC
NC_015758/2780806-2780855     .CAAAACCGGCUGGC.AACCA-.GCCGUU.........................----UACCGCAACGAUGCGUGUGCCUGAC
NZ_CP003494/2467080-2467129   .CAAAACCGGCUGGC.AACCA-.GCCGUU.........................----UACCGCAACGAUGCGUGUGCCUGAC
NC_020245/2727463-2727512     .CAAAACCGGCUGGC.AACCA-.GCCGUU.........................----UACCGCAACGAUGCGUGUGCCUGAC
NC_016804/2737183-2737232     .CAAAACCGGCUGGC.AACCA-.GCCGUU.........................----UACCGCAACGAUGCGUGUGCCUGAC
NZ_AM412059/2750312-2750361   .CAAAACCGGCUGGC.AACCA-.GCCGUU.........................----UACCGCAACGAUGCGUGUGCCUGAC
NC_008769/2757818-2757867     .CAAAACCGGCUGGC.AACCA-.GCCGUU.........................----UACCGCAACGAUGCGUGUGCCUGAC
NC_012207/2750589-2750638     .CAAAACCGGCUGGC.AACCA-.GCCGUU.........................----UACCGCAACGAUGCGUGUGCCUGAC
NZ_CP014566/2750588-2750637   .CAAAACCGGCUGGC.AACCA-.GCCGUU.........................----UACCGCAACGAUGCGUGUGCCUGAC
NZ_CP008744/2737952-2738001   .CAAAACCGGCUGGC.AACCA-.GCCGUU.........................----UACCGCAACGAUGCGUGUGCCUGAC
NZ_CP009243/2750593-2750642   .CAAAACCGGCUGGC.AACCA-.GCCGUU.........................----UACCGCAACGAUGCGUGUGCCUGAC
NZ_CP012095/2765911-2765960   .CAAAACCGGCUGGC.AACCA-.GCCGUU.........................----UACCGCAACGAUGCGUGUGCCUGAC
NZ_CP009449/2715394-2715443   .CAAAACCGGCUGGC.AACCA-.GCCGUU.........................----UACCGCAACGAUGCGUGUGCCUGAC
NZ_CP013741/2750888-2750937   .CAAAACCGGCUGGC.AACCA-.GCCGUU.........................----UACCGCAACGAUGCGUGUGCCUGAC
NZ_CP015773/1667473-1667522   .CAAAACCGGCUGGC.AACCA-.GCCGUU.........................----UACCGCAACGAUGCGUGUGCCUGAC
NC_015848/2855879-2855928     .CAAAACCGGCUGGC.AACCA-.GCCGUU.........................----UACCGCAACGAUGCGUGUGCCUGAC
NC_020089/2794623-2794672     .CAAAACCGGCUGGC.AACCA-.GCCGUU.........................----UACCGCAACGAUGCGUGUGCCUGAC
NC_002755/2789497-2789546     .CAAAACCGGCUGGC.AACCA-.GCCGUU.........................----UACCGCAACGAUGCGUGUGCCUGAC
NC_017524/2788961-2789010     .CAAAACCGGCUGGC.AACCA-.GCCGUU.........................----UACCGCAACGAUGCGUGUGCCUGAC
NC_021740/2781801-2781850     .CAAAACCGGCUGGC.AACCA-.GCCGUU.........................----UACCGCAACGAUGCGUGUGCCUGAC
NC_021194/2781313-2781362     .CAAAACCGGCUGGC.AACCA-.GCCGUU.........................----UACCGCAACGAUGCGUGUGCCUGAC
NC_009525/2805977-2806026     .CAAAACCGGCUGGC.AACCA-.GCCGUU.........................----UACCGCAACGAUGCGUGUGCCUGAC
NC_000962/2794009-2794058     .CAAAACCGGCUGGC.AACCA-.GCCGUU.........................----UACCGCAACGAUGCGUGUGCCUGAC
NC_018143/2794019-2794068     .CAAAACCGGCUGGC.AACCA-.GCCGUU.........................----UACCGCAACGAUGCGUGUGCCUGAC
NZ_CP009480/2788437-2788486   .CAAAACCGGCUGGC.AACCA-.GCCGUU.........................----UACCGCAACGAUGCGUGUGCCUGAC
NZ_CP007027/2793397-2793446   .CAAAACCGGCUGGC.AACCA-.GCCGUU.........................----UACCGCAACGAUGCGUGUGCCUGAC
NC_012943/1620982-1621031     .CAAAACCGGCUGGC.AACCA-.GCCGUU.........................----UACCGCAACGAUGCGUGUGCCUGAC
NC_016768/1620926-1620975     .CAAAACCGGCUGGC.AACCA-.GCCGUU.........................----UACCGCAACGAUGCGUGUGCCUGAC
NC_018078/1621097-1621146     .CAAAACCGGCUGGC.AACCA-.GCCGUU.........................----UACCGCAACGAUGCGUGUGCCUGAC
NC_020559/2781596-2781645     .CAAAACCGGCUGGC.AACCA-.GCCGUU.........................----UACCGCAACGAUGCGUGUGCCUGAC
NC_022350/2788615-2788664     .CAAAACCGGCUGGC.AACCA-.GCCGUU.........................----UACCGCAACGAUGCGUGUGCCUGAC
NZ_AP014573/2798411-2798460   .CAAAACCGGCUGGC.AACCA-.GCCGUU.........................----UACCGCAACGAUGCGUGUGCCUGAC
NZ_CP010339/2793212-2793261   .CAAAACCGGCUGGC.AACCA-.GCCGUU.........................----UACCGCAACGAUGCGUGUGCCUGAC
NZ_CP010337/2801876-2801925   .CAAAACCGGCUGGC.AACCA-.GCCGUU.........................----UACCGCAACGAUGCGUGUGCCUGAC
NZ_CP010340/2812494-2812543   .CAAAACCGGCUGGC.AACCA-.GCCGUU.........................----UACCGCAACGAUGCGUGUGCCUGAC
NZ_CP010338/2793070-2793119   .CAAAACCGGCUGGC.AACCA-.GCCGUU.........................----UACCGCAACGAUGCGUGUGCCUGAC
NZ_CP009427/2805466-2805515   .CAAAACCGGCUGGC.AACCA-.GCCGUU.........................----UACCGCAACGAUGCGUGUGCCUGAC
NZ_CP010330/2803025-2803074   .CAAAACCGGCUGGC.AACCA-.GCCGUU.........................----UACCGCAACGAUGCGUGUGCCUGAC
RF                            .AaaAaaaauagcAC.AAaGgc.GUgcUuu.......................uauuaaAaAGCAAuGaAGCUcauaaaaAaa
SS_cons                       .::::::::::<<<<.______.>>>>---.......................--------<<<______>>>::::::::::
```

*Table S4: GO analyses and literature curation of differentially expressed genes in M.smegmatis under ANTAR~P conditions*

GO enrichment analyses is carried out using GOA (Gene Ontology Analysis Consortium; ftp://ftp.ebi.ac.uk/pub/databases/GO/goa/proteomes/25827.M_smegmatis.goa) and BLAST2GO (PMID: 18483572). GO IDs and GO descriptions are shown with uncorrected and Bonferroni-corrected p-values. GO terms are classified in biological processes (BP), molecular function (MF) and cellular component (CC). The genes are categorized based on GO and literature based curation.

| **Up-regulated genes** | | | | | | | | | |
|---|---|---|---|---|---|---|---|---|---|
| **Reductive Metabolism** | | | | | | | | | |
| | | | | | **GOA based Gene Annotation** | | | **BLAST2GO based GO annotation** | |
| **Locus tag (gene name)** | **Gene Annotation** | **Location** | **Strand** | **Length** | **GO category: GO ID \| GO terms** | **Uncorrected p-value** | **Bonferroni Corrected p-value** | **GO category: GO ID \| GO terms** | **Literature based Annotation** |
| MSMEG_4511 | linear gramicidin synthetase subunit B | 4589756-4595215 | - | 5460 | MF: GO:0048037 \| cofactor binding<br>MF: GO:0043168 \| anion binding<br>MF: GO:0036094 \| small molecule binding<br>MF: GO:0043167 \| ion binding<br>MF: GO:0003824 \| catalytic activity<br>MF: GO:0005488 \| binding | 0.499078338762607<br>0.839250716415205<br>0.895685477437695<br>0.950043978332138<br>0.977353128529576<br>0.989284577095953 | 1.00<br>1.00<br>1.00<br>1.00<br>1.00<br>1.00 | MF: GO:0016853 \| isomerase activity<br>MF: GO:0016874 \| ligase activity<br>MF: GO:0043167 \| ion binding<br>MF: GO:0003824 \| catalytic activity<br>MF: GO:0031177 \| phosphopantetheine binding | 1. gramicidin-membrane peptide for transport of monovalent ions across lipid bilayer, phosphopantetheien binding;<br>2. Msmeg is sensitive to iron depletion and the effect increases with fxbA<br>3. MSMEG_4511 also called Mycobactin synthase E (mbtE) essential for mycobactin biosynthesis pathway |
| MSMEG_1680 | conserved hypothetical protein, interpro based domain: Nucleophile aminohydrolases, N-terminal (IPR029055) | 1773456-1774130 | - | 675 | NA | NA | NA | NA | DUF1028 domain protein |
| MSMEG_1677 (aspA) | aspartate ammonia-lyase | 1770884-1772305 | - | 1422 | BP: GO:0006091 \| generation of precursor metabolites and energy<br>BP: GO:0055114 \| oxidation-reduction process<br>BP: GO:0044281 \| small molecule metabolic process<br>BP: GO:0044237 \| cellular metabolic process<br>BP: GO:0071704 \| organic substance metabolic process<br>BP: GO:0044238 \| primary metabolic process<br>BP: GO:0009987 \| cellular process<br>BP: GO:0008152 \| metabolic process<br>MF: GO:0003824 \| catalytic activity | 0.134282946379772<br>0.773615546714231<br>0.86265269554733<br>0.954670564139802<br>0.969155736563623<br>0.976923093638226<br>0.979738654330216<br>0.990623690614229<br>0.977353128529576 | 1.00<br>1.00<br>1.00<br>1.00<br>1.00<br>1.00<br>1.00<br>1.00<br>1.00 | BP: GO:0006091 \| generation of precursor metabolites and energy<br>BP: GO:0044281 \| small molecule metabolic process<br>BP: GO:0006099 \| tricarboxylic acid cycle<br>MF: GO:0016829 \| lyase activity<br>MF: GO:0003824 \| catalytic activity | aspartate, alanine metabolism, nitrogen cycle, TCA cycle |
| MSMEG_1679 | AmiB; Peptidase M20 domain-containing protein 2 | 1772302-1773459 | - | 1158 | MF: GO:0016787 \| hydrolase activity | 0.548264938 | 1.00 | MF: GO:0016787 \| hydrolase activity<br>MF: GO:0003674 \| molecular function | yjeD, MurNAC alanine amidase |
| MSMEG_0113 | taurine transport system permease protein TauC; ABC-type nitrate/sulfonate/bicarbonate transport system, permease component (Orthology based) | 136950-137810 | + | 861 | BP: GO:0055085 \| transmembrane transport<br>BP: GO:0006810 \| transport<br>BP: GO:0051234 \| establishment of localization<br>BP: GO:0051179 \| localization<br>CC: GO:0016021 \| integral component of membrane<br>CC: GO:0031224 \| intrinsic component of membrane<br>CC: GO:0016020 \| membrane<br>CC: GO:0044425 \| membrane part | 0.492252741577174<br>0.610940825770566<br>0.610940825770566<br>0.562362624294413<br>0.0279921560341479<br>0.0279921560341479<br>0.0421000042900072<br>0.0287272290997126 | 1.00<br>1.00<br>1.00<br>1.00<br>0.13996078017074<br>0.13996078017074<br>0.210500021450036<br>0.143636145498563 | BP: GO:0055085 \| transmembrane transport<br>CC: GO:0005886 \| plasma membrane<br>CC: GO:0016020 \| membrane | nitrogen/carbon utilization, nitrogen cycle/sulfur cycle |

| | | | | | | | | | |
|---|---|---|---|---|---|---|---|---|---|
| MSMEG_5575 | repressor, marR type possibly DeoR; sugar binding | 5663425-5664351 | - | 927 | BP: GO:0006355 \| regulation of transcription, DNA-templated<br>BP: GO:1903506 \| regulation of nucleic acid-templated transcription<br>BP: GO:2001141 \| regulation of RNA biosynthetic process<br>BP: GO:0051252 \| regulation of RNA metabolic process<br>BP: GO:0019219 \| regulation of nucleobase-containing compound metabolic process<br>BP: GO:0009889 \| regulation of biosynthetic process<br>BP: GO:0010556 \| regulation of macromolecule biosynthetic process<br>BP: GO:0031326 \| regulation of cellular biosynthetic process<br>BP: GO:2000112 \| regulation of cellular macromolecule biosynthetic process<br>BP: GO:0010468 \| regulation of gene expression<br>BP: GO:0051171 \| regulation of nitrogen compound metabolic process<br>BP: GO:0080090 \| regulation of primary metabolic process<br>BP: GO:0060255 \| regulation of macromolecule metabolic process<br>BP: GO:0031323 \| regulation of cellular metabolic process<br>BP: GO:0019222 \| regulation of metabolic process<br>BP: GO:0050794 \| regulation of cellular process<br>BP: GO:0050789 \| regulation of biological process<br>BP: GO:0065007 \| biological regulation<br>MF: GO:0003676 \| nucleic acid binding<br>MF: GO:0005488 \| binding<br>MF: GO:0097159 \| organic cyclic compound binding<br>MF: GO:1901363 \| heterocyclic compound binding | 0.671382820793333<br>0.671382820793333<br>0.671382820793333<br>0.672865283740888<br>0.675814880235901<br>0.68454140923867<br>0.68454140923867<br>0.68454140923867<br>0.68454140923867<br>0.685978058920411<br>0.690257659202602<br>0.690257659202602<br>0.691674096952492<br>0.693085497428075<br>0.695893212658836<br>0.798541663428491<br>0.803492192042164<br>0.830219934607928<br>0.960169166760883<br>0.989284577095953<br>0.994381822820708<br>0.994381822820708 | 1.00<br>1.00<br>1.00<br>1.00 | BP: GO:0034641 \| cellular nitrogen compound metabolic process<br>BP: GO:0009058 \| biosynthetic process<br>BP: GO:0006355 \| regulation of transcription, DNA-templated<br>MF: GO:0003677 \| DNA binding<br>MF: GO:0003700 \| DNA-binding transcription factor activity<br>MF: GO:0030246 \| carbohydrate binding | 1. repressor for dNTP/NTP breakdown genes, cellular nitrogen compound metabolic process;<br>2. Downregulated when vapC toxin is overexpressed and is annotated as smoR (repressor for sugar alcohol). It is part of a smo operon and is the first gene in the operon regulating downstream genes |

### Down-regulated genes

#### Redox/ Stress

| | | | | | GOA based Gene Annotation | | | BLAST2GO based GO annotation | |
|---|---|---|---|---|---|---|---|---|---|
| **Locus tag (gene name)** | **Gene Annotation** | **Location** | **Strand** | **Length** | **GO category: GO ID \| GO terms** | **Uncorrected p-value** | **Bonferroni Corrected p-value** | **GO category: GO ID \| GO terms** | **Literature based Annotation** |
| MSMEG_1794 | dehydrogenase/LLM class F420 dependent oxidoreducatse | 1872888-1873895 | + | 1008 | BP: GO:0055114 \| oxidation-reduction process<br>BP: GO:0008152 \| metabolic process<br>MF: GO:0016491 \| oxidoreductase activity<br>MF: GO:0003824 \| catalytic activity | 0.773615546714231<br>0.990623690614229<br>0.875300204731563<br>0.977353128529576 | 1.00<br>1.00<br>1.00<br>1.00 | BP: GO:0055114 \| oxidation-reduction process<br>MF: GO:0016491 \| oxidoreductase activity<br>MF: GO:0016705 \| oxidoreductase activity, acting on paired donors, with incorporation or reduction of molecular oxygen | 1. oxidoreductase activity acting on CHNH group of donors, glucose6P dehydrogenase, coA F420, NO2 stress/redox<br>2. Downregulated in sigF mutant implicated in response to oxidative stress |
| MSMEG_1077 | conserved hypothetical protein; Deazaflavin-dependent nitroreductase family protein | 1141706-1142155 | - | 450 | BP: GO:0055114 \| oxidation-reduction process<br>BP: GO:0008152 \| metabolic process<br>MF: GO:0048037 \| cofactor binding<br>MF: GO:0016491 \| oxidoreductase activity<br>MF: GO:0003824 \| catalytic activity<br>MF: GO:0005488 \| binding | 0.773615546714231<br>0.990623690614229<br>0.499078338762607<br>0.875300204731563<br>0.977353128529576<br>0.989284577095953 | 1.00<br>1.00<br>1.00<br>1.00<br>1.00<br>1.00 | BP: GO:0055114 \| oxidation-reduction process<br>MF: GO:0016491 \| oxidoreductase activity<br>MF: GO:0048037 \| cofactor binding | 1. uses nitroimidazole substrates, releases RNS, (breaks down several drugs),<br>2. oxidoreductase |
| MSMEG_6212 | hemerythrin HHE cation binding domain subfamily protein, putative | 6276681-6277265 | + | 585 | NA | NA | NA | NA | 1. Can bind Fe, Cd, other metals, two examples involved in nitric oxide response.<br>2. *S. aureus* protein repairs oxxidative/nitrosative damage to FE-S cluster proteins- probably involved in oxygen binding /oxygen storage<br>3. Downregulated in sigF mutant implicated in response to oxidative stress |
| MSMEG_6213 | Manganese containing catalase; interpro based domain: IPR009078 Ferritin-like_SF IPR007760 Mn_catalase IPR039377 Mn_catalase_dom | 6277327-6278220 | - | 894 | NA | NA | NA | MF: GO:0016491 \| oxidoreductase activity | 1. metal binding, oxygen stress/ superoxide stress, peroxidase<br>2. Downregulated in sigF mutant implicated in response to oxidative stress |
| MSMEG_5189 | oxidoreductase; NAD-dependent epimerase/dehydratase (orthology based) | 5288282-5289418 | - | 1137 | NA | NA | NA | MF: GO:0016829 \| lyase activity | 1. NAD(P) cofactor, dTDP-glucose 4,6 dehydratase<br>2. Downregulated in sigF mutant implicated in response to oxidative stress |

| | | | | | | | | | |
|---|---|---|---|---|---|---|---|---|---|
| MSMEG_6768 | halogenase; FAD/ NAD binding (IPR036188) | 6816577-6818328 | - | 1752 | NA | NA | NA | BP: GO:0006091 \| generation of precursor metabolites and energy<br>MF: GO:0016491 \| oxidoreductase activity | 1. works with a cognate flavin reductase to reduce FADH2 while oxidizing halide, generation of precursor metabolites and energy, electron transfering flavoprotein dehydrogenase<br>2. Downregulated in sigF mutant implicated in response to oxidative stress |
| MSMEG_0686 | oxidoreductase | 769773-770294 | - | 522 | BP: GO:0006091 \| generation of precursor metabolites and energy<br>BP: GO:0055114 \| oxidation-reduction process<br>BP: GO:0044237 \| cellular metabolic process<br>BP: GO:0009987 \| cellular process<br>BP: GO:0008152 \| metabolic process<br>MF: GO:0048037 \| cofactor binding<br>MF: GO:0016491 \| oxidoreductase activity<br>MF: GO:0043167 \| ion binding<br>MF: GO:0003824 \| catalytic activity<br>MF: GO:0005488 \| binding | 0.134282946379772<br>0.773615546714231<br>0.954670564139802<br>0.979738654330216<br>0.990623690614229<br>0.499078338762607<br>0.875300204731563<br>0.950043978332138<br>0.977353128529576<br>0.989284577095953 | 1.00<br>1.00<br>1.00<br>1.00<br>1.00<br>1.00<br>1.00<br>1.00<br>1.00<br>1.00 | BP: GO:0006091 \| generation of precursor metabolites and energy<br>MF: GO:0016491 \| oxidoreductase activity<br>MF: GO:0043167 \| ion binding<br>MF: GO:0009055 \| electron transfer activity<br>MF: GO:0046872 \| metal ion binding<br>MF: GO:0051536 \| iron-sulfur cluster binding<br>MF: GO:0051537 \| 2 iron, 2 sulfur cluster binding | 1. Fe-S cluster binding, electron transport chain<br>2. Downregulated in sigF mutant implicated in response to oxidative stress |
| MSMEG_1132 (menJ) | FAD binding domain, putative | 1193567-1194796 | - | 1230 | BP: GO:0055114 \| oxidation-reduction process<br>BP: GO:0044281 \| small molecule metabolic process<br>BP: GO:0044249 \| cellular biosynthetic process<br>BP: GO:1901576 \| organic substance biosynthetic process<br>BP: GO:0009058 \| biosynthetic process<br>BP: GO:0044237 \| cellular metabolic process<br>BP: GO:0071704 \| organic substance metabolic process<br>BP: GO:0009987 \| cellular process<br>BP: GO:0008152 \| metabolic process<br>MF: GO:0048037 \| cofactor binding<br>MF: GO:0043168 \| anion binding<br>MF: GO:0016491 \| oxidoreductase activity<br>MF: GO:0036094 \| small molecule binding<br>MF: GO:0000166 \| nucleotide binding<br>MF: GO:1901265 \| nucleoside phosphate binding<br>MF: GO:0043167 \| ion binding<br>MF: GO:0003824 \| catalytic activity<br>MF: GO:0005488 \| binding<br>MF: GO:0097159 \| organic cyclic compound binding<br>MF: GO:1901363 \| heterocyclic compound binding | 0.773615546714231<br>0.86265269554733<br>0.895349081680126<br>0.893041017196166<br>0.929335307087835<br>0.954670564139802<br>0.969155736563623<br>0.979738654330216<br>0.990623690614229<br>0.499078338762607<br>0.839250716415205<br>0.875300204731563<br>0.895685477437695<br>0.939734266779146<br>0.939734266779146<br>0.950043978332138<br>0.977353128529576<br>0.989284577095953<br>0.994381822820708<br>0.994381822820708 | 1.00<br>1.00<br>1.00<br>1.00<br>1.00<br>1.00<br>1.00<br>1.00<br>1.00<br>1.00<br>1.00<br>1.00<br>1.00<br>1.00<br>1.00<br>1.00<br>1.00<br>1.00<br>1.00<br>1.00 | BP: GO:0055114 \| oxidation-reduction process<br>BP: GO:0051186 \| cofactor metabolic process<br>BP: GO:0044281 \| small molecule metabolic process<br>BP: GO:0009058 \| biosynthetic process<br>MF: GO:0016628 \| oxidoreductase activity, acting on the CH-CH group of donors, NAD or NADP as acceptor<br>MF: GO:0016491 \| oxidoreductase activity<br>MF: GO:0071949 \| FAD binding<br>MF: GO:0043167 \| ion binding | ion binding, FAD linked oxidoreductase,cofactor metabolic process |

### Membrane proteins/ ion transport

| | | | | | GOA based Gene Annotation | | | BLAST2GO based GO annotation | |
|---|---|---|---|---|---|---|---|---|---|
| Locus tag (gene name) | Gene Annotation | Location | Strand | Length | GO category: GO ID \| GO terms | Uncorrected p-value | Bonferroni Corrected p-value | GO category: GO ID \| GO terms | Literature based Annotation |
| MSMEG_3254 | RDD family protein, putative, transmembrane | 3336282-3336767 | - | 486 | NA | NA | NA | CC: GO:0005575 \| cellular_component | 1. transmembrane protein, one Helicobacter example is a Li/Na/K antiporter<br>2. Downregulated in sigF mutant implicated in response to oxidative stress |
| MSMEG_3255 | DoxX subfamily protein, putative | 3336777-3337652 | - | 876 | NA | NA | NA | NA | 1. transmembrane protein, phenotype in eukaryotes something to do with sodium depletion<br>2. Downregulated in sigF mutant implicated in response to oxidative stress |
| MSMEG_1131 | tryptophan-rich sensory protein | 1192990-1193475 | + | 486 | CC: GO:0016021 \| integral component of membrane<br>CC: GO:0031224 \| intrinsic component of membrane<br>CC: GO:0044425 \| membrane part<br>CC: GO:0016020 \| membrane | 0.0279921560341479<br>0.0279921560341479<br>0.0421000042900072<br>0.0287272290997126 | 0.13996078017074<br>0.13996078017074<br>0.210500021450036<br>0.143636145498563 | CC: GO:0016021 \| integral component of membrane<br>CC: GO:0005575 \| cellular_component | 1. integral membrane<br>2. Downregulated in sigF mutant implicated in response to oxidative stress |
| MSMEG_5496 | MscS Mechanosensitive ion channel | 5579234-5580490 | + | 1257 | BP: GO:0055085 \| transmembrane transport<br>BP: GO:0006810 \| transport<br>BP: GO:0051234 \| establishment of localization<br>BP: GO:0051179 \| localization<br>CC: GO:0016021 \| integral component of membrane<br>CC: GO:0031224 \| intrinsic component of membrane<br>CC: GO:0044425 \| membrane part<br>CC: GO:0016020 \| membrane | 0.492252741577174<br>0.610940825770566<br>0.610940825770566<br>0.562362624294413<br>0.0279921560341479<br>0.0279921560341479<br>0.0421000042900072<br>0.0287272290997126 | 1.00<br>1.00<br>1.00<br>1.00<br>0.13996078017074<br>0.13996078017074<br>0.210500021450036<br>0.143636145498563 | BP: GO:0055085 \| transmembrane transport<br>CC: GO:0016020 \| membrane<br>CC: GO:0005575 \| cellular_component | 1. interacts with ftsZ, osmolarity control, transmembrane transport<br>2. Downregulated in sigF mutant implicated in response to oxidative stress |
| MSMEG_6466 | aquaporin Z | 6532006-6532779 | + | 774 | BP: GO:0055085 \| transmembrane transport<br>BP: GO:0006810 \| transport<br>BP: GO:0051234 \| establishment of localization<br>BP: GO:0051179 \| localization<br>CC: GO:0016021 \| integral component of membrane<br>CC: GO:0031224 \| intrinsic component of membrane<br>CC: GO:0044425 \| membrane part<br>CC: GO:0016020 \| membrane | 0.492252741577174<br>0.610940825770566<br>0.610940825770566<br>0.562362624294413<br>0.0279921560341479<br>0.0279921560341479<br>0.0421000042900072<br>0.0287272290997126 | 1.00<br>1.00<br>1.00<br>1.00<br>0.13996078017074<br>0.13996078017074<br>0.210500021450036<br>0.143636145498563 | BP: GO:0055085 \| transmembrane transport<br>MF: GO:0022857 \| transmembrane transporter activity<br>MF: GO:0015267 \| channel activity<br>CC: GO:0016020 \| membrane<br>CC: GO:0005575 \| cellular_component | osmotic balance, transmembrane transport |

### Lipid catabolism

| | | | | | GOA based Gene Annotation | | | BLAST2GO based GO annotation | |
|---|---|---|---|---|---|---|---|---|---|
| Locus tag (gene name) | Gene Annotation | Location | Strand | Length | GO category: GO ID \| GO terms | Uncorrected p-value | Bonferroni Corrected p-value | GO category: GO ID \| GO terms | Literature based Annotation |
| MSMEG_2913 | hydrolase, alpha beta hydrolase fold protein | 2977263-2978126 | - | 864 | MF: GO:0016787 \| hydrolase activity<br>MF: GO:0003824 \| catalytic activity | 0.548264937793926<br>0.977353128529576 | 1.00<br>1.00 | MF: GO:0003824 \| catalytic activity<br>MF: GO:0003674 \| molecular_function | triglyceride lipase activity, haloalkane dehalogenase, carboxylesterase |
| MSMEG_0267 | esterase; Esterase/lipase lipF | 301478-302494 | + | 1017 | BP: GO:0006807 \| nitrogen compound metabolic process<br>BP: GO:0043170 \| macromolecule metabolic process<br>BP: GO:0071704 \| organic substance metabolic process<br>BP: GO:0044238 \| primary metabolic process<br>BP: GO:0008152 \| metabolic process<br>MF: GO:0016787 \| hydrolase activity<br>MF: GO:0003824 \| catalytic activity | 0.988878773902595<br>0.890726079425172<br>0.969155736563623<br>0.976923093638226<br>0.990623690614229<br>0.548264937793926<br>0.977353128529576 | 1.00<br>1.00<br>1.00<br>1.00<br>1.00<br>1.00<br>1.00 | BP: GO:0006508 \| proteolysis<br>MF: GO:0004252 \| serine-type endopeptidase activity<br>MF: GO:0008233 \| peptidase activity<br>MF: GO:0016787 \| hydrolase activity | is this ortholog of Rv3487c which has PLC activity on phosphatidylcholine,<br>1. serine type endopeptidase activity for proteolysis.<br>2. Downregulated in sigF mutant implicated in response to oxidative stress |

### Protein modification

| | | | | | GOA based Gene Annotation | | | BLAST2GO based GO annotation | |
|---|---|---|---|---|---|---|---|---|---|
| Locus tag (gene name) | Gene Annotation | Location | Strand | Length | GO category: GO ID \| GO terms | Uncorrected p-value | Bonferroni Corrected p-value | GO category: GO ID \| GO terms | Literature based Annotation |
| MSMEG_3289 | gp61 protein; alpha beta hydrolase fold protein | 3369713-3370798 | + | 1086 | NA | NA | NA | MF: GO:0016746 \| transferase activity, transferring acyl groups | 1. transferase activity, S-acetyl transferase or dihydrolipoyl lysine residue acetyl transferase<br>2. Downregulated in sigF mutant implicated in response to oxidative stress |

| | | | | | | | | | |
|---|---|---|---|---|---|---|---|---|---|
| MSMEG_1771 | methylase, putative | 1859306-1859974 | + | 669 | BP: GO:0008152 \| metabolic process<br>MF: GO:0016740 \| transferase activity<br>MF: GO:0003676 \| nucleic acid binding<br>MF: GO:0003824 \| catalytic activity<br>MF: GO:0005488 \| binding<br>MF: GO:0097159 \| organic cyclic compound binding<br>MF: GO:1901363 \| heterocyclic compound binding | 0.990623690614229<br>0.938690628502899<br>0.960169166760883<br>0.977353128529576<br>0.989284577095953<br>0.994381822820708<br>0.994381822820708 | 1.00<br>1.00<br>1.00<br>1.00<br>1.00<br>1.00<br>1.00 | BP: GO:0006464 \| cellular protein modification process<br>BP: GO:0032259 \| methylation<br>MF: GO:0008168 \| methyltransferase activity<br>MF: GO:0008276 \| protein methyltransferase activity<br>MF: GO:0003676 \| nucleic acid binding | 1. nucleic acid binding, methylation<br>2. Downregulated in sigF mutant implicated in response to oxidative stress |
| MSMEG_1097 | glycosyl transferase, group 2 family protein | 1160901-1161584 | + | 684 | MF: GO:0016740 \| transferase activity<br>MF: GO:0003824 \| catalytic activity | 0.938690628502899<br>0.977353128529576 | 1.00<br>1.00 | MF: GO:0003674 \| molecular_function | Downregulated in sigF mutant implicated in response to oxidative stress |
| MSMEG_3022 | transglycosylase associated protein | 3095235-3095540 | - | 306 | CC: GO:0016021 \| integral component of membrane<br>CC: GO:0031224 \| intrinsic component of membrane<br>CC: GO:0044425 \| membrane part<br>CC: GO:0016020 \| membrane | 0.0279921560341479<br>0.0279921560341479<br>0.0421000042900072<br>0.0287272290997126 | 0.13996078017074<br>0.13996078017074<br>0.210500021450036<br>0.143636145498563 | CC: GO:0016021 \| integral component of membrane<br>CC: GO:0005575 \| cellular_component | Downregulated in sigF mutant implicated in response to oxidative stress |

### Secreted proteins

| | | | | | GOA based Gene Annotation | | | BLAST2GO based GO annotation | |
|---|---|---|---|---|---|---|---|---|---|
| **Locus tag (gene name)** | **Gene Annotation** | **Location** | **Strand** | **Length** | **GO category: GO ID \| GO terms** | **Uncorrected p-value** | **Bonferroni Corrected p-value** | **GO category: GO ID \| GO terms** | **Literature based Annotation** |
| MSMEG_2112 | secreted protein; sequence based: might have Transmembrane helix | 2189268-2189852 | + | 585 | CC: GO:0016021 \| integral component of membrane<br>CC: GO:0031224 \| intrinsic component of membrane<br>CC: GO:0044425 \| membrane part<br>CC: GO:0016020 \| membrane | 0.0279921560341479<br>0.0279921560341479<br>0.0421000042900072<br>0.0287272290997126 | 0.13996078017074<br>0.13996078017074<br>0.210500021450036<br>0.143636145498563 | CC: GO:0005575 \| cellular_component | Downregulated in sigF mutant implicated in response to oxidative stress |
| MSMEG_5617 | immunogenic protein MPT63<br>MPT63 is a secreted protein of unknown function that is specific to Mycobacterium tuberculosis having IG-like fold. putative role includes involvement in host-bacterial interactions involved in endocytosis or phagocytosis, possibly during bacterial internalisation although function is unknown | 5700826-5701302 | - | 477 | NA | NA | NA | CC: GO:0005615 \| extracellular space | Downregulated in sigF mutant implicated in response to oxidative stress |
| MSMEG_2958 | | 3020304-3021023 | + | 720 | CC: GO:0016021 \| integral component of membrane<br>CC: GO:0031224 \| intrinsic component of membrane<br>CC: GO:0044425 \| membrane part<br>CC: GO:0016020 \| membrane | 0.0279921560341479<br>0.0279921560341479<br>0.0421000042900072<br>0.0287272290997126 | 0.13996078017074<br>0.13996078017074<br>0.210500021450036<br>0.143636145498563 | NA | NA |

### Transcriptional regulator

| | | | | | GOA based Gene Annotation | | | BLAST2GO based GO annotation | |
|---|---|---|---|---|---|---|---|---|---|
| **Locus tag (gene name)** | **Gene Annotation** | **Location** | **Strand** | **Length** | **GO category: GO ID \| GO terms** | **Uncorrected p-value** | **Bonferroni Corrected p-value** | **GO category: GO ID \| GO terms** | **Literature based Annotation** |
| MSMEG_1802 | ChaB protein (cation transport regulator); pfam domain: Rho termination factor, N-terminal domain (PF07498) present | 1880724-1881143 | + | 420 | BP: GO:0006807 \| nitrogen compound metabolic process<br>BP: GO:0043170 \| macromolecule metabolic process<br>BP: GO:0071704 \| organic substance metabolic process<br>BP: GO:1901576 \| organic substance biosynthetic process<br>BP: GO:0044238 \| primary metabolic process<br>BP: GO:0044249 \| cellular biosynthetic process<br>BP: GO:0009058 \| biosynthetic process<br>BP: GO:0044237 \| cellular metabolic process<br>BP: GO:0008152 \| metabolic process<br>BP: GO:0009987 \| cellular process | 0.988878773902595<br>0.890726079425172<br>0.969155736563623<br>0.914545046870011<br>0.976923093638226<br>0.895349081680126<br>0.929335307087835<br>0.954670564139802<br>0.990623690614229<br>0.979738654330216 | 1.00<br>1.00<br>1.00<br>1.00<br>1.00<br>1.00<br>1.00<br>1.00<br>1.00<br>1.00 | BP: GO:0034641 \| cellular nitrogen compound metabolic process<br>BP: GO:0006353 \| DNA-templated transcription, termination<br>BP: GO:0009058 \| biosynthetic process | 1. positive regulator of ChaA which is a Na/K/ antiporter in e.coli, cellular nitrogen compound metabolic process<br>2. Downregulated in sigF mutant implicated in response to oxidative stress |

### Hypothetical

| | | | | | GOA based Gene Annotation | | | BLAST2GO based GO annotation | |
|---|---|---|---|---|---|---|---|---|---|
| **Locus tag (gene name)** | **Gene Annotation** | **Location** | **Strand** | **Length** | **GO category: GO ID \| GO terms** | **Uncorrected p-value** | **Bonferroni Corrected p-value** | **GO category: GO ID \| GO terms** | **Literature based Annotation** |
| MSMEG_1793 | membrane protein | 1873863-1874426 | - | 564 | CC: GO:0016021 \| integral component of membrane<br>CC: GO:0031224 \| intrinsic component of membrane<br>CC: GO:0044425 \| membrane part<br>CC: GO:0016020 \| membrane | 0.0279921560341479<br>0.0279921560341479<br>0.0421000042900072<br>0.0287272290997126 | 0.13996078017074<br>0.13996078017074<br>0.210500021450036<br>0.143636145498563 | NA | NA |
| MSMEG_1774 | conserved hypothetical protein | 1861547-1862041 | - | 495 | CC: GO:0016021 \| integral component of membrane<br>CC: GO:0031224 \| intrinsic component of membrane<br>CC: GO:0044425 \| membrane part<br>CC: GO:0016020 \| membrane | 0.0279921560341479<br>0.0279921560341479<br>0.0421000042900072<br>0.0287272290997126 | 0.13996078017074<br>0.13996078017074<br>0.210500021450036<br>0.143636145498563 | NA | 1. DUF4383 domain containing protein<br>2. Downregulated in sigF mutant implicated in response to oxidative stress |
| MSMEG_5543 | hypothetical protein | 5632985-5633326 | - | 342 | NA | NA | NA | NA | Downregulated in sigF mutant implicated in response to oxidative stress |
| MSMEG_1076 | conserved hypothetical protein | 1141429-1141704 | + | 276 | NA | NA | NA | NA | Downregulated in sigF mutant implicated in response to oxidative stress |
| MSMEG_3419 | hypothetical protein | 3489877-3490839 | - | 963 | NA | NA | NA | NA | Downregulated in sigF mutant implicated in response to oxidative stress |
| MSMEG_3439 | hypothetical protein | 3508893-3509048 | + | 156 | CC: GO:0016021 \| integral component of membrane<br>CC: GO:0031224 \| intrinsic component of membrane<br>CC: GO:0044425 \| membrane part<br>CC: GO:0016020 \| membrane | 0.0279921560341479<br>0.0279921560341479<br>0.0421000042900072<br>0.0287272290997126 | 0.13996078017074<br>0.13996078017074<br>0.210500021450036<br>0.143636145498563 | NA | Downregulated in sigF mutant implicated in response to oxidative stress |
| MSMEG_3438 | hypothetical protein | 3508887-3509057 | - | 171 | NA | NA | NA | NA | NA |
| MSMEG_3418 | conserved hypothetical protein | 3489356-3489784 | - | 429 | NA | NA | NA | NA | Downregulated in sigF mutant implicated in response to oxidative stress |
| MSMEG_5343 | conserved hypothetical protein | 5427543-5427890 | - | 348 | NA | NA | NA | NA | Downregulated in sigF mutant implicated in response to oxidative stress |
| MSMEG_1789 | conserved hypothetical protein | 1871520-1871675 | + | 156 | CC: GO:0016021 \| integral component of membrane<br>CC: GO:0031224 \| intrinsic component of membrane<br>CC: GO:0044425 \| membrane part<br>CC: GO:0016020 \| membrane | 0.0279921560341479<br>0.0279921560341479<br>0.0421000042900072<br>0.0287272290997126 | 0.13996078017074<br>0.13996078017074<br>0.210500021450036<br>0.143636145498563 | NA | Downregulated in sigF mutant implicated in response to oxidative stress |
| MSMEG_6667 | conserved hypothetical protein | 6720444-6720974 | - | 531 | CC: GO:0016021 \| integral component of membrane<br>CC: GO:0031224 \| intrinsic component of membrane<br>CC: GO:0044425 \| membrane part<br>CC: GO:0016020 \| membrane | 0.0279921560341479<br>0.0279921560341479<br>0.0421000042900072<br>0.0287272290997126 | 0.13996078017074<br>0.13996078017074<br>0.210500021450036<br>0.143636145498563 | NA | Downregulated in sigF mutant implicated in response to oxidative stress |
| MSMEG_1950 | conserved hypothetical protein | 2027837-2028406 | + | 570 | NA | NA | NA | NA | Downregulated in sigF mutant implicated in response to oxidative stress |

| | | | | | | | | | |
|---|---|---|---|---|---|---|---|---|---|
| MSMEG_6612 | ATPase, MoxR family protein | 6664363-6665364 | - | 1002 | MF: GO:0016787 \| hydrolase activity<br>MF: GO:0043168 \| anion binding<br>MF: GO:0036094 \| small molecule binding<br>MF: GO:1901265 \| nucleoside phosphate binding<br>MF: GO:0000166 \| nucleotide binding<br>MF: GO:0043167 \| ion binding<br>MF: GO:0003824 \| catalytic activity<br>MF: GO:0005488 \| binding<br>MF: GO:0097159 \| organic cyclic compound binding<br>MF: GO:1901363 \| heterocyclic compound binding | 0.548264937793926<br>0.839250716415205<br>0.895685477437695<br>0.939734266779146<br>0.939734266779146<br>0.950043978332138<br>0.977353128529576<br>0.989284577095953<br>0.994381822820708<br>0.994381822820708 | 1.00<br>1.00<br>1.00<br>1.00<br>1.00<br>1.00<br>1.00<br>1.00<br>1.00<br>1.00 | MF: GO:0016887 \| ATPase activity<br>MF: GO:0043167 \| ion binding | Downregulated in sigF mutant implicated in response to oxidative stress |
| MSMEG_3685 | conserved hypothetical protein | 3750230-3750661 | + | 432 | NA | NA | NA | NA | putative transmembrane protein |
| MSMEG_2376 | conserved hypothetical protein | 2458418-2458879 | - | 462 | NA | NA | NA | NA | 1. transmembrane protein, one helicobacter example is a Li/Na/K antiporter<br>2. Downregulated in sigF mutant implicated in response to oxidative stress |
| MSMEG_6211 | hypothetical protein | 6276404-6276556 | - | 153 | NA | NA | NA | NA | Downregulated in sigF mutant implicated in response to oxidative stress |

*Table S5: Plasmids and strains used*

| ***Episomal overexpression in M.smegmatis $MC^2$ 155 strain*** | | | |
|---|---|---|---|
| **Strain** | **Plasmid used** | **Base Plasmid** | **Details of construct** |
| SSA009 | pSSA008 | pMV261 | P*hsp60* -3Xflag-(RV1626D65E residues 4-205) |
| SSA004 | pMV261 | pMV261 | EMPTY VECTOR |
| AK001 | pSS005 | pMV261 | P*hsp60*-3Xflag-(EUT V residues 2-190 ) |
| AK002 | pSS004 | pMV261 | P*hsp60*-3Xflag-(EUT VD54E residues 2-190 ) |

| ***Integrated strains in M. smegmatis $MC^2$ 155 strain*** | | | |
|---|---|---|---|
| AK003 | pAK003 | pMV761 | P*groel*- 3Xflag-(RV1626D65E residues 4-205) |

| ***Protein expression in E. coli BL21 (DE3) strain*** | | | | | |
|---|---|---|---|---|---|
| | **Plasmid name** | **Base plasmid** | **Construct** | **Primer Name** | **Primer sequence** |
| | pHP002 | p*hisII* | 6his-(Rv3220c residues 2-501) | AR307 | 5'CATGCCATGGGTTCAACAGGAGGTAGCCAACC |
| | | | | HP001b | 5'CCCGAATTCTTACGCCGCCGCATGCGC |
| | pAR020 | p*hisII* | 6his-(Rv1626 residues 4-205) | AR301 | 5'CCGCTCGAGTTAGGTGTCTTTGGGTGTTCC |
| | | | | AR302 | 5'CATGCCATGGCGACACTCGGTGATCTGCTCG |
| | pTR001 | p*hisII* | 6his-(Rv1626D65E residues 4-205) | TR001 | 5'GACCTGGTGATCATGGAAGTGAAGATGCCGCGCC |
| | | | | TR002 | 5'GGCGCGGCATCTTCACTTCCATGATCACCAGGTC |
| | pAR006 | pET28a | 6his-(EutVD54E residues 2-190) | AR052 | 5'CCTGACCTAGTATTAATGGAAATTCAAATGCCAATTTTAG |
| | | | | AR053 | 5'CTAAAATTGGCATTTGAATTTCCATTAATACTAGGTCAGG |

| ***quantitative RT-PCR*** | | |
|---|---|---|
| **Gene name** | **Primer Name** | **Primer Sequence** |
| MSMEG_0267 | AR664 | AGTCGCTCTACTCCGATTCC |
| | AR665 | AGCGGGAAATCGTGCACCT |
| MSMEG_5575 | AR668 | TTGGAGGACATCCGCATGTG |
| | AR669 | TGGTTCAACAGCCAACGGG |
| MSMEG_1793 | AR656 | CTGTTGCTGGGGTTCGCGAT |
| | AR657 | GATCCAACAGCCCACGAAC |
| MSMEG_1076 | AR660 | AAGATGCTGAAGGACGAGAAG |
| | AR661 | GCGGCGCTTCCTTGATCTT |
| MSMEG_0267 | AR662 | GCACCCGCCATGTCACGT |
| | AR663 | TTGCTGCCGCTGGATACCC |
| MSMEG_1680 | AR652 | TCCCAACTCCTGAACGGCTA |
| | AR653 | TCCTCCACCACGATCATTCC |
| SigA MSMEG_2758 | AR678 | GAAGACACCGACCTGGAACT |
| | AR679 | GACTCTTCCTCGTCCCACAC |

| ***Templates for EMSA/ Inline Probing*** | | |
|---|---|---|
| **Accession #/coordinates** | **Name of gene/ sequence** | **DNA sequence** |
| | T7 Promoter- | TAATACGACTCACTATAGG |
| M gilvum NC_009338.1/1209109-1209187 | Esterase | T7_CGAAATGAG**AGATCACCACAAAGGTGTGGTTGACTACAGGGCAATTGGGCCGGACTCGGGG**ATCGCTTGGGAAAAGC |
| M.tuberculosis H37Rv NC_000962.3/2187263-2187311 | RV1936/ mono-oxygenase | T7_AAATTCCCATGCAAAGAAGCACTTGTGTACAACGAAGTATCTTGGTAGT |
| M.tuberculosis H37Rv NC_000962.3/4340162-4340234 | EspE | T7_CAAAAAATTCGTGCACCAACCCCCCTCCGAGCGCTGCTAAGCTCAATGTGCAGTGCAAAGGTGCAGATAATGATG |
| | random RNA | T7_TTACGCTTGAATTAGCGTCCTCACCTGCTTCAGCCGTCTCCAGGTAATCAGCCCCGGCATCAGTGCCGTCGCCAATAAAATCTAGGTGTTGTCGGATAAAATCCAAAT |
| M.smegmatis NC_008596.1/3682246-3682309 | MSMEG_3619 | T7_GTGATCCCCGGTACATGGAGGTACATCATGGCGGAGGTGCTGGTCACCGGCGGCGACACCGAAC |

| ***In vitro transcription and translation coupled assay*** | | | |
|---|---|---|---|
| | | **Primer Names** | **DNA sequence** |
| M.smegmatis NC_008596.11/3682273-3682944 | P*T7*- (5'UTR up to residue 222 of MSMEG_3619)-3Xflag | AK188 | 5' TAATACGACTCACTATAGGGGTGATCCCCGGTACATGG |
| | | AK192 | 5' TCACTTGTCGTCGTCGTCCTTGTAGTCGATGTCGTGGTCCTTGTAGTCGCCGTCGTGGTCCTTGTAGTCGACGGTTGCGAGCGCACC |
